# Genome-wide association study of susceptibility to idiopathic pulmonary fibrosis

**DOI:** 10.1101/636761

**Authors:** Richard J Allen, Beatriz Guillen-Guio, Justin M Oldham, Shwu-Fan Ma, Amy Dressen, Megan L Paynton, Luke M Kraven, Ma’en Obeidat, Xuan Li, Michael Ng, Rebecca Braybrooke, Maria Molina-Molina, Brian D Hobbs, Rachel K Putman, Phuwanat Sakornsakolpat, Helen L Booth, William A Fahy, Simon P Hart, Mike R Hill, Nik Hirani, Richard B Hubbard, Robin J McAnulty, Ann B Millar, Vidyia Navaratnam, Eunice Oballa, Helen Parfrey, Gauri Saini, Moira K B Whyte, Gunnar Gudmundsson, Vilmundur Gudnason, Hiroto Hatabu, David J Lederer, Ani Manichaikul, John D Newell, George T O’Connor, Victor E Ortega, Hanfei Xu, Tasha E Fingerlin, Yohan Bossé, Ke Hao, Philippe Joubert, David C Nickle, Don D Sin, Wim Timens, Dominic Furniss, Andrew P Morris, Krina Zondervan, Ian P Hall, Ian Sayers, Martin D Tobin, Toby M Maher, Michael H Cho, Gary M Hunninghake, David A Schwartz, Brian L Yaspan, Philip L Molyneaux, Carlos Flores, Imre Noth, R Gisli Jenkins, Louise V Wain

## Abstract

**Rationale:** Idiopathic pulmonary fibrosis (IPF) is a complex lung disease characterised by scarring of the lung that is believed to result from an atypical response to injury of the epithelium. The mechanisms by which this arises are poorly understood and it is likely that multiple pathways are involved. The strongest genetic association with IPF is a variant in the promoter of *MUC5B* where each copy of the risk allele confers a five-fold risk of disease. However, genome-wide association studies have reported additional signals of association implicating multiple pathways including host defence, telomere maintenance, signalling and cell-cell adhesion.

**Objectives:** To improve our understanding of mechanisms that increase IPF susceptibility by identifying previously unreported genetic associations.

**Methods and measurements:** We performed the largest genome-wide association study undertaken for IPF susceptibility with a discovery stage comprising up to 2,668 IPF cases and 8,591 controls with replication in an additional 1,467 IPF cases and 11,874 controls. Polygenic risk scores were used to assess the collective effect of variants not reported as associated with IPF.

**Main results:** We identified and replicated three new genome-wide significant (*P*<5×10^-8^) signals of association with IPF susceptibility (near *KIF15, MAD1L1* and *DEPTOR)* and confirm associations at 11 previously reported loci. Polygenic risk score analyses showed that the combined effect of many thousands of as-yet unreported IPF risk variants contribute to IPF susceptibility.

**Conclusions:** Novel association signals support the importance of mTOR signalling in lung fibrosis and suggest a possible role of mitotic spindle-assembly genes in IPF susceptibility.

## Introduction

Idiopathic pulmonary fibrosis (IPF) is a devastating lung disease characterised by the build-up of scar tissue. It is believed that damage to the alveolar epithelium is followed by an aberrant wound healing response leading to the deposition of dense fibrotic tissue, reducing the lungs’ flexibility and inhibiting gas transfer^1^. Treatment options are limited and half of individuals diagnosed with IPF die within 3 years^1,2^. Two drugs (pirfenidone and nintedanib) have been approved for the treatment of IPF, but neither offer a cure and only slow disease progression.

IPF is associated with a number of environmental and genetic factors. Identifying regions of the genome contributing to disease risk improves our understanding of the biological processes underlying IPF and helps in the development of new treatments^3^. To date, genome-wide association studies^4–8^ (GWAS) have reported 17 common variant (minor allele frequency [MAF]>5%) signals associated with IPF; stressing the importance of host defence, telomere maintenance, cell-cell adhesion and signalling with respect to disease susceptibility. The sentinel (most strongly associated) variant, rs35705950, in one of these signals that maps to the promoter region of the *MUC5B* gene, has a much larger effect on disease susceptibility than other reported risk variants with each copy of the risk allele associated with a five-fold increase in odds of disease^9^. Despite this, the variant rs35705950 has a risk allele frequency of 35% in cases (compared to 11% in the general population) and so does not explain all IPF risk. Rare variants (MAF<1%) in telomere-related and surfactant genes have also been implicated in familial pulmonary fibrosis and sporadic IPF^10,11^.

In this study, we performed the largest GWAS of IPF susceptibility to date, followed by bioinformatics analysis of gene expression data to identify the genes underlying identified association signals and inform our understanding of IPF pathogenesis and risk. As specific IPF associated variants have also been shown to overlap with other respiratory traits including lung function, chronic obstructive pulmonary disease (COPD)^12,13^ and interstitial lung abnormalities^14^ (ILAs, which might be a precursor lesion for IPF), we tested for association of the IPF risk variants with other respiratory phenotypes in independent datasets. Finally, using polygenic risk scores, we tested whether there was a still substantial contribution to IPF risk from genetic variants with as-yet unconfirmed associations with IPF susceptibility.

## Methods

### Study design

We analysed genome-wide association data from three independent IPF case-control collections. These three studies (named here as the Chicago^5^, Colorado^6^ and UK^8^ studies) are previously described and comprise of patients with IPF and non-IPF controls (**Appendix**). Two more case-control collections (named here as the UUS [USA, UK and Spain] and Genentech studies) that had not contributed to any previous IPF GWAS were included as a replication dataset. In the UUS study, cases were recruited from across the USA, UK and Spain and controls were selected from UK Biobank to follow a similar sex and smoking distribution seen in the IPF cases (**Appendix**). The Genentech study consisted of cases from three IPF clinical trials and controls from four non-IPF clinical trials that have been previously described (Appendix)^15^. All five studies restricted to unrelated individuals of European ancestry and we applied stringent quality control measures (**Appendix**). All studies diagnosed IPF cases using American Thoracic Society and European Respiratory Society guidelines^16–18^ and had appropriate institutional review board or ethics approval.

### Procedures and statistical analysis

For this study, genotype data for the Colorado, Chicago, UK and UUS were imputed separately using the Haplotype Reference Consortium (HRC) r1.1 panel^19^ (**Appendix**). Whole-genome sequencing data was available for the Genentech study. Overlap of cases and controls between studies was assessed using KING^20^ v2.1.2 (**Appendix**) to remove duplicate samples.

#### Identification of IPF susceptibility signals

In each of the Chicago, Colorado and UK studies separately, a genome-wide analysis of IPF susceptibility, using SNPTEST^21^ v2.5.2, was conducted assuming an additive genetic effect and adjusting for the first 10 principal components to account for fine-scale population structure. Only bi-allelic autosomal variants that had a minor allele count ≥10 within the study, were in Hardy-Weinberg Equilibrium (*P*>10^-6^) and were well imputed (imputation quality R^2^>0.5) were included.

For identification of genetic signals associated with IPF susceptibility, a fixed effect inverse-variance weighted genome-wide meta-analysis (**Figure 1**) of the IPF susceptibility association summary statistics was performed across the Chicago, Colorado and UK studies using R v3.5.1 (discovery stage). Results were corrected for inflation due to residual fine-scale population structure using genomic control at both the study and meta-analysis level. Only genetic variants represented in at least two studies were included in the discovery analysis. Conditional analyses were performed to identify independent association signals in each locus (**Appendix**).

**Figure 1 -.**
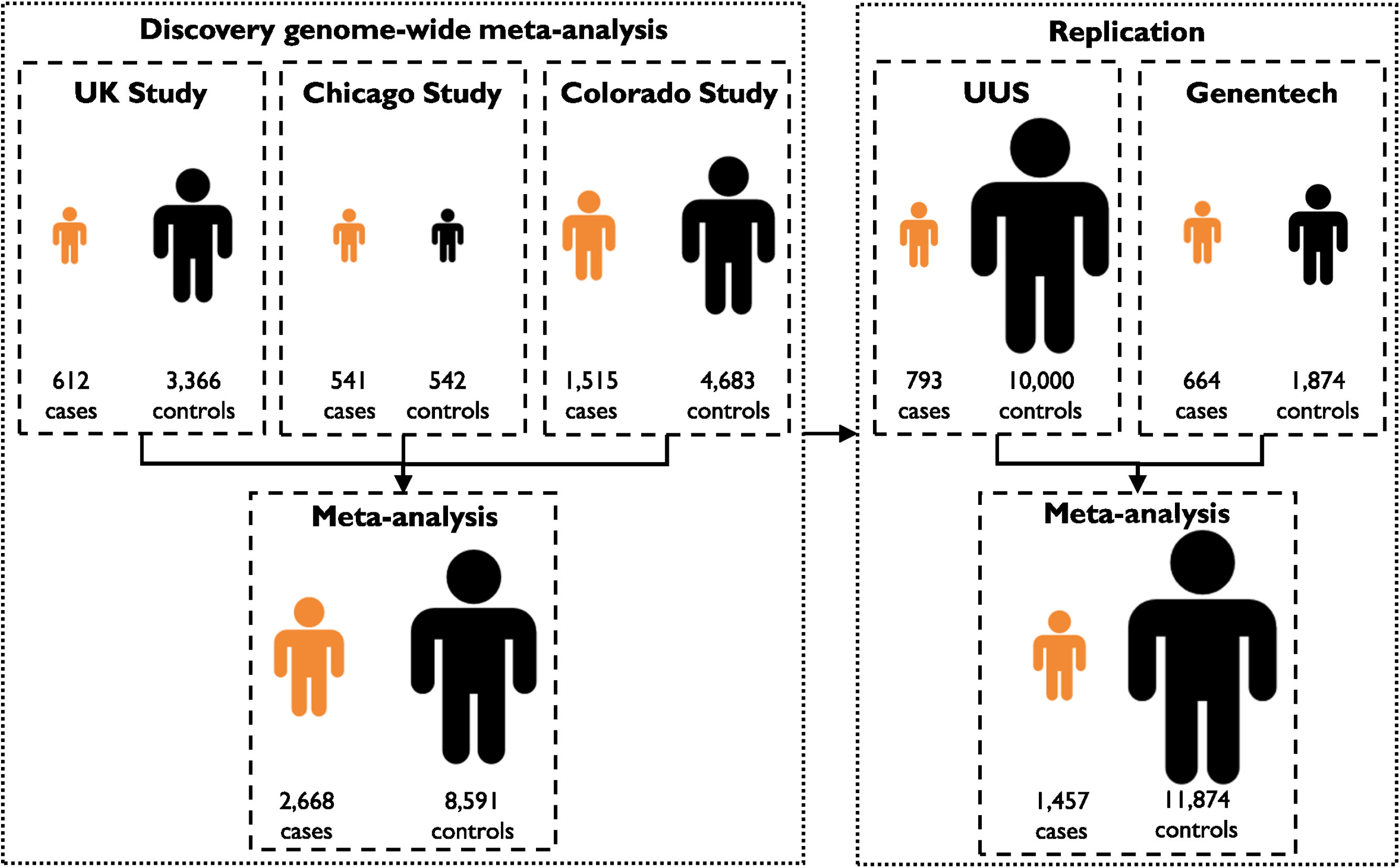
Sample sizes for genome-wide analyses

Novel signals reaching genome-wide significance and showing nominal significance (*P*<0.05) with consistent direction of effect in each contributing study were further tested in the replication samples. We considered novel signals to be associated with IPF risk if they reached a Bonferroni-corrected threshold (*P*<0.05 / number of signals followed-up) in a replication meta-analysis of the UUS and Genentech studies (replication stage, **Appendix**). Previously reported signals reaching genome-wide significance (*P*<5×10^-8^) in the discovery meta-analysis were deemed as showing a confirmed association with IPF risk.

#### Characterisation of signals and functional follow-up

For each association signal the posterior probability of each variant in the region being causal was calculated (assuming there is one causal variant and it has been measured)^22^. From this, a 95% credible set (i.e. the smallest set of variants that is 95% likely to contain the causal variant) was generated for each IPF signal (**Appendix**). VEP (Variant Effect Predictor)^23^ was used to annotate each credible set variant to identify deleterious variants (**Appendix**).

Linked genotype and gene expression data resources were interrogated to identify putative causal genes for the novel association signals. Variants in the 95% credible sets were investigated in three eQTL databases; a lung eQTL database consisting of individuals from three cohorts (Universities of British Columbia, Laval and Groningen, n=1,111)^24–26^, the NESDA-NTR (Netherlands Study of Depression and Anxiety and the Netherlands Twin Register) blood eQTL database (n=4,896)^27^ and 48 tissue types in GTEx (Genotype-Tissue Expression project, n between 80 and 491)^28^. An FDR threshold of 10% was used for the lung eQTL database and NESDA-NTR, and an FDR threshold of 5% was used for the smaller GTEx resource. Where a variant in a credible set for a novel association signal was found to be an eQTL for a gene, we calculated the posterior probability of the IPF GWAS signal and eQTL signal being driven by the same variant (colocalisation) using coloc^29^. Colocalisation between IPF susceptibility and eQTL signal was defined as when the posterior probability of the IPF GWAS and eQTL signals being driven by the same variant was greater than 80% (**Appendix**). Only genes where there was colocalisation of the IPF susceptibility and eQTL signal are reported.

The variants with the highest posterior probability of causality for the novel signals were explored using HaploReg v4.1^30^ to identify overlap with histone mark promoters and enhancers in relevant tissue (i.e. lung). We used DeepSEA^31^, a deep-learning variant effect predictor, to identify whether any of the IPF risk signals were predicted to have a functional effect on chromatin features and transcription factor binding sites. The IPF risk signals were tested for enrichment in regulatory regions using FORGE^32^ and GARFIELD^33^. SNPsea^34^ was used to assess enrichment of genes in linkage disequilibrium with IPF risk variants in i) 1,751 genetic pathways and, ii) in genes showing differential expression between IPF cases and controls in four epithelial cell types^35^ (Appendix).

#### Association with other respiratory traits

The variant with the highest posterior probability of causality in the credible set for each IPF risk signal was investigated for its association with measures of lung function^36^ and interstitial lung abnormalities (**Appendix**). Lung function (namely FEV_1_, FVC, the ratio FEV_1_/FVC and PEF) was tested in 400,102 individuals^36^. ILA analyses were performed comparing up to 1,699 individuals with any ILA to 10,274 individuals without any ILA; an additional analysis restricted the ILA cases to 1,287 individuals with subpleural ILAs. Variants were reported as associated with lung function or ILA if they met a Bonferroni corrected *P* value threshold for the number of variants and traits investigated.

#### Polygenic risk scores

Polygenic risk scores were utilised to assess the contribution of as-yet unreported variants to IPF risk. Polygenic risk scores allow for the cumulative effect of many genetic variants to be studied. The polygenic risk score was equal to the number of risk alleles carried multiplied by the effect size of the variant, summed across all variants included in the score, i.e.:

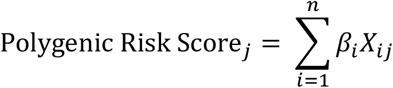

where *β_i_* is the log(OR) of variant *i* from the genome-wide meta-analysis of the UK, Chicago and Colorado studies, *X_ij_* is the genotype of variant *i* for person *j* and *n* is the number of variants. Scores were generated for individuals in the independent UUS study using independent variants selected after LD-clumping (r^2^≤0.1). This score was tested to identify whether it was associated with IPF susceptibility, adjusting for 10 principal components to account for fine-scale population structure, using PRSice v1.25^37^. As we wanted to explore the contribution to IPF risk from variants not yet reported, we excluded variants within 1Mb of each IPF risk locus identified in this IPF susceptibility GWAS. We altered the number of variants included in the risk score calculation by setting a P-threshold (*P*_T_) criteria such that the variant had to have a *P* value<*P*_T_ in the genome-wide meta-analysis to be included in the score. Given multiple testing, we used the recommended significance threshold of *P*<0.001 for determining significantly associated risk scores^37^.

## Results

Following quality control, 541 cases and 542 controls from the Chicago study, 1,515 cases and 4,683 controls from the Colorado study and 612 cases and 3,366 controls from the UK study were available (**Table 1, Supplementary Figure 1**) to contribute to the discovery stage of the genome-wide susceptibility analysis. For the replication stage of the GWAS, after quality control, there were 803 cases and 10,000 controls available in the UUS study and 664 cases and 1,874 controls available in the Genentech study (**Appendix**).

**Table 1:** Demographics

|  | Chicago |  | Colorado |  | UK |  | UUS |  | Genentech |  |
| --- | --- | --- | --- | --- | --- | --- | --- | --- | --- | --- |
|  | Cases | Controls | Cases | Controls | Cases | Controls | Cases | Controls | Cases | Controls |
| **n** | 541 | 542 | 1,515 | 4,683 | 612 | 3,366 | 793 | 10,000 | 664 | 1,874 |
| **Genotyping array /sequencing** | Affymetrix 6.0 SNP array |  | Illumina Human 660W Quad BeadChip |  | Affymetrix UK BiLEVE array | Affymetrix UK BiLEVE and UK Biobank arrays | Affymetrix UK Biobank and Spain Biobank arrays | Affymetrix UK BiLEVE and UK Biobank arrays | HiSeq X Ten platform (Illumina) |  |
| **Imputation panel** | HRC |  | HRC |  | HRC |  | HRC |  | - |  |
| **Age (mean)** | 68 | 63 <sup>a</sup> | 66 | - | 70 | 65 | 69 | 58 | 68 | - |
| **Sex (% males)** | 71% | 47% | 68% | 49% | 71% | 70% | 75.2% | 72.1% | 73.5% | 27.1% |
| **% ever smokers** | 72% | 42% | - | - | 70% | 70% | 68.7% <sup>b</sup> | 68% | 67.3% | 18.1% <sup>c</sup> |

<sup>a</sup> Age only available for 103 Chicago controls
<sup>b</sup> Smoking status only recorded for 753 IPF cases in UUS
<sup>c</sup> Smoking status only recorded for 481 of the Genentech controls

To identify new signals of association, we meta-analysed the genome-wide association results for IPF susceptibility for the Chicago, Colorado and UK studies. This gave a maximum sample size of up to 2,668 cases and 8,591 controls for 10,790,934 well imputed (R^2^>0.5) variants with minor allele count ≥10 in each study and which were available in two or more of the studies (**Supplementary Figure 2**). We identified 14 IPF risk signals (11 of which have been previously reported and three were novel, **Figure 2** and **Supplementary Figure 3**). Conditional analyses did not identify any additional independent association signals at these loci.

**Figure 2 -.**
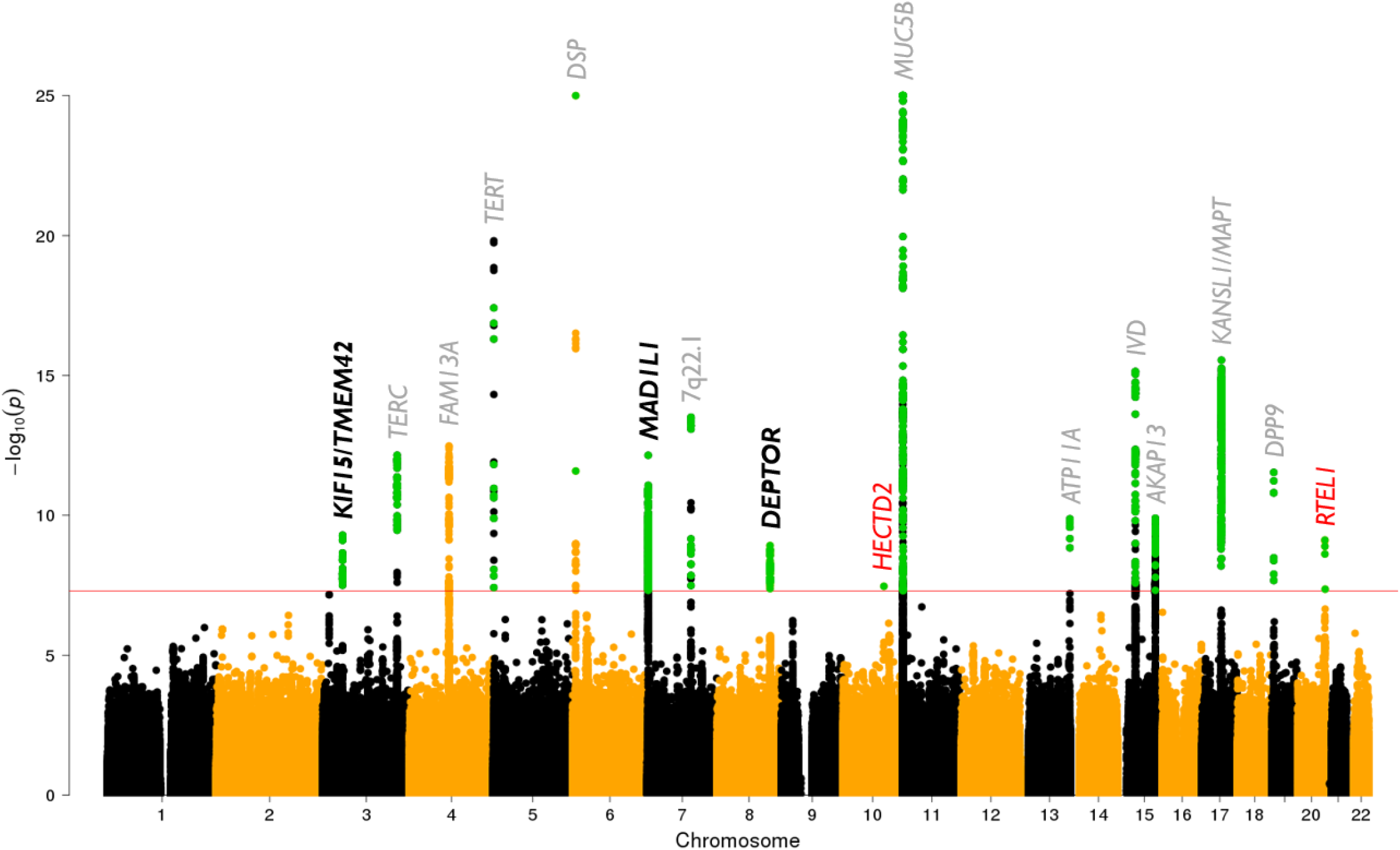
Manhattan plot of discovery analysis results X axis shows chromosomal position and the y axis shows the –log(P value) for each variant in the discovery genome-wide analysis. The red line shows genome-wide significance (*P*<5×10^-8^) and variants in green met the criteria for further study in the replication analysis (i.e. reached genome-wide significance in the discovery meta-analysis and had *P*<0.05 and consistent direction of effects in each study). Genes labelled in grey are previously reported signals that reach significance in the discovery genome-wide meta-analysis. Genes labelled in black are the novel signals identified in the discovery analysis that reach genome-wide significance when meta-analysing discovery and replication samples. The signals which did not replicate are shown by red labels. For ease of visualisation the y axis has been truncated at 25.

We identified three novel signals (in 3p21.31 [near *KIF15,* **Figure 3i**], 7p22.3 [near *MAD1L1,* **Figure 3ii**] and 8q24.12 [near *DEPTOR,* **Figure 3iii**]) that showed an association in the discovery meta-analysis and were also significant after adjusting for multiple testing (*P*<0.01) in the replication stage comprising 1,467 IPF cases and 11,874 controls. Two additional loci were genome-wide significant in the genome-wide discovery analysis but did not reach significance in the replication studies. The sentinel variants of these two signals were a low frequency intronic variant in *RTEL1* (MAF=2.1%, replication *P*=0.012) and a rare intronic variant in *HECTD2* (MAF=0.3%, replication *P*=0.155) (**Table 2, Supplementary Table 1, Supplementary Figures 5 and 6**).

**Figure 3.**
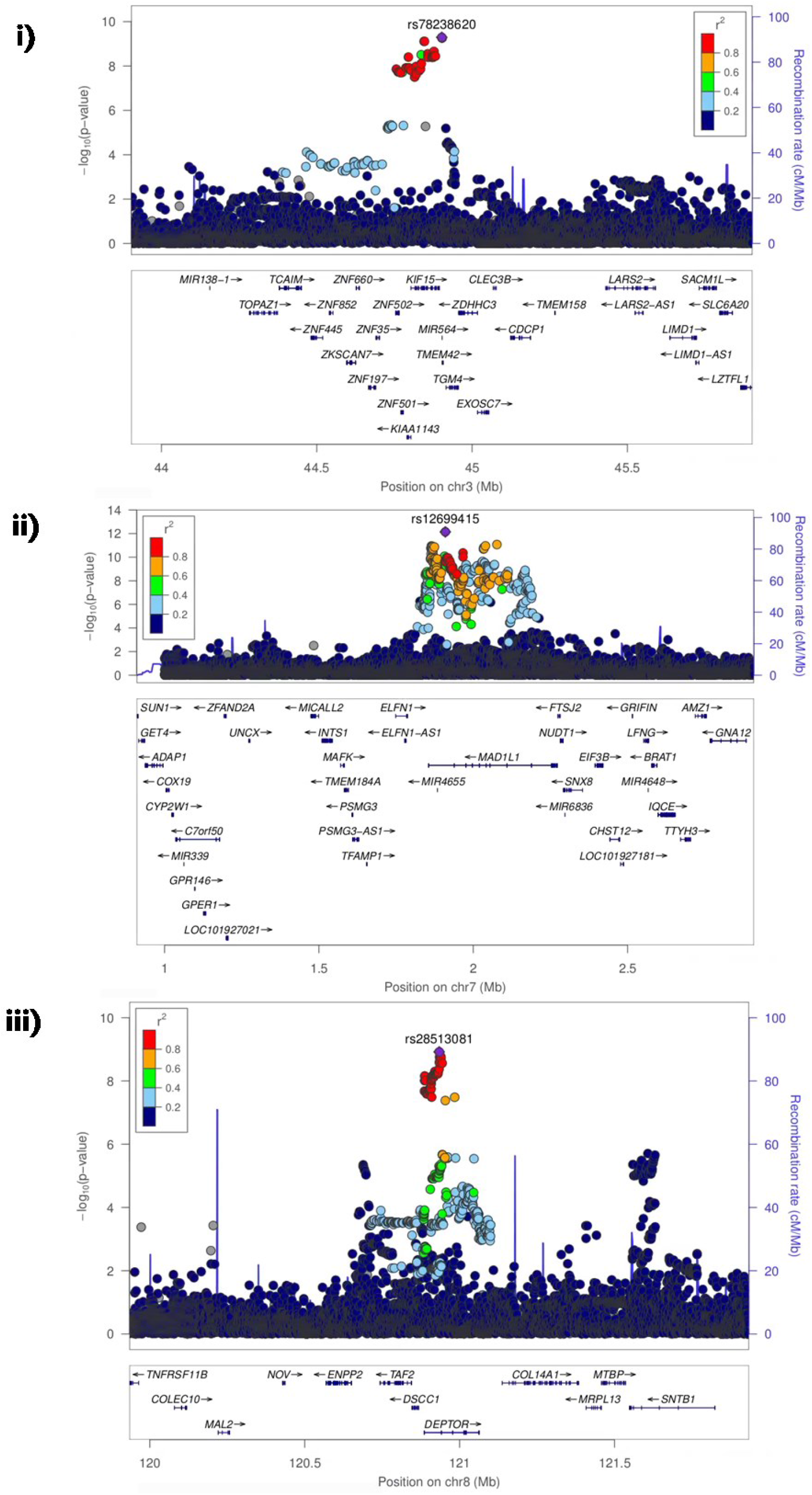
Region plots of three novel risk loci

**Table 2 -.** Main results. The minor allele is the effect allele and the minor allele frequency (MAF) is taken from across the studies used in the discovery meta-analysis.

| Chr | Pos | rsid | Locus | Major allele | Minor allele | MAF | Discovery meta-analysis |  | Replication meta-analysis |  | Meta-analysis of discovery and replication |  |
| --- | --- | --- | --- | --- | --- | --- | --- | --- | --- | --- | --- | --- |
|  |  |  |  |  |  |  | OR [95% CI] | P | OR [95% CI] | P | OR [95% CI] | P |
| 3 | 44902386 | rs78238620 | *KIF15* | T | A | 5.3% | 1.58 [1.37, 1.83] | $5.12 \times 10^{-10}$ | 1.48 [1.24, 1.77] | $1.43 \times 10^{-5}$ | 1.54 [1.38, 1.73] | $4.05 \times 10^{-14}$ |
| 7 | 1909479 | rs12699415 | *MAD1L1* | G | A | 42.0% | 1.28 [1.19, 1.37] | $7.15 \times 10^{-13}$ | 1.29 [1.18, 1.41] | $2.27 \times 10^{-8}$ | 1.28 [1.21, 1.35] | $9.38 \times 10^{-20}$ |
| 8 | 120934126 | rs28513081 | *DEPTOR* | A | G | 42.8% | 0.82 [0.76, 0.87] | $1.20 \times 10^{-9}$ | 0.87 [0.80, 0.95] | 0.002 | 0.83 [0.79, 0.88] | $1.93 \times 10^{-11}$ |
| 10 | 93271016 | rs537322302 | *HECTD2* | C | G | 0.3% | 7.82 [3.77, 16.2] | $3.43 \times 10^{-8}$ | 1.75 [0.81, 3.78] | 0.155 | 3.85 [2.27, 6.54] | $6.25 \times 10^{-7}$ |
| 20 | 62324391 | rs41308092 | *RTEL1* | G | A | 2.1% | 2.12 [1.67, 2.69] | $7.65 \times 10^{-10}$ | 1.45 [1.08, 1.94] | 0.012 | 1.82 [1.51, 2.19] | $2.24 \times 10^{-10}$ |

For the novel signal on chromosome 3, the sentinel variant (rs78238620) was a low frequency variant (MAF=5%) in an intron of *KIF15* with the minor allele being associated with increased susceptibility to IPF. The IPF susceptibility signal was associated with decreased expression of *KIF15* in brain tissue and the nearby gene *TMEM42* in thyroid (**Supplementary Figure 7, Supplementary Tables 2 and 3i**).

In the statistical fine-mapping of the novel signal on chromosome 7, the sentinel variant (rs12699415, in an intron of *MAD1L1)* had a much higher posterior probability of causality for IPF (36%) compared to the other variants in the credible set for that signal (all others had probability<4%). The IPF susceptibility signal was associated with decreased expression of *MAD1L1* in heart tissue (**Supplementary Figure 8, Supplementary Tables 2 and 3ii**).

For the signal on chromosome 8, the sentinel variant (rs28513081) was located in an intron of *DEPTOR.* The fine-mapping analysis identified 95 variants in the 95% credible set (all located in introns of *DEPTOR*) each contributing <5% posterior probability of causality. The IPF risk allele was associated with decreased expression of *DEPTOR* in colon, lung (in both the lung eQTL database and GTEx) and skin. The allele was also associated with increased expression of *DEPTOR* in the NESDA-NTR database (whole blood), increased expression of *TAF2* (in colon), RP11-760H22.2 (increased in adipose, and decreased in colon, and lung) and increased expression of KB-1471A8.1 (in adipose and skin, **Supplementary Figure 9, Supplementary Tables 2 and 3iii**).

We confirmed genome-wide significant associations with IPF susceptibility for 11 of the 17 previously reported signals (in or near *TERC, TERT, DSP,* 7q22.1, *MUC5B, ATP11A, IVD, AKAP13, KANSL1, FAM13A* and *DPP9;* **Supplementary Table 1, Supplementary Figure 4**). Three further signals at 11p15.5 (near *MUC5B*) were no longer genome-wide significant after conditioning on the *MUC5B* promoter variant (**Supplementary Table 1**), consistent with previous reports^6^.

Of the 14 IPF risk signals (i.e. the 11 previously reported signals we confirmed and three novel signals), the only variant predicted to have a functional effect using DeepSEA was rs2013701 (in an intron of *FAM13A),* which was associated with a change in DNase in 18 cell types and FOXA1 in the T-47D cell line (a breast cancer cell line derived from a pleural effusion). The 14 IPF risk signals were found to be enriched in DNase I hypersensitivity site regions in foetal lung tissue (**Supplementary Figure 10**). Taking all variants with genome-wide significant (*P*<5×10^-8^) association with IPF susceptibility from our discovery analysis, we found enrichment in DNase I hypersensitivity sites in multiple tissues (including foetal lung) (**Supplementary Figure 11**). No pathway-specific geneexpression enrichment or enrichment in differential expression in airway epithelial cells between IPF cases and healthy controls was observed for the 14 IPF risk signals when using SNPsea (**Supplementary Table 4**).

The sentinel variants of 12 of the 14 IPF risk loci were at least nominally associated (*P*<0.05) with one or more lung function trait in general population studies (**Supplementary Table 5**). After adjustments for multiple testing (*P*<5.2×10^-4^), the previously reported variants at *FAM13A, DSP* and *IVD* were associated with decreased FVC and variants at *FAM13A, DSP,* 7q22.1 *(ZKSCAN1)* and *ATP11A* were associated with increased FEV1/FVC. Similarly, for the three novel risk variants, all showed at least a nominal association with decreased FVC and increased FEV1/FVC. We observed a nominally significant association of the *MUC5B* IPF risk allele with decreased FVC and increased FEV1/FVC. The IPF risk alleles at *MAPT* were significantly associated with both increased FEV1 and FVC. Eight of the IPF risk loci were at least nominally significantly associated with either ILA or subpleural ILA with consistent direction of effects (i.e. the allele associated with increased IPF risk was also associated with increased ILA risk). The new *KIF15, MAD1L1* and *DEPTOR* signals were not associated with ILA (although the rare risk allele at *HECTD2* that did not replicate in our study showed some association with an increased risk of subpleural ILA [*P*=0.003] with a large effect size similar to that observed in the IPF discovery meta-analysis).

To quantify the impact of as-yet unreported variants on IPF susceptibility, polygenic risk scores were calculated excluding the 14 IPF risk variants (as well as all variants within 1Mb). The polygenic risk scores were significantly associated with increased IPF risk despite exclusion of the known genetic association signals (including *MUC5B*). The most significant risk score was observed when including variants from the genome-wide discovery meta-analysis with *P*<0.664 (risk score *P*=1.41×10^-24^, **Supplementary Figure 12**). This risk score contained 806,476 independent variants and explained approximately 2% of the phenotypic variation (Nagelkerke’s R^2^=0.023). These results suggest that there is a significant contribution of additional as-yet undetected common and low frequency variants to IPF susceptibility.

## Discussion

We undertook the largest GWAS of IPF susceptibility to date and identified three novel signals of association that implicated genes not previously known to be important in IPF.

The new signal on chromosome 8 implicates *DEPTOR,* which encodes the DEP Domain containing MTOR interacting protein. *DEPTOR* inhibits mTOR kinase activity as part of both the mTORC1 and mTORC2 protein complexes. The IPF susceptibility risk allele at this locus was associated with decreased gene expression of *DEPTOR* in lung tissue. TGFβ-induced DEPTOR suppression can stimulate collagen synthesis^38^ and the importance of mTORC1 signalling via 4E-BP1 for TGFβ induced collagen synthesis has recently been demonstrated in fibrogenesis^39^. The signal on chromosome 8 did also colocalise with expression of *TAF2,* RP11-760H22.2 and KB-1471A8.1 in similar tissues to *DEPTOR. MAD1L1,* implicated by a new signal on chromosome 7, is a mitotic checkpoint gene, mutations in which have been associated with multiple cancers including lung cancer^40^. Another spindle-assembly related gene, *KIF15,* was implicated by the new signal on chromosome 3 (along with *TMEM42).*

The genome-wide study also identified two signals that were not replicated after multiple testing adjustments. The intronic variant in *RTEL1* had the same direction of effect in all five studies and was nominally significant in four studies. Multiple genes of the telomere complex and short telomere length are associated with IPF risk^41–43^. *RTEL1,* a gene involved in telomere elongation regulation, has not previously been identified in an IPF GWAS, however the collective effect of rare variants in *RTEL1* have been reported as associated with IPF risk^44–47^. The rare variant in the promoter region of *HECTD2* was significantly associated with IPF susceptibility with an OR>7.5 in two out of the three discovery studies, but was unsupported by the replication data. This variant showed some association with subpleural ILA in an independent dataset with a similarly large effect size in the same direction. The ubiquitin E3 ligase encoded by *HECTD2* has been shown to have a pro-inflammatory role in the lung and *HECTD2* polymorphisms may be protective against acute respiratory distress syndrome^48^. The inconsistent evidence for an association suggests that further exploration of the relationship of *HECTD2* to risk of interstitial lung diseases is warranted.

By combining the largest available GWAS datasets for IPF, we were able to confirm 11 of 17 previously reported signals. Of note, the signal at *FAM13A* whilst genome-wide significant in the discovery meta-analysis, was not significant in the Chicago study. Conditional analysis at the 11p15.5 region indicated that previously reported signals at *MUC2* and *TOLLIP* were not independent of the association with the *MUC5B* promoter variant. Previously reported signals at *EHMT2, OBFC1* and *MDGA2* were found to only be associated in one of the discovery studies and showed no evidence of an association with IPF risk in the other two discovery studies.

The IPF susceptibility signals at *DSP, FAM13A,* 7q22.1 *(ZKSCAN1)* and 17q21.31 *(MAPT)* have also been reported as associated with COPD, although with opposite effects (i.e. the allele associated with increased risk of IPF being associated with decreased risk of COPD). Spirometric diagnosis of COPD was based on a reduced FEV1/FVC ratio. In an independent dataset of 400,102 individuals, eight of the IPF signals were associated with decreased FVC and with a comparatively weaker effect on FEV1. This is consistent with the lung function abnormalities associated with IPF, as well as the decreased risk of COPD. We also showed, for the first time, a modest effect of the *MUC5B* risk allele on lung function in the general population.

Using polygenic risk scores, we demonstrated that, despite the relatively large proportion of disease risk explained by the known genetic signals of association reported here, IPF is highly polygenic with potentially hundreds (or thousands) of as-yet unidentified variants associated with disease susceptibility. These unidentified variants may have small effect sizes or be of low frequency meaning larger more powerful studies are needed to detect them. However, they could individually and collectively advance our knowledge of IPF risk and disease mechanisms. This motivates the pursuit of larger GWAS of IPF susceptibility through collation of existing and new IPF case-control data sets and through improved analytic approaches.

A strength of our study was the large sample size compared to previous GWAS and the availability of an independent replication data set. A limitation of the study was that the controls used were generally younger in all studies included and there were differences in sex and smoking distributions in some of the studies. As we had limited information beyond IPF diagnosis status for a large proportion of the individuals included in the studies, we cannot rule out some association with other age-related conditions that are comorbid with IPF. It is worth noting however, that individuals with non-IPF diseases were not excluded from the control sets.

In summary, we report new biological insights into IPF risk and demonstrate that further studies to identify the genetic determinants of IPF susceptibility are needed. Our new signals of association with IPF risk provide increased support for the importance of mTOR signalling in pulmonary fibrosis as well as the possible implication of mitotic spindle-assembly genes.

## Acknowledgements, funding and data sharing

Genome-wide summary statistics are available on request via the corresponding author.

R. Allen is an Action for Pulmonary Fibrosis Research Fellow.

L. Wain holds a GSK/British Lung Foundation Chair in Respiratory Research.

RG. Jenkins is supported by an NIHR Research Professorship (NIHR reference RP-2017-08-ST2-014).

I. Noth: National Heart Lung and Blood Institute (R01HL130796).

B. Guillen is funded by Agencia Canaria de Investigación, Innovación y Sociedad de la Información (TESIS2015010057) co-funded by European Social Fund.

J. Oldham: National Heart Lung and Blood Institute (K23HL138190).

C. Flores: Spanish Ministry of Science, Innovation and Universities (grant RTC-2017-6471-1; MINECO/AEI/FEDER, UE) co-financed by the European Regional Development Funds (ERDF) ‘A way of making Europe’ from the European Union, and by the agreement 0A17/008 with Instituto Tecnológico y de Energías Renovables (ITER) to strengthen scientific and technological education, training, research, development and innovation in Genomics, Personalized Medicine and Biotechnology. The Spain Biobank array genotyping service was carried out at CEGEN-PRB3-ISCIII; which is supported by PT17/0019, of the PE I+D+i 2013-2016, funded by Instituto de Salud Carlos III, and co-financed by ERDF.

P. Molyneaux is an Action for Pulmonary Fibrosis Research Fellow.

M Obeidat is a fellow of the Parker B Francis Foundation and a Scholar of the Michael Smith Foundation for Health Research (MSFHR).

B. Hobbs: NIH K08 HL136928, Parker B. Francis Research Opportunity Award.

M. Cho and G Hunninghake: This work was supported by NHLBI grants R01HL113264 (M.H.C), R01HL137927 (M.H.C.), R01HL135142 (M.H.C. and G.M.H) and R01111024 (G.M.H.). The content is solely the responsibility of the authors and does not necessarily represent the official views of the NIH. The funding body has no role in the design of the study and collection, analysis, and interpretation of data and in writing the manuscript.

T. Maher is supported by an NIHR Clinician Scientist Fellowship (NIHR Ref: CS-2013-13-017) and a British Lung Foundation Chair in Respiratory Research (C17-3).

M. Tobin is supported by a Wellcome Trust Investigator Award (WT202849/Z/16/Z). The research was partially supported by the National Institute for Health Research (NIHR) Leicester Biomedical Research Centre; the views expressed are those of the author(s) and not necessarily those of the National Health Service (NHS), the NIHR or the Department of Health.

I. Hall was partially supported by the NIHR Nottingham Biomedical Research Centre; the views expressed are those of the author(s) and not necessarily those of the NHS, the NIHR or the Department of Health.

I. Sayers: MRC (G1000861) and Asthma UK (AUK-PG-2013-188).

D. Furniss was supported by an Intermediate Fellowship from the Wellcome Trust (097152/Z/11/Z). This work was partially supported by the National Institute for Health Research (NIHR) Oxford Biomedical Research Centre.

V. Navaratnam is funded by an NIHR Clinical Lectureship.

G. Gudmundsson is supported by project grant 141513-051 from the Icelandic Research Fund and Landspitali Scientific Fund A-2016-023, A-2017-029 and A-2018-025.

A. Manichaikul and D. Lederer: MESA and the MESA SHARe project are conducted and supported by the National Heart, Lung, and Blood Institute (NHLBI) in collaboration with MESA investigators. Support for MESA is provided by contracts HHSN268201500003I, N01-HC-95159, N01-HC-95160, N01-HC-95161, N01-HC-95162, N01-HC-95163, N01-HC-95164, N01-HC-95165, N01-HC-95166, N01-HC-95167, N01-HC-95168, N01-HC-95169, UL1-TR-000040, UL1-TR-001079, UL1-TR-001420, UL1-TR-001881, and DK063491. Funding for SHARe genotyping was provided by NHLBI Contract N02-HL-64278. Genotyping was performed at Affymetrix (Santa Clara, California, USA) and the Broad Institute of Harvard and MIT (Boston, Massachusetts, USA) using the Affymetrix Genome-Wide Human SNP Array 6.0. This work was supported by NIH grants R01 HL131565 (A.M.), R01 HL103676 (D.J.L.) and R01 HL137234 (D.J.L.).

This research has been conducted using the UK Biobank Resource under application 8389.

This research used the ALICE and SPECTRE High Performance Computing Facilities at the University of Leicester.

