## Supplementary material for "Genome-wide association study of susceptibility to idiopathic pulmonary fibrosis"

### Appendix

ii) *DEPTOR* - Lung - Colocalisation probability = 89.2% 54
iii) *DEPTOR* - Lung - Colocalisation probability = 89.5% 55
iv) *DEPTOR* - Lung - Colocalisation probability = 89.9% 56
v) *DEPTOR* - Skin (Not sun exposed) - Colocalisation probability = 90.0% 57
vi) *DEPTOR* - Skin (Sun exposed) - Colocalisation probability = 86.5% 58
vii) *DEPTOR* - Whole blood - Colocalisation probability = 93.7% 59
viii) *TAF2* - Colon (Transverse) - Colocalisation probability = 87.5% 60
ix) RP11-760H22.2 - Adipose (Subcutaneous) - Colocalisation probability = 84.9% 61
x) RP11-760H22.2 - Colon (Sigmoid) - Colocalisation probability = 88.6% 62
xi) RP11-760H22.2 - Lung - Colocalisation probability = 90.0% 63
xii) KB-1471A8.1 - Adipose (Subcutaneous) - Colocalisation probability = 85.6% 64
xiii) KB-1471A8.1 - Adipose (Visceral) - Colocalisation probability = 90.9% 65
xiv) KB-1471A8.1 - Skin (Sun exposed) - Colocalisation probability = 88.7% 66
Supplementary Figure 10 - FORGE analysis for enrichment in regulatory regions 67
Supplementary Figure 11 - GARFIELD analysis for enrichment in DNase I hypersensitivity sites by tissue 68
Supplementary Figure 12 - Polygenic risk score in target dataset by *P* threshold used 69
Supplementary References 70

### Supplementary Methods

#### Summary of previously reported studies

The Chicago study[1] consisted of individuals of 541 IPF cases and 542 controls. All individuals were unrelated, of European-American ancestry and had call rate > 97%. Sex mismatches were removed and controls were selected so they were genetically matched to a case based on the first 4 principal components. All individuals were genotyped using the Affymetrix 6.0 SNP array.

The Colorado Study[2,3] consisted of 1,616 IIP (idiopathic interstitial pneumonia) cases and 4,683 controls selected such that they were genetically similar to the cases based on IBS (identical by state) estimates. Of the cases, 101 were removed from this study as they had also been included in the Chicago study (see **Duplicated individuals between studies** section). All individuals were self-reported as non-Hispanic white and were removed if they had call rate < 98%, were sex mismatches, had genome-wide heterozygosity more than four standard deviations away from the mean or were genetic outliers based on IBS estimates. All individuals were genotyped using the Illumina Human 660W Quad BeadChip array.

The UK study[4] consisted of 612 IPF cases and 3,366 controls selected from UK Biobank such that they followed a similar age, sex and smoking distribution to the cases. Individuals were removed if they had high missingness (call rate < 95%), were heterozygote outliers, were ancestry outliers based on principal components or were sex mismatches. All individuals were of European ancestry and were unrelated. Genotyping of cases was performed using the Affymetrix UK BiLEVE array. For the controls, 1,231 were genotyped using the Affymetrix UK BiLEVE array and the remaining 2,135 were genotyped using the similar Affymetrix UK Biobank array.

The Genentech study[5] consisted of 664 unrelated European IPF cases taken from the ASCEND, CAPACITY and RIFF clinical trials and 1,874 unrelated European non-IPF controls taken from the EXCELS, SUMMACTA, LITHE and OPTION clinical trials. The original Genentech cohort also included IPF cases from the Vanderbilt, UCSF and INSPIRE cohorts, however as these had been included in other studies included in the discovery analysis, these individuals were excluded. Individuals were sequenced using the HiSeq X Ten platform (Illumina) to an average read depth of 30X. Individuals were excluded from analyses if they had a call rate < 10%, had excess heterozygosity, were ancestry outliers or were aged less than 40 years old.

#### Recruitment and genotyping of cases for UUS study

A total of 1,366 individuals were recruited and sent for genotyping from 9 study cohorts; ACE (Anticoagulant Effectiveness in IPF, n = 98), PANTHER (Prednisone, Azathioprine, and N-Acetylcysteine: A Study That Evaluates Response in Idiopathic Pulmonary Fibrosis, n = 166), UCD (University of California Davis, n = 54), Chicago (n = 314), ILDCON (Interstitial Lung Disease Controls, n = 33), RECITAL (Rituximab Versus Cyclophosphamide in Connective Tissue Disease-ILD, n = 44), UCSF (University of California San Francisco, n = 53), PROFILE (n = 554) and Spain (n = 50).

Cases in the replication study were genotyped by Affymetrix on the Affymetrix Axiom UK Biobank array, apart from the 50 Spanish IPF cases who were genotyped on the Axiom Spain Biobank array. The UK Biobank array was designed to optimise imputation quality of common (MAF [minor allele frequency] > 5%) and low-frequency (MAF 1% to 5%) variants in a European population, measure rare functional variation and to include custom content of known genetic associations with a variety of traits (including *MUC5B* promoter and *TERT* variants with known associations with IPF).

Quality control was performed on the individuals on the two arrays separately before being merged for the selection of controls and for imputation.

#### Quality control for UUS study

##### Quality control for individuals genotyped on UK Biobank array

For the individuals genotyped on the UK Biobank array, the following quality control measures were applied.

1. **Affymetrix quality control:** Individuals were removed if they had scanning issues, failed dish QC and or had a sample call rate < 97% in step 1 genotype calling. The genotype calling and quality control was originally performed by Affymetrix and was repeated using Axiom Analysis Suite and APT (Analysis Power Tools). All three calling methods gave the same results.
2. **Individual call rate:** Individuals were excluded for having a final individual call rate < 95%.
3. **Sex mismatches:** Genetic sex was inferred using PLINK v1.9. Individuals who had a recorded sex different to that inferred from their genetic sample and were not included in further analyses.
4. **High heterozygosity:** Individuals were excluded if they had a high heterozygosity rate (defined as more than 5 standard deviations above the mean after adjusting for ancestry). Heterozygosity rates were calculated using autosomal variants in Hardy-Weinberg equilibrium with MAF > 1% and variant call rate > 95%.
5. **Non-IPF cases:** Individuals found to not be IPF cases were removed.
6. **Duplicates:** Duplicates were identified using KING on all samples, including those individuals already excluded for failing other quality control measures. The duplicate analysis was performed on autosomal variants in Hardy-Weinberg equilibrium ($P > 10^{-6}$), had call rate > 95%, MAF > 1% and not found in regions of high linkage disequilibrium (LD, namely positions 44Mb to 51.5Mb on chromosome 5, 25Mb to 33.5Mb on chromosome 6 [HLA region], 8Mb to 12Mb on chromosome 8 and 45Mb to 57Mb on chromosome 11). If the phenotype data suggested the same person had been recruited twice then the sample with the highest call rate was kept. In instances where the phenotype data suggested there had been a potential genetic sample mix-up, both pairs were removed. Duplicates were also identified between studies. More details on this analysis can be found in the "Duplicated individuals between studies" section in the supplementary methods.
7. **Ancestry:** Ancestry was inferred from the genetic data using principal components analysis. Principal components were calculated using PLINK v1.9 on autosomal variants with MAF > 1%, in Hardy-Weinberg equilibrium, were variants included in HapMap, had genotyping call rate > 95% and were not in regions of high LD. Variants were pruned using an $r^2$ threshold of 0.1. Principal components were calculated for the individuals who passed Affymetrix QC, had call rate > 95%, were not sex mismatches and were not duplicates and for all unrelated samples from the HapMap project (a collection of genotyped individuals from multiple populations). K-means clustering on the first two principal components was used to define ancestry groups. The number of clusters was increased until a cluster was formed which contained all European HapMap samples and no HapMap samples of other ancestries. Using seven clusters was found to form a cluster of European samples. Individuals in the other ancestry clusters were not included in further analyses.
8. **Relatedness:** Relatedness between individuals passing quality control measures was calculated using KING. First-degree relatives were defined as those with kingship coefficient between 0.177 and 0.354 and second-degree relatives as a kingship coefficient between 0.0884 and 0.177. When first and second-degree relatives were identified, the individual with the lower genotyping call rate was removed from further analyses.
9. **RECITAL samples:** Cases from the RECITAL study were excluded for not having a definite diagnosis of IPF.

10. **Relatedness with the discovery:** Individuals who were first or second-degree relatives with an individual in the discovery analysis were also excluded. Relatedness was estimated using KING.

Of the 1,316 individuals genotyped on the UK Biobank array, 764 unrelated European IPF cases passed quality control.

**Quality control of individuals genotyped on Spain Biobank array for UUS study**
For the individuals genotyped on the Spanish custom array, the following quality control measures were applied.

1. **Affymetrix QC:** Individuals were removed if they had scanning issues, failed dish QC and or had a sample call rate < 97% in step 1 genotype calling. The genotype calling and quality control was performed using Axiom Analysis Suite.
2. **Individual call rate:** Individuals were excluded for having a final individual call rate < 95%.
3. **Sex mismatches:** Sex was inferred from the genetic data using PLINK v1.9. Individuals were excluded from future analyses if their genetically inferred sex was different to their recorded sex.
4. **High heterozygosity:** Heterozygosity rates were estimated using PLINK v1.9 on autosomal variants with call rate > 95%, in Hardy-Weinberg equilibrium ($P > 10^{-6}$) and MAF > 1%. Heterozygosity rates were adjusted for ancestry and individuals found to have a high genome-wide ancestry-adjusted heterozygosity rate (more than 5 standard deviations above mean) were removed.
5. **Duplicates and relatedness:** Duplicates and relatedness between individuals was calculated using PLINK v1.9 on autosomal variants with call rate > 95%, in Hardy-Weinberg equilibrium ($P > 10^{-6}$), MAF > 1% and not in a region of high LD. Variants were pruned using an $r^2$ threshold of 0.13. In instances of duplicates, first or second-degree relatives the sample with the lowest genotyping call rate was removed (apart from instances where it appeared genetic sample mix-up had occurred in which case both individuals were removed).
6. **Ancestry:** Ancestry outliers for the IPF cases passing previous quality control measures were inferred from the genetic data through principal components analysis. Principal components were calculated using PLINK v1.9 on autosomal variants with MAF > 1%, in Hardy-Weinberg equilibrium, were variants included in HapMap, had genotyping call rate > 95% and were not in regions of high LD. Variants were pruned using an $r^2$ threshold of 0.22 (the lowest $r^2$ value that left more than 100,000 variants). Principal components were calculated alongside HapMap samples. Individuals who had either the first or second principal component greater than two standard deviations away from the mean were deemed to be ancestry outliers and removed from further analyses.
7. **Relatedness with the discovery:** Individuals who were first or second-degree relatives with an individual in the discovery analysis were also excluded. Relatedness was estimated using KING.

Of the 50 individuals genotyped on the Spanish custom array, 39 unrelated European IPF cases passed quality control.

**Selection and quality control of controls for UUS study**
Controls were selected from UK Biobank such that they were European (defined by k-means clustering of first two principal components), not a possible ILD case, were related to another UK Biobank individual, or were a control in the UK study.

ILD cases in UK Biobank were identified using the self-reported questionnaire (field 20002 - Non-cancer illness code, self-reported) and from HES data (i.e. any hospital episode recorded with ICD10 codes J84, J841, J848, J849 or ICD9 codes 516, 5160, 5161, 5162, 5163, 51630, 51631, 51632, 51633, 51634, 51635, 51636, 51637, 5164, 5165, 5166, 51661, 51662, 51663, 51664, 51669, 5168, 5169).

Of the 300,909 individuals passing the above selection criteria, 10,000 were selected as controls such that they followed a similar sex and smoking distribution to that seen in the IPF cases.

#### Imputation of all studies

Each study was imputed separately to the Haplotype Reference Consortium reference panel using the Michigan Imputation Server. Only variants in Hardy-Weinberg equilibrium ($P > 10^{-6}$), had call rate > 95% and had MAF > 1% were considered.

When more than one genotyping array was used in a study (i.e. the UK study and replication study) only variants that appeared on all arrays used in that study were included in the imputation. For the replication study, the concordance between the imputed genotypes and the directly measured genotypes not included in the imputation (i.e. due to not being on all the arrays used) was found to be high (concordance = 99.6%).

#### Duplicated individuals between studies

It is possible for individuals to be recruited to multiple studies. To ensure the studies included in this analysis were completely independent, individuals who had been recruited to multiple studies were identified from the genetic data and removed. This was conducted using PLINK v1.9 and verified using KING.

Variants were included in the PLINK IBD (identical by descent) analysis if they were on an autosome, had call rate > 95%, MAF > 1%, in Hardy-Weinberg equilibrium ($P > 10^{-6}$) and not in a region of high LD. Variants were pruned using an $r^2$ threshold of 0.3 leaving 120,864 variants to be included in the IBD analysis (as a sensitivity analysis an $r^2$ threshold of 0.1 was used and the same results were observed). Pairs of genetic samples with PI_HAT > 0.8 were considered as duplicates.

The duplicate analysis was repeated using KING on autosomal variants with call rate > 95%, MAF > 1%, in Hardy-Weinberg equilibrium ($P > 10^{-6}$) and not in an area of high linkage disequilibrium. Duplicates were identified as those with kingship > 0.354. The KING analysis gave the same results as seen in the analysis performed using PLINK.

#### Genomic control

Genomic control was applied for each study in the discovery meta-analysis ($\lambda = 1.027$ in UK, $\lambda = 1.065$ in Colorado and $\lambda = 1.030$ in Chicago). Genomic control was also applied to the meta-analysis of all three studies ($\lambda = 1.016$).

#### Conditional analyses

To identify additional independent signals within each locus in the discovery meta-analysis, conditional analyses were performed by repeating the association analyses for all variants within 1Mb of the sentinel variant, adjusting for the sentinel variant in each study separately and then meta-analysing the results. Variants reaching genome-wide significance after conditioning on the top variant were deemed as independent signals and analyses were repeated until no more independent signals in the region were identified.

#### Bayesian fine-mapping

Credible sets were calculated for each novel signal to produce a set of variants likely to contain the causal variant at 95% confidence (under the assumption there is a single causal variant and that variant had been measured). Posterior probabilities of the variant being causal were calculated for all variants within 1Mb of the sentinel variant and in at least weak LD with the sentinel variant ($r^2 > 0.1$) in the discovery meta-analysis. Posterior probabilities were calculated from approximate Bayes factors (ABFs) using the formula proposed by Wakefield[6]:

$$\mathrm{ABF} = \frac{1}{\sqrt{1 - \frac{W}{V + W}}} \exp\left(-\frac{Z^2}{2}\frac{W}{V + W}\right)$$

where $W$ is the Wakefield prior (which we set to 0.4 which is equivalent to a 95% belief that departure from the null model for the relative risk is less than 1.5), $Z$ is the Z statistic for the variant and $V$ is the variance of the effect size.

The approximate posterior probability was set to equal the ABF for that variant divided by the sum of ABFs for all variants in the signal.  Variants were added to the credible set until the sum of the posterior probabilities was greater than or equal to 0.95.

#### Colocalisation

Analyses were performed to investigate whether the novel IPF GWAS signals observed in the discovery analyses colocalised with the eQTL signals identified in the lung eQTL database, NESDA-NTR and GTEx. Analyses were performed using the coloc package in R v3.5.1 on all variants in the region with $P < 0.01$ in either the IPF GWAS analysis or eQTL analysis.

The coloc package implements the colocalisation approach described by Giambartolomei et al[7]. In summary, it uses approximate Bayes factors to estimate the probability of each of the following models:

- $H_o$: There is no association in the region with either IPF risk or the eQTL result
- $H_1$: There is an association in the region with IPF but not with the expression of the gene
- $H_2$: There is an association in the region with the expression of the gene but not with IPF
- $H_3$: There is an association in the region with both IPF and the expression of the gene but these are driven by two different variants
- $H_4$: There is an association in the region with both IPF and the expression of the gene which is driven by the same variant.

We took colocalisation to be when the probability of $H_4$ (i.e. the same variant drives IPF risk and the expression of the gene) was greater than 80%.

#### Functional follow-up

##### VEP

All variants in the credible sets were annotated using VEP[8]. Variants were defined as deleterious if they were recorded as either "deleterious" in SIFT, "probably damaging" in PolyPhen, "likely deleterious" from the CADD score, "likely disease causing" in REVEL, "damaging" in MetaLR or "high" in MutationAssessor.

**DeepSEA**
DeepSEA[9] (deep learning-based sequence analyzer) is a deep learning method to predict chromatin effects. The variant with the highest posterior probability in each of the credible sets for the 14 IPF risk signals identified by the discovery meta-analysis was included in the DeepSEA analysis.
We reported functional effects for any chromatin feature and lung-related cell line that had an E-value < 0.05 (i.e. the expected proportion of SNPs with larger predicted effect for this chromatin feature based on empirical distributions of predicted effects for 1000 Genomes SNPs) and an absolute difference in probability of > 0.1 (threshold for "high confidence") between the reference and alternative allele.

**FORGE**
FORGE[10] (Functional element overlap analysis of the results of GWAS experiments) is a tool for identifying whether signals in a GWAS are enriched in DNase I hypersensitivity sites in specific tissues. The variant with the highest posterior probability in each of the credible sets for the 14 IPF risk signals identified by the discovery meta-analysis was included in the FORGE analysis. Enrichment was tested in 299 cell lines across 24 tissues including lung and foetal lung.

**GARFIELD**
GARFIELD[11] (GWAS analysis of regulatory and functional information enrichment with LD correction) is an analysis tool to test if GWAS signals are enriched in functional features. Variants meeting a *P* threshold in the IPF discovery genome-wide analysis were tested for enrichment (*P* thresholds of $5 \times 10^{-8}$ and $5 \times 10^{-5}$ were used). Enrichment was tested in DNase I hypersensitivity sites in 424 tissues.

**SNPsea**
SNPsea[12] is a method to identify if gene expression is altered by a set of variants in different cell types or pathways. For this analysis, the variants with the highest posterior probability in each of the credible sets for the 14 IPF risk loci were entered. Genes are implicated using an LD matrix and the expression of these genes are investigated. A score based on the expression of these genes is calculated and compared to a score generated when selecting a random set of variants as the input. The IPF risk loci were tested for enrichment in biological pathways and epithelial cell types were tested.

Pathway-specific gene-expression was calculated using gene expression from 1,751 pathways in Gene Ontology. Expression of genes in four epithelial cell types (normal AT2 cells, indeterminate cells, basal and club/goblet cells) was calculated from lung tissue from six IPF cases and three healthy controls using single cell RNA sequencing data from Xu et al[13].

Gene expression was deemed to be enriched in tissues or pathways if they met a Bonferroni corrected *P* threshold.

#### Association with other respiratory traits
A genome-wide association analysis of interstitial lung abnormalities (ILA) was conducted by meta-analysis of results from the AGES, COPDGene NHW, ECLIPSE, Framingham, MESA white and SPIROMICS studies. Two analyses were performed; firstly defining cases as any individual with any ILA (n = 1,699) and controls as any individual without an ILA (n = 10,247), and secondly defining cases as those with a subpleural subtype of ILA (n = 1,287) and controls as individuals without any ILA (n = 10,247).

Association with lung function was assessed using data from a genome-wide association study meta-analysis of lung function for 400,102 individuals of European ancestry in UK Biobank and the

SpiroMeta consortium[14]. The measures of lung function analysed were $FEV_1$ (a measure of how much air an individual can forcibly exhale in the first second), FVC (forced vital capacity, i.e. the total volume of air forcibly exhaled), the ratio of $FEV_1$/FVC (a measure used in the diagnosis of chronic obstructive pulmonary disease) and PEF (peak expiratory flow, i.e. the highest airflow), which were all measured through spirometry.

#### Supplementary Tables

##### Supplementary Table 1 - Discovery study level results

Odds ratios are presented treating the minor allele as the effect allele. Minor allele frequency (MAF) was the allele frequency across the three studies and info is the imputation quality in that study.

| Chr | Pos | Sentinel rsid | $r^2$ with previous reported sentinel | Locus | Major allele | Minor allele | MAF | Chicago | | | Colorado | | | UK | | | Discovery meta-analysis | |
|---|---|---|---|---|---|---|---|---|---|---|---|---|---|---|---|---|---|---|
| | | | | | | | | Info | OR [95% CI] | *P* | Info | OR [95% CI] | *P* | Info | OR [95% CI] | *P* | OR [95% CI] | *P* |
| **i) Novel signals meeting significance criteria** | | | | | | | | | | | | | | | | | | |
| 3 | 44902386 | rs78238620 | - | *KIF15* | T | A | 5.3% | 0.97 | 1.74 [1.16, 2.60] | 0.007 | 0.98 | 1.47 [1.23, 1.78] | $4.01\times10^{-5}$ | 0.99 | 1.77 [1.35, 2.33] | $4.54\times10^{-5}$ | 1.58 [1.37, 1.83] | $5.12\times10^{-10}$ |
| 7 | 1909479 | rs12699415 | - | *MAD1L1* | G | A | 42.0% | 0.97 | 1.43 [1.20, 1.69] | $5.31\times10^{-5}$ | 0.98 | 1.23 [1.12, 1.33] | $3.67\times10^{-6}$ | 0.99 | 1.30 [1.15, 1.47] | $3.51\times10^{-5}$ | 1.28 [1.19, 1.37] | $7.15\times10^{-13}$ |
| 8 | 120934126 | rs28513081 | - | *DEPTOR* | A | G | 42.8% | 0.99 | 0.78 [0.66, 0.93] | 0.005 | 0.99 | 0.84 [0.77, 0.91] | $4.69\times10^{-5}$ | 0.99 | 0.79 [0.70, 0.89] | $1.94\times10^{-4}$ | 0.82 [0.76, 0.87] | $1.20\times10^{-9}$ |
| 10 | 93271016 | rs537322302 | - | *HECTD2* | C | G | 0.3% | - | - | - | 0.55 | 7.52 [2.46, 23.0] | $3.98\times10^{-4}$ | 0.90 | 8.04 [3.10, 20.8] | $1.79\times10^{-5}$ | 7.82 [3.77, 16.2] | $3.43\times10^{-8}$ |
| 20 | 62324391 | rs41308092 | - | *RTEL1* | G | A | 2.1% | 0.57 | 2.06 [1.03, 4.12] | 0.040 | 0.79 | 2.10 [1.54, 2.86] | $2.61\times10^{-6}$ | 0.94 | 2.18 [1.41, 3.37] | $4.86\times10^{-4}$ | 2.12 [1.67, 2.69] | $7.65\times10^{-10}$ |
| **ii) Previously reported signals that reached genome-wide significance in discovery analysis** | | | | | | | | | | | | | | | | | | |
| 3 | 169481271 | rs12696304 | 0.97 | *LRRC34 /TERC* | C | G | 27.9% | 0.99 | 1.38 [1.15, 1.66] | $5.57\times10^{-4}$ | 0.98 | 1.33 [1.21, 1.46] | $5.52\times10^{-9}$ | 1.00 | 1.22 [1.06, 1.41] | 0.005 | 1.31 [1.21, 1.40] | $7.09\times10^{-13}$ |
| 4 | 89885086 | rs2013701 | 0.28 | *FAM13A* | G | T | 48.7% | 1.00 | 0.94 [0.79, 1.12] | 0.496 | 1.00 | 0.78 [0.72, 0.85] | $9.16\times10^{-9}$ | 1.00 | 0.72 [0.63, 0.81] | $2.27\times10^{-7}$ | 0.78 [0.74, 0.84] | $3.30\times10^{-13}$ |
| 5 | 1282414 | rs7725218 [a] | 0.55 | *TERT* | G | A | 32.5% | 0.52 | 0.83 [0.69, 1.00] | 0.051 | 0.90 | 0.68 [0.62, 0.74] | $4.88\times10^{-17}$ | 1.00 | 0.76 [0.66, 0.86] | $2.68\times10^{-5}$ | 0.72 [0.67, 0.77] | $1.54\times10^{-20}$ |
| 6 | 7563232 | rs2076295 | Same | *DSP* | T | G | 46.9% | 0.98 | 1.19 [1.00, 1.42] | 0.044 | 1.00 | 1.45 [1.33, 1.58] | $9.56\times10^{-18}$ | 0.99 | 1.66 [1.47, 1.87] | $8.81\times10^{-16}$ | 1.46 [1.37, 1.56] | $2.79\times10^{-30}$ |
| 7 | 99630342 | rs2897075 | 0.72 | 7q22.1 | C | T | 39.1% | 0.98 | 1.30 [1.09, 1.54] | 0.003 | 0.99 | 1.34 [1.23, 1.46] | $2.14\times10^{-11}$ | 1.00 | 1.19 [1.05, 1.36] | 0.008 | 1.30 [1.21, 1.38] | $3.10\times10^{-14}$ |

| | | | | | | | | | | | | | | | | | | |
|---|---|---|---|---|---|---|---|---|---|---|---|---|---|---|---|---|---|---|
| 11 | 1241221 | rs35705950 | Same | *MUC5B* | G | T | 14.9% | - | - | - | 0.77 | 4.51 [3.99, 5.09] | $1.14\times10^{-128}$ | 0.92 | 5.64 [4.72, 6.73] | $3.99\times10^{-81}$ | 4.84 [4.37, 5.36] | $1.18\times10^{-203}$ |
| 13 | 113534984 | rs9577395 | 0.86 | *ATP11A* | C | G | 20.7% | 0.77 | 0.75 [0.61, 0.93] | 0.008 | 0.99 | 0.75 [0.68, 0.84] | $9.27\times10^{-8}$ | 1.00 | 0.81 [0.69, 0.95] | 0.008 | 0.77 [0.71, 0.83] | $1.34\times10^{-10}$ |
| 15 | 40720542 | rs59424629 | 0.97 | *IVD* | T | G | 46.1% | 0.98 | 0.64 [0.54, 0.76] | $5.39\times10^{-7}$ | 0.99 | 0.78 [0.71, 0.85] | $3.19\times10^{-9}$ | 1.00 | 0.81 [0.71, 0.92] | $9.85\times10^{-4}$ | 0.76 [0.71, 0.82] | $7.30\times10^{-16}$ |
| 15 | 86097216 | rs62023891 | 0.54 | *AKAP13* | G | A | 30.0% | 0.97 | 1.27 [1.06, 1.53] | 0.011 | 0.99 | 1.25 [1.14, 1.37] | $2.98\times10^{-5}$ | 0.99 | 1.30 [1.13, 1.49] | $1.86\times10^{-4}$ | 1.27 [1.18, 1.36] | $1.27\times10^{-10}$ |
| 17 | 44214888 | rs2077551 | 0.90 | *MAPT* | T | C | 18.6% | 0.85 | 0.63 [0.51, 0.79] | $4.50\times10^{-5}$ | 0.86 | 0.71 [0.64, 0.79] | $3.78\times10^{-10}$ | 0.96 | 0.75 [0.65, 0.87] | $2.01\times10^{-4}$ | 0.71 [0.65, 0.77] | $2.83\times10^{-16}$ |
| 19 | 4717672 | rs12610495 | Same | *DPP9* | A | G | 30.5% | - | - | - | 1.00 | 1.29 [1.17, 1.41] | $7.84\times10^{-8}$ | 0.97 | 1.37 [1.20, 1.57] | $3.91\times10^{-6}$ | 1.31 [1.22, 1.42] | $2.92\times10^{-12}$ |
| **iii) Previously reported signals that do not reach genome-wide significance in the discovery meta-analysis** | | | | | | | | | | | | | | | | | | |
| 6 | 31864547 | rs7887 | - | *EHMT2* | G | T | 33.9% | 0.96 | 0.92 [0.77, 1.11] | 0.388 | 0.99 | 0.84 [0.77, 0.92] | $1.10\times10^{-4}$ | 1.00 | 1.03 [0.91, 1.18] | 0.625 | 0.90 [0.84, 0.96] | 0.002 |
| 10 | 105672842 | rs11191865 | - | *OBFC1* | G | A | 49.1% | 1.00 | 0.98 [0.82, 1.16] | 0.809 | 1.00 | 1.27 [1.16, 1.38] | $5.08\times10^{-8}$ | 0.99 | 1.05 [0.93, 1.19] | 0.455 | 1.16 [1.09, 1.24] | $8.91\times10^{-6}$ |
| 14 | 48040375 | rs7144383 | - | *MDGA2* | A | G | 11.2% | 0.98 | 1.81 [1.37, 2.38] | $2.94\times10^{-5}$ | 0.88 | 1.03 [0.90, 1.18] | 0.671 | 0.98 | 0.95 [0.78, 1.15] | 0.568 | 1.09 [0.98, 1.21] | 0.119 |
| **iv) Previously reported signals in the 11p15.5 region after conditioning on rs35705950** [b] | | | | | | | | | | | | | | | | | | |
| 11 | 1093945 | rs7934606 | - | *MUC2* | C | T | 44.9% | - | - | - | 1.00 | 0.93 [0.85, 1.02] | 0.109 | 0.94 | 1.00 [0.87, 1.16] | 0.956 | 0.95 [0.88, 1.03] | 0.189 |
| 11 | 1312706 | rs111521887 | - | *TOLLIP* | C | G | 19.8% | - | - | - | 0.99 | 1.00 [0.89, 1.12] | 0.972 | 0.99 | 1.00 [0.85, 1.19] | 0.965 | 1.00 [0.91, 1.10] | 0.996 |
| 11 | 1325829 | rs5743890 | - | *TOLLIP* | T | C | 13.8% | - | - | - | 0.93 | 0.84 [0.74, 0.95] | 0.006 | 0.95 | 0.88 [0.72, 1.07] | 0.193 | 0.85 [0.76, 0.95] | 0.002 |

[a] This variant was the most significant variant for this signal in the discovery meta-analysis. Although this variant did not quite reach nominal significance ($p<0.05$) in the Chicago study other variants in the signal did reach nominal in each study, had consistent direction of effects in each study and were genome-wide significant in the discovery meta-analysis

[b] The *MUC5B* promoter polymorphism rs35705950 was not imputed in the Chicago study so it was not possible to perform the conditional analysis

**Supplementary Table 2 - Summary of eQTL analysis for novel IPF susceptibility signals**

The table below contains all of the genes for which at least one of the variants in the credible set was recorded as an eQTL variant and the tissue this was recorded in. This table also includes the colocalisation probability and IPF risk signals that colocalise with the expression of a gene (taken to be probability > 80%) are shown in green.

| Chromosome | GWAS sentinel (risk allele) | eQTL gene | Source | eQTL Tissue (and probe if multiple probes used) | Risk allele effect on gene expression | eQTL sentinel | Colocalisation probability |
|---|---|---|---|---|---|---|---|
| 3 | rs78238620_A | *KIF15* | GTEx | Brain - Putamen | Decrease | rs149000267 | 95.6% |
| | | *TMEM42* | GTEx | Thyroid | Decrease | rs80059929 | 93.1% |
| | | *KIAA1143* | GTEx | Adipose - Subcutaneous | Decrease | rs6792299 | 0.0% |
| 7 | rs12699415_A | *MAD1L1* | NESDA-NTR [a] | Whole blood (11724206_a_at) | Increase | rs35578480 | 0.1% |
| | | | | Whole blood (11724207_x_at) | Increase | rs35578480 | 0.1% |
| | | | | Whole blood (11737802_a_at) | Increase | rs35578480 | 0.0% |
| | | | GTEx | Heart - Atrial Appendage | Decrease | rs57193072 | 95.3% |
| | | | | Nerve - Tibial | Increase | rs7803147 | 36.3% |
| | | *FTSJ2* | GTEx | Adipose - Visceral | Decrease | rs34418140 | 0.0% |
| | | | | Artery - Aorta | Decrease | rs73673559 | 0.1% |
| | | | | Artery - Tibial | Decrease | rs57431109 | 0.0% |
| | | | | Brain - Cerebellum | Decrease | rs34418140 | 0.0% |
| | | | | Esophagus - Muscularis | Decrease | rs7810970 | 0.0% |
| | | | | Muscle - Skeletal | Decrease | rs4719462 | 0.0% |
| | | | | Testis | Decrease | rs7810970 | 0.0% |
| | | AC110781.3 | GTEx | Brain - Frontal Cortex | Increase | rs6952808 | 64.8% |
| | | | | Brain - Nucleus accumbens | Increase | rs4236272 | 51.9% |
| | | | | Testis | Increase | rs10237989 | 60.5% |
| 8 | rs28513081_A | *DEPTOR* | NESDA-NTR [a] | Whole Blood (11751331_a_at) | Increase | rs55892034 | 93.7% |
| | | | Lung eQTL [b] | Lung (100154484_TGI_at) | Decrease | rs1519812 | 89.5% |
| | | | | Lung (100312124_TGI_at) | Decrease | rs1519812 | 89.9% |
| | | | GTEx | Brain - Spinal Cord | Decrease | rs72673678 | 55.2% |
| | | | | Colon - Sigmoid | Decrease | rs10217077 | 89.6% |
| | | | | Colon - Transverse | Decrease | rs7005380 | 58.1% |
| | | | | Esophagus - Gastroesophageal Junction | Decrease | rs56177421 | 0.9% |
| | | | | Esophagus - Mucosa | Decrease | rs56177421 | 41.6% |
| | | | | Esophagus - Muscularis | Decrease | rs56177421 | 0.9% |
| | | | | Lung | Decrease | rs10217077 | 89.2% |
| | | | | Muscle - Skeletal | Decrease | rs7818296 | 8.7% |

| | | | | | | | |
|---|---|---|---|---|---|---|---|
| | | | | Skin - Not Sun Exposed | Decrease | rs7814520 | 90.0% |
| | | | | Skin - Sun Exposed | Decrease | rs1467044 | 86.5% |
| | | *DSCC1* | Lung eQTL | Lung (100129753_TGI_at) | Increase | rs7815122 | 0.0% |
| | | | | Lung (100132546_TGI_at) | Increase | rs7815122 | 0.0% |
| | | | | Lung (100301732_TGI_at) | Increase | rs55741337 | 0.0% |
| | | | GTEx | Artery - Tibial | Increase | rs113408398 | 0.0% |
| | | | | Muscle - Skeletal | Increase | rs77647593 | 0.0% |
| | | | | Skin - Sun Exposed | Increase | rs28700049 | 0.0% |
| | | RP11-760H22.2 | GTEx | Adipose - Subcutaneous | Increase | rs10217077 | 84.9% |
| | | | | Brain - Spinal cord | Decrease | rs7825920 | 46.8% |
| | | | | Colon - Sigmoid | Decrease | rs6469868 | 88.6% |
| | | | | Esophagus - Gastroesophageal Junction | Decrease | rs56177421 | 1.0% |
| | | | | Esophagus - Muscularis | Decrease | rs12541326 | 0.9% |
| | | | | Lung | Decrease | rs796666096 | 90.0% |
| | | KB-1471A8.1 | GTEx | Adipose - Subcutaneous | Increase | rs1467044 | 85.6% |
| | | | | Adipose - Visceral | Increase | rs7818471 | 90.9% |
| | | | | Muscle - Skeletal | Increase | rs7840728 | 0.1% |
| | | | | Nerve - Tibial | Increase | rs4870988 | 3.2% |
| | | | | Skin - Sun Exposed | Increase | rs13263296 | 88.7% |
| | | | | Thyroid | Increase | rs73703111 | 0.1% |
| | | *TAF2* | GTEx | Colon - Transverse | Increase | rs112349158 | 87.5% |
| 20 | rs41308092_A | *LIME1* | GTEx | Muscle - Skeletal | Increase | rs4809330 | 20.5% |

[a] Only results for significantly associated variants in the NESDA-NTR dataset were available, therefore the colocalisation analysis was run only including variants significantly associated with gene expression in blood rather than all variants in the region.
[b] The lung eQTL dataset showed two independent signals of association for *DEPTOR* expression. The eQTL results here are those obtained after conditioning on the top eQTL for *DEPTOR* to condition out the strongest signal which was driven by different variants to those driving the IPF risk association.

**<u>Supplementary Table 3 - Annotation and eQTL results for variants in 95% credible sets of novel IPF susceptibility signals</u>**

##### i) Chromosome 3

| rsid | chr | Position | GWAS *P* | Posterior Probability | Annotation | $R^2$ with sentinel | eQTL | | | |
|---|---|---|---|---|---|---|---|---|---|---|
| | | | | | | | Lung eQTL | GTEx (lung) | GTEx (non-lung tissue) | NESDA-NTR |
| rs78238620 | 3 | 44902386 | $5.12\times10^{-10}$ | 14.71% | intron (*KIF15*) | Sentinel | - | - | *KIF15, TMEM42, KIAA1143* | - |
| rs2292180 | 3 | 44903349 | $5.42\times10^{-10}$ | 14.01% | intron (*KIF15*) | 1.00 | - | - | *KIF15, TMEM42, KIAA1143* | - |
| rs2292181 | 3 | 44903434 | $5.42\times10^{-10}$ | 14.01% | synonymous (*TMEM42*), intron (*KIF15*), non-coding exon (*MIR564*) | 1.00 | - | - | *KIF15, TMEM42, KIAA1143* | - |
| rs74341405 | 3 | 44845649 | $7.74\times10^{-10}$ | 8.90% | intron (*KIF15*) | 0.89 | - | - | *KIF15, TMEM42* | - |
| rs80059929 | 3 | 44846722 | $7.74\times10^{-10}$ | 8.90% | intron (*KIF15*) | 0.89 | - | - | *KIF15, TMEM42* | - |
| rs76304484 | 3 | 44877209 | $2.20\times10^{-9}$ | 4.24% | intron (*KIF15*) | 0.99 | - | - | *KIF15, TMEM42, KIAA1143* | - |
| rs6792918 | 3 | 44857004 | $2.83\times10^{-9}$ | 3.43% | intron (*KIF15*) | 0.99 | - | - | *KIF15, TMEM42, KIAA1143* | - |
| rs77568017 | 3 | 44877853 | $3.57\times10^{-9}$ | 2.79% | intron (*KIF15*) | 0.99 | | - | *KIF15, TMEM42, KIAA1143* | - |
| rs76526953 | 3 | 44881909 | $3.57\times10^{-9}$ | 2.79% | intron (*KIF15*) | 0.99 | - | - | *KIF15, TMEM42, KIAA1143* | - |
| rs141979279 | 3 | 44858131 | $4.01\times10^{-9}$ | 2.52% | intron (*KIF15*) | 0.99 | - | - | *KIF15, TMEM42, KIAA1143* | - |
| rs55661644 | 3 | 44869509 | $4.01\times10^{-9}$ | 2.52% | intron (*KIF15*) | 0.99 | - | - | *KIF15, TMEM42, KIAA1143* | - |
| rs7340559 | 3 | 44871986 | $4.01\times10^{-9}$ | 2.52% | intron (*KIF15*) | 0.99 | - | - | *KIF15, TMEM42, KIAA1143* | - |
| rs77136835 | 3 | 44874693 | $4.01\times10^{-9}$ | 2.52% | intron (*KIF15*) | 0.99 | - | - | *KIF15, TMEM42, KIAA1143* | - |
| rs112645395 | 3 | 44794881 | $3.97\times10^{-9}$ | 1.92% | missense (*KIAA1143*)<br><br>[SIFT: Deleterious<br>PolyPhen: Possibly damaging<br>CADD: Likely benign<br>REVEL: Likely benign<br>MetaLR: Tolerated<br>MutationAssessor: Medium] | 0.79 | - | - | *TMEM42* | - |
| rs149000267 | 3 | 44836326 | $3.09\times10^{-9}$ | 1.56% | intron (*KIF15*) | 0.64 | - | - | *KIF15, TMEM42* | - |
| rs77938604 | 3 | 44836543 | $7.81\times10^{-9}$ | 1.28% | intron (*KIF15*) | 0.90 | - | - | *KIF15, KIAA1143* | - |
| rs4682996 | 3 | 44819436 | $1.05\times10^{-8}$ | 0.99% | intron (*KIF15*) | 0.89 | - | - | *KIAA1143, TMEM42* | - |
| rs4682992 | 3 | 44786946 | $1.22\times10^{-8}$ | 0.88% | intron (*KIAA1143*) | 0.89 | - | - | *KIAA1143, TMEM42* | - |
| rs112842175 | 3 | 44788306 | $1.22\times10^{-8}$ | 0.88% | intron (*KIAA1143*) | 0.89 | - | - | *KIAA1143, TMEM42* | - |

| | | | | | | | | | | |
|---|---|---|---|---|---|---|---|---|---|---|
| rs4682993 | 3 | 44795238 | $1.22\times10^{-8}$ | 0.88% | intron (*KIAA1143*) | 0.89 | - | - | *KIAA1143, TMEM42* | - |
| rs77805183 | 3 | 44797277 | $1.22\times10^{-8}$ | 0.88% | intron (*KIAA1143*) | 0.89 | - | - | *KIAA1143, TMEM42* | - |
| rs111788055 | 3 | 44833973 | $1.36\times10^{-8}$ | 0.81% | intron (*KIF15*) | 0.90 | - | - | *KIF15, KIAA1143* | - |
| rs79850585 | 3 | 44756245 | $1.38\times10^{-8}$ | 0.80% | intron (*ZNF502*) | 0.88 | - | - | *KIAA1143, TMEM42* | - |
| rs4682994 | 3 | 44803130 | $1.58\times10^{-8}$ | 0.71% | intron (*KIF15*) | 0.89 | - | - | *KIF15, TMEM42, KIAA1143* | - |

**ii) Chromosome 7**

| rsid | Chr | Position | GWAS *P* | Posterior Probability | Annotation | $R^2$ with Sentinel | eQTL | | | |
|---|---|---|---|---|---|---|---|---|---|---|
| | | | | | | | Lung eQTL | GTEx (lung) | GTEx (non-lung tissue) | NESDA-NTR |
| rs12699415 | 7 | 1909479 | $7.15\times10^{-13}$ | 35.67% | intron (*MAD1L1*) | Sentinel | - | - | *MAD1L1* | - |
| rs7795126 | 7 | 2076626 | $8.47\times10^{-12}$ | 3.38% | intron (*MAD1L1*) | 0.71 | - | - | *MAD1L1* | - |
| rs10950503 | 7 | 2039594 | $1.10\times10^{-11}$ | 2.64% | intron (*MAD1L1*) | 0.80 | - | - | *MAD1L1* | - |
| rs4455739 | 7 | 1864356 | $1.12\times10^{-11}$ | 2.61% | intron (*MAD1L1*) | 0.68 | - | - | *AC110781.3* | - |
| rs34373690 | 7 | 1869473 | $1.27\times10^{-11}$ | 2.31% | intron (*MAD1L1*) | 0.68 | - | - | *AC110781.3* | - |
| rs34074471 | 7 | 1865249 | $1.27\times10^{-11}$ | 2.30% | intron (*MAD1L1*) | 0.68 | - | - | *AC110781.3* | - |
| rs35091011 | 7 | 1865174 | $1.33\times10^{-11}$ | 2.21% | intron (*MAD1L1*) | 0.68 | | - | *AC110781.3* | - |
| rs35406566 | 7 | 1865527 | $1.33\times10^{-11}$ | 2.21% | intron (*MAD1L1*) | 0.68 | - | - | *AC110781.3* | - |
| rs6974455 | 7 | 1865921 | $1.35\times10^{-11}$ | 2.17% | intron (*MAD1L1*) | 0.68 | - | - | *AC110781.3* | - |
| rs61164094 | 7 | 1874264 | $1.37\times10^{-11}$ | 2.14% | intron (*MAD1L1*) | 0.68 | - | - | *AC110781.3* | - |
| rs4255035 | 7 | 1864444 | $1.41\times10^{-11}$ | 2.09% | intron (*MAD1L1*) | 0.68 | - | - | *AC110781.3* | - |
| rs4379359 | 7 | 1864415 | $1.52\times10^{-11}$ | 1.94% | intron (*MAD1L1*) | 0.68 | - | - | *AC110781.3* | - |
| rs7806394 | 7 | 1864129 | $1.53\times10^{-11}$ | 1.93% | intron (*MAD1L1*) | 0.68 | - | - | *AC110781.3* | - |
| rs4631355 | 7 | 1864245 | $1.53\times10^{-11}$ | 1.93% | intron (*MAD1L1*) | 0.68 | - | - | *AC110781.3* | - |
| rs3857706 | 7 | 2034193 | $1.53\times10^{-11}$ | 1.93% | intron (*MAD1L1*) | 0.81 | - | - | *MAD1L1* | - |
| rs13225346 | 7 | 1866916 | $1.59\times10^{-11}$ | 1.86% | intron (*MAD1L1*) | 0.68 | - | - | *AC110781.3* | - |
| rs57193069 | 7 | 1862417 | $1.74\times10^{-11}$ | 1.71% | intron (*MAD1L1*) | 0.68 | - | - | *AC110781.3* | - |
| rs872464 | 7 | 2034562 | $1.93\times10^{-11}$ | 1.54% | intron (*MAD1L1*) | 0.81 | - | - | *MAD1L1* | - |
| rs6955652 | 7 | 1865583 | $1.98\times10^{-11}$ | 1.51% | intron (*MAD1L1*) | 0.68 | - | - | *AC110781.3* | - |
| rs7799807 | 7 | 1868092 | $2.01\times10^{-11}$ | 1.48% | intron (*MAD1L1*) | 0.37 | - | - | *AC110781.3, FTSJ2* | - |
| rs35641411 | 7 | 1870242 | $2.03\times10^{-11}$ | 1.48% | intron (*MAD1L1*) | 0.68 | - | - | *AC110781.3* | - |
| rs1403174 | 7 | 2032865 | $2.03\times10^{-11}$ | 1.47% | intron (*MAD1L1*) | 0.82 | - | - | *MAD1L1* | - |
| rs28661143 | 7 | 1866395 | $2.24\times10^{-11}$ | 1.34% | intron (*MAD1L1*) | 0.68 | - | - | *AC110781.3* | - |
| rs12537430 | 7 | 1868761 | $3.25\times10^{-11}$ | 0.93% | intron (*MAD1L1*) | 0.36 | - | - | *AC110781.3, FTSJ2* | - |
| rs12537479 | 7 | 1868995 | $3.48\times10^{-11}$ | 0.88% | intron (*MAD1L1*) | 0.36 | - | - | *AC110781.3, FTSJ2* | - |
| rs35935754 | 7 | 1869242 | $3.60\times10^{-11}$ | 0.85% | intron (*MAD1L1*) | 0.36 | - | - | *AC110781.3, FTSJ2* | - |

| | | | | | | | | | | |
|---|---|---|---|---|---|---|---|---|---|---|
| rs7799782 | 7 | 1868039 | $4.05\times10^{-11}$ | 0.76% | intron (*MAD1L1*) | 0.36 | - | - | *AC110781.3, FTSJ2* | - |
| rs6959688 | 7 | 1966831 | $4.33\times10^{-11}$ | 0.72% | intron (*MAD1L1*) | 0.86 | - | - | *MAD1L1* | - |
| rs56053419 | 7 | 1863463 | $5.32\times10^{-11}$ | 0.59% | intron (*MAD1L1*) | 0.36 | - | - | *AC110781.3, FTSJ2* | - |
| rs55948146 | 7 | 1866953 | $8.08\times10^{-11}$ | 0.39% | intron (*MAD1L1*) | 0.36 | - | - | *AC110781.3, FTSJ2* | - |
| rs12672286 | 7 | 1907009 | $8.29\times10^{-11}$ | 0.38% | intron (*MAD1L1*) | 0.38 | - | - | *AC110781.3, FTSJ2* | - |
| rs6978112 | 7 | 1966841 | $9.52\times10^{-11}$ | 0.34% | intron (*MAD1L1*) | 0.84 | - | - | *MAD1L1* | - |
| rs4721090 | 7 | 1873084 | $1.03\times10^{-10}$ | 0.31% | intron (*MAD1L1*) | 0.32 | - | - | *AC110781.3, FTSJ2* | - |
| rs11761670 | 7 | 1904709 | $1.12\times10^{-10}$ | 0.29% | intron (*MAD1L1*) | 0.37 | - | - | *AC110781.3, FTSJ2* | - |
| rs13224015 | 7 | 1913869 | $1.11\times10^{-10}$ | 0.29% | intron (*MAD1L1*) | 0.84 | - | - | *MAD1L1* | *MAD1L1* |
| rs4721143 | 7 | 1918179 | $1.14\times10^{-10}$ | 0.28% | intron (*MAD1L1*) | 0.84 | - | - | *MAD1L1* | *MAD1L1* |
| rs34120092 | 7 | 1861952 | $1.25\times10^{-10}$ | 0.26% | intron (*MAD1L1*) | 0.35 | - | - | *AC110781.3, FTSJ2* | - |
| rs7786367 | 7 | 1863828 | $1.27\times10^{-10}$ | 0.26% | intron (*MAD1L1*) | 0.71 | - | - | *AC110781.3* | - |
| rs13235380 | 7 | 1873894 | $1.27\times10^{-10}$ | 0.26% | intron (*MAD1L1*) | 0.32 | - | - | *AC110781.3, FTSJ2* | - |
| rs9770241 | 7 | 1863164 | $1.37\times10^{-10}$ | 0.24% | intron (*MAD1L1*) | 0.37 | - | - | *AC110781.3, FTSJ2* | - |
| rs10807751 | 7 | 1883476 | $1.39\times10^{-10}$ | 0.24% | intron (*MAD1L1*), intron (*AC110781*.3) | 0.70 | - | - | *AC110781.3* | - |
| rs13221208 | 7 | 1913856 | $1.35\times10^{-10}$ | 0.24% | intron (*MAD1L1*) | 0.84 | - | - | *MAD1L1* | *MAD1L1* |
| rs34256344 | 7 | 1861460 | $1.44\times10^{-10}$ | 0.23% | intron (*MAD1L1*) | 0.72 | - | - | *AC110781.3* | - |
| rs10237989 | 7 | 1873343 | $1.42\times10^{-10}$ | 0.23% | intron (*MAD1L1*) | 0.32 | - | - | *AC110781.3, FTSJ2* | - |
| rs13222183 | 7 | 1873879 | $1.44\times10^{-10}$ | 0.23% | intron (*MAD1L1*) | 0.32 | - | - | *AC110781.3, FTSJ2* | - |
| rs6460944 | 7 | 1876199 | $1.45\times10^{-10}$ | 0.23% | intron (*MAD1L1*) | 0.58 | - | - | *AC110781.3* | *MAD1L1* |
| rs12537387 | 7 | 1868582 | $1.64\times10^{-10}$ | 0.20% | intron (*MAD1L1*) | 0.57 | - | - | - | *MAD1L1* |
| rs4719319 | 7 | 1888094 | $1.68\times10^{-10}$ | 0.20% | intron (*MAD1L1*) | 0.33 | - | - | *AC110781.3, FTSJ2* | - |
| rs4719330 | 7 | 1914613 | $1.65\times10^{-10}$ | 0.20% | intron (*MAD1L1*) | 0.85 | - | - | *MAD1L1* | *MAD1L1* |
| rs4721139 | 7 | 1917337 | $1.64\times10^{-10}$ | 0.20% | intron (*MAD1L1*) | 0.84 | - | - | *MAD1L1* | *MAD1L1* |
| rs6949794 | 7 | 1908727 | $1.83\times10^{-10}$ | 0.18% | intron (*MAD1L1*) | 0.39 | - | - | *MAD1L1, AC110781.3, FTSJ2* | - |
| rs10950400 | 7 | 1882470 | $1.99\times10^{-10}$ | 0.17% | intron (*MAD1L1*), intron (*AC110781*.3) | 0.33 | - | - | *AC110781.3, FTSJ2* | - |
| rs4639400 | 7 | 1917806 | $1.93\times10^{-10}$ | 0.17% | intron (*MAD1L1*) | 0.84 | - | - | *MAD1L1* | *MAD1L1* |

| | | | | | | | | | | |
|---|---|---|---|---|---|---|---|---|---|---|
| rs7783715 | 7 | 1923385 | $2.00\times10^{-10}$ | 0.17% | intron (*MAD1L1*) | 0.85 | - | - | - | *MAD1L1* |
| rs6977733 | 7 | 1886725 | $2.05\times10^{-10}$ | 0.16% | intron (*MAD1L1*), intron (*AC110781*.3) | 0.33 | - | - | *AC110781.3, FTSJ2* | - |
| rs6954521 | 7 | 1886865 | $2.13\times10^{-10}$ | 0.16% | intron (*MAD1L1*), intron (*AC110781*.3) | 0.33 | - | - | *AC110781.3, FTSJ2* | - |
| rs6978048 | 7 | 1886872 | $2.13\times10^{-10}$ | 0.16% | intron (*MAD1L1*), intron (*AC110781*.3) | 0.33 | - | - | *AC110781.3, FTSJ2* | - |
| rs6954673 | 7 | 1886937 | $2.05\times10^{-10}$ | 0.16% | intron (*MAD1L1*), missense (*AC110781*.3) [PolyPhen: Benign CADD: Likely benign] | 0.33 | - | - | *AC110781.3, FTSJ2* | - |
| rs4610628 | 7 | 1903100 | $2.08\times10^{-10}$ | 0.16% | intron (*MAD1L1*) | 0.36 | - | - | *AC110781.3, FTSJ2* | - |
| rs10950411 | 7 | 1909153 | $2.13\times10^{-10}$ | 0.16% | intron (*MAD1L1*) | 0.85 | - | - | *MAD1L1* | *MAD1L1* |
| rs4458759 | 7 | 1876081 | $2.23\times10^{-10}$ | 0.15% | intron (*MAD1L1*) | 0.58 | - | - | *AC110781.3* | *MAD1L1* |
| rs6948403 | 7 | 1876768 | $2.28\times10^{-10}$ | 0.15% | intron (*MAD1L1*) | 0.58 | - | - | *AC110781.3* | *MAD1L1* |
| rs6953693 | 7 | 1886388 | $2.25\times10^{-10}$ | 0.15% | intron (*MAD1L1*), intron (*AC110781*.3) | 0.33 | - | - | *AC110781.3, FTSJ2* | - |
| rs10950410 | 7 | 1909086 | $2.20\times10^{-10}$ | 0.15% | intron (*MAD1L1*) | 0.73 | - | - | *AC110781.3* | *MAD1L1* |
| rs57216949 | 7 | 2030287 | $2.30\times10^{-10}$ | 0.15% | intron (*MAD1L1*) | 0.37 | - | - | *MAD1L1, FTSJ2* | - |
| rs12534763 | 7 | 1868711 | $2.45\times10^{-10}$ | 0.14% | intron (*MAD1L1*) | 0.58 | - | - | - | *MAD1L1* |
| rs6965935 | 7 | 1876895 | $2.44\times10^{-10}$ | 0.14% | intron (*MAD1L1*) | 0.58 | - | - | *AC110781.3* | *MAD1L1* |
| rs6957894 | 7 | 1887362 | $2.38\times10^{-10}$ | 0.14% | intron (*MAD1L1*), missense (*AC110781*.3) [PolyPhen: Probably damaging CADD: Likely benign] | 0.33 | - | - | *AC110781.3, FTSJ2* | - |
| rs4719318 | 7 | 1887930 | $2.38\times10^{-10}$ | 0.14% | intron (*MAD1L1*) | 0.33 | - | - | *AC110781.3, FTSJ2* | - |
| rs35349665 | 7 | 1911166 | $2.48\times10^{-10}$ | 0.14% | intron (*MAD1L1*) | 0.84 | - | - | - | *MAD1L1* |
| rs4721134 | 7 | 1912057 | $2.35\times10^{-10}$ | 0.14% | intron (*MAD1L1*) | 0.39 | - | - | *AC110781.3, FTSJ2* | - |
| rs4449693 | 7 | 1884630 | $2.53\times10^{-10}$ | 0.13% | intron (*MAD1L1*), intron (*AC110781*.3) | 0.33 | - | - | *AC110781.3, FTSJ2* | - |
| rs6952808 | 7 | 1886535 | $2.66\times10^{-10}$ | 0.13% | intron (*MAD1L1*), intron (*AC110781*.3) | 0.33 | - | - | *AC110781.3, FTSJ2* | - |

| | | | | | | | | | | |
|---|---|---|---|---|---|---|---|---|---|---|
| rs10260585 | 7 | 1889521 | $2.54\times10^{-10}$ | 0.13% | intron (*MAD1L1*) | 0.33 | - | - | *AC110781.3, FTSJ2* | - |
| rs4256490 | 7 | 1890764 | $2.63\times10^{-10}$ | 0.13% | intron (*MAD1L1*) | 0.33 | - | - | *AC110781.3, FTSJ2* | - |
| rs4601204 | 7 | 1890925 | $2.51\times10^{-10}$ | 0.13% | intron (*MAD1L1*) | 0.33 | - | - | *AC110781.3, FTSJ2* | - |
| rs6948707 | 7 | 1870794 | $2.89\times10^{-10}$ | 0.12% | intron (*MAD1L1*) | 0.58 | - | - | - | *MAD1L1* |
| rs3889797 | 7 | 1877924 | $2.91\times10^{-10}$ | 0.12% | intron (*MAD1L1*) | 0.58 | - | - | *AC110781.3* | *MAD1L1* |
| rs4721122 | 7 | 1893311 | $2.74\times10^{-10}$ | 0.12% | intron (*MAD1L1*) | 0.33 | - | - | *AC110781.3, FTSJ2* | - |
| rs12155225 | 7 | 1899479 | $2.91\times10^{-10}$ | 0.12% | intron (*MAD1L1*) | 0.38 | - | - | *MAD1L1, AC110781.3, FTSJ2* | - |
| rs12538674 | 7 | 1925166 | $2.79\times10^{-10}$ | 0.12% | intron (*MAD1L1*) | 0.85 | - | - | *MAD1L1* | *MAD1L1* |
| rs4721287 | 7 | 2028663 | $2.75\times10^{-10}$ | 0.12% | intron (*MAD1L1*) | 0.37 | - | - | *MAD1L1, FTSJ2* | - |
| rs56727870 | 7 | 2029940 | $2.98\times10^{-10}$ | 0.11% | intron (*MAD1L1*) | 0.37 | - | - | *MAD1L1, FTSJ2* | - |
| rs60995052 | 7 | 2030007 | $3.03\times10^{-10}$ | 0.11% | intron (*MAD1L1*) | 0.37 | - | - | *MAD1L1, FTSJ2* | - |
| rs60755037 | 7 | 2030104 | $3.03\times10^{-10}$ | 0.11% | intron (*MAD1L1*) | 0.37 | - | - | *MAD1L1, FTSJ2* | - |
| 7:2036550 | 7 | 2036550 | $2.97\times10^{-10}$ | 0.11% | intron (*MAD1L1*) | 0.37 | - | - | - | - |

**iii) Chromosome 8**

| rsid | Chr | Position | GWAS *P* | Posterior Probability | Annotation | $R^2$ with Sentinel | eQTL | | | |
|---|---|---|---|---|---|---|---|---|---|---|
| | | | | | | | Lung eQTL | GTEx (lung) | GTEx (non-lung tissue) | NESDA-NTR |
| rs28513081 | 8 | 120934126 | $1.20\times10^{-9}$ | 4.51% | intron (*DEPTOR*) | Sentinel | *DEPTOR* | *DEPTOR, RP11-760H22.2* | *DEPTOR, DSCC1, KB-1471A8.1, RP11-760H22.2, TAF2* | - |
| rs6469878 | 8 | 120938448 | $1.66\times10^{-9}$ | 3.33% | intron (*DEPTOR*) | 0.93 | *DEPTOR* | *DEPTOR, RP11-760H22.2* | *DEPTOR, DSCC1, KB-1471A8.1, RP11-760H22.2, TAF2* | *DEPTOR* |
| rs7814294 | 8 | 120939436 | $1.71\times10^{-9}$ | 3.23% | intron (*DEPTOR*) | 0.93 | *DEPTOR* | *DEPTOR, RP11-760H22.2* | *DEPTOR, DSCC1, KB-1471A8.1, RP11-760H22.2, TAF2* | *DEPTOR* |
| rs2037346 | 8 | 120935452 | $1.73\times10^{-9}$ | 3.19% | intron (*DEPTOR*) | 0.93 | *DEPTOR* | *DEPTOR, RP11-760H22.2* | *DEPTOR, DSCC1, KB-1471A8.1, RP11-760H22.2, TAF2* | *DEPTOR* |
| rs10808505 | 8 | 120940206 | $1.85\times10^{-9}$ | 3.00% | intron (*DEPTOR*) | 0.93 | *DEPTOR* | *DEPTOR, RP11-760H22.2* | *DEPTOR, DSCC1, KB-1471A8.1, RP11-760H22.2, TAF2* | *DEPTOR* |
| rs4871787 | 8 | 120936418 | $2.08\times10^{-9}$ | 2.68% | intron (*DEPTOR*) | 0.93 | *DEPTOR* | *DEPTOR, RP11-760H22.2* | *DEPTOR, DSCC1, KB-1471A8.1, RP11-760H22.2, TAF2* | *DEPTOR* |
| rs1464276 | 8 | 120937041 | $2.21\times10^{-9}$ | 2.53% | intron (*DEPTOR*) | 0.93 | *DEPTOR* | *DEPTOR, RP11-760H22.2* | *DEPTOR, DSCC1, KB-1471A8.1, RP11-760H22.2, TAF2* | *DEPTOR* |
| rs10107579 | 8 | 120934569 | $2.62\times10^{-9}$ | 2.16% | intron (*DEPTOR*) | 0.93 | *DEPTOR* | *DEPTOR, RP11-760H22.2* | *DEPTOR, DSCC1, KB-1471A8.1, RP11-760H22.2, TAF2* | *DEPTOR* |
| rs56864850 | 8 | 120943444 | $2.74\times10^{-9}$ | 2.07% | intron (*DEPTOR*) | 0.93 | *DEPTOR* | *DEPTOR, RP11-760H22.2* | *DEPTOR, DSCC1, KB-1471A8.1, RP11-760H22.2, TAF2* | *DEPTOR* |
| rs55892034 | 8 | 120943507 | $2.74\times10^{-9}$ | 2.07% | intron (*DEPTOR*) | 0.93 | *DEPTOR* | *DEPTOR, RP11-760H22.2* | *DEPTOR, DSCC1, KB-1471A8.1, RP11-760H22.2, TAF2* | *DEPTOR* |
| rs9987332 | 8 | 120933963 | $4.18\times10^{-9}$ | 1.39% | intron (*DEPTOR*) | 0.96 | *DEPTOR, DSCC1* | *DEPTOR, RP11-760H22.2* | *DEPTOR, DSCC1, KB-1471A8.1, RP11-760H22.2, TAF2* | - |
| rs13265546 | 8 | 120919975 | $5.01\times10^{-9}$ | 1.17% | intron (*DEPTOR*) | 0.93 | *DEPTOR* | *DEPTOR, RP11-760H22.2* | *DEPTOR, DSCC1, KB-1471A8.1, RP11-760H22.2, TAF2* | *DEPTOR* |
| rs35006524 | 8 | 120930769 | $5.28\times10^{-9}$ | 1.11% | intron (*DEPTOR*) | 0.93 | *DEPTOR* | *DEPTOR, RP11-760H22.2* | *DEPTOR, DSCC1, KB-1471A8.1, RP11-760H22.2, TAF2* | - |
| rs1607624 | 8 | 120929289 | $5.90\times10^{-9}$ | 1.00% | intron (*DEPTOR*) | 0.93 | *DEPTOR* | *DEPTOR, RP11-760H22.2* | *DEPTOR, DSCC1, KB-1471A8.1, RP11-760H22.2, TAF2* | *DEPTOR* |
| rs7829901 | 8 | 120929834 | $5.90\times10^{-9}$ | 1.00% | intron (*DEPTOR*) | 0.93 | *DEPTOR* | *DEPTOR, RP11-760H22.2* | *DEPTOR, DSCC1, KB-1471A8.1, RP11-760H22.2, TAF2* | - |
| rs13267896 | 8 | 120920654 | $5.91\times10^{-9}$ | 1.00% | intron (*DEPTOR*) | 0.93 | *DEPTOR* | *DEPTOR, RP11-760H22.2* | *DEPTOR, DSCC1, KB-1471A8.1, RP11-760H22.2, TAF2* | *DEPTOR* |

| | | | | | | | | | | |
|---|---|---|---|---|---|---|---|---|---|---|
| rs13275524 | 8 | 120920941 | $5.91\times10^{-9}$ | 1.00% | intron (*DEPTOR*) | 0.93 | *DEPTOR* | *DEPTOR, RP11-760H22.2* | *DEPTOR, DSCC1, KB-1471A8.1, RP11-760H22.2, TAF2* | - |
| rs6469867 | 8 | 120921412 | $5.91\times10^{-9}$ | 1.00% | intron (*DEPTOR*) | 0.93 | *DEPTOR* | *DEPTOR, RP11-760H22.2* | *DEPTOR, DSCC1, KB-1471A8.1, RP11-760H22.2, TAF2* | *DEPTOR* |
| rs6469868 | 8 | 120921841 | $5.91\times10^{-9}$ | 1.00% | intron (*DEPTOR*) | 0.93 | *DEPTOR* | *DEPTOR, RP11-760H22.2* | *DEPTOR, DSCC1, KB-1471A8.1, RP11-760H22.2, TAF2* | *DEPTOR* |
| rs6469871 | 8 | 120922079 | $5.91\times10^{-9}$ | 1.00% | intron (*DEPTOR*) | 0.93 | *DEPTOR* | *DEPTOR, RP11-760H22.2* | *DEPTOR, DSCC1, KB-1471A8.1, RP11-760H22.2, TAF2* | - |
| rs6469872 | 8 | 120922247 | $5.91\times10^{-9}$ | 1.00% | intron (*DEPTOR*) | 0.93 | *DEPTOR* | *DEPTOR, RP11-760H22.2* | *DEPTOR, DSCC1, KB-1471A8.1, RP11-760H22.2, TAF2* | - |
| rs7015470 | 8 | 120922341 | $5.91\times10^{-9}$ | 1.00% | intron (*DEPTOR*) | 0.93 | *DEPTOR* | *DEPTOR, RP11-760H22.2* | *DEPTOR, DSCC1, KB-1471A8.1, RP11-760H22.2, TAF2* | - |
| rs6995139 | 8 | 120922397 | $5.91\times10^{-9}$ | 1.00% | intron (*DEPTOR*) | 0.93 | *DEPTOR* | *DEPTOR, RP11-760H22.2* | *DEPTOR, DSCC1, KB-1471A8.1, RP11-760H22.2, TAF2* | - |
| rs10217077 | 8 | 120923183 | $5.91\times10^{-9}$ | 1.00% | intron (*DEPTOR*) | 0.93 | *DEPTOR* | *DEPTOR, RP11-760H22.2* | *DEPTOR, DSCC1, KB-1471A8.1, RP11-760H22.2, TAF2* | - |
| rs10217083 | 8 | 120923285 | $5.91\times10^{-9}$ | 1.00% | intron (*DEPTOR*) | 0.93 | *DEPTOR* | *DEPTOR, RP11-760H22.2* | *DEPTOR, DSCC1, KB-1471A8.1, RP11-760H22.2, TAF2* | *DEPTOR* |
| rs939242 | 8 | 120924270 | $5.91\times10^{-9}$ | 1.00% | intron (*DEPTOR*) | 0.93 | *DEPTOR* | *DEPTOR, RP11-760H22.2* | *DEPTOR, DSCC1, KB-1471A8.1, RP11-760H22.2, TAF2* | - |
| rs939241 | 8 | 120924537 | $5.91\times10^{-9}$ | 1.00% | intron (*DEPTOR*) | 0.93 | *DEPTOR* | *DEPTOR, RP11-760H22.2* | *DEPTOR, DSCC1, KB-1471A8.1, RP11-760H22.2, TAF2* | - |
| rs10216503 | 8 | 120925083 | $5.91\times10^{-9}$ | 1.00% | intron (*DEPTOR*) | 0.93 | *DEPTOR* | *DEPTOR, RP11-760H22.2* | *DEPTOR, DSCC1, KB-1471A8.1, RP11-760H22.2, TAF2* | - |
| rs11785871 | 8 | 120925199 | $5.91\times10^{-9}$ | 1.00% | intron (*DEPTOR*) | 0.93 | *DEPTOR, DSCC1* | *DEPTOR, RP11-760H22.2* | *DEPTOR, DSCC1, KB-1471A8.1, RP11-760H22.2, TAF2* | - |
| rs7824545 | 8 | 120925621 | $5.91\times10^{-9}$ | 1.00% | intron (*DEPTOR*) | 0.93 | *DEPTOR* | *DEPTOR, RP11-760H22.2* | *DEPTOR, DSCC1, KB-1471A8.1, RP11-760H22.2, TAF2* | - |
| rs7387264 | 8 | 120925998 | $5.91\times10^{-9}$ | 1.00% | intron (*DEPTOR*) | 0.93 | *DEPTOR* | *DEPTOR, RP11-760H22.2* | *DEPTOR, DSCC1, KB-1471A8.1, RP11-760H22.2, TAF2* | - |
| rs7388508 | 8 | 120926153 | $5.91\times10^{-9}$ | 1.00% | intron (*DEPTOR*) | 0.93 | *DEPTOR* | *DEPTOR, RP11-760H22.2* | *DEPTOR, DSCC1, KB-1471A8.1, RP11-760H22.2, TAF2* | - |
| rs7462302 | 8 | 120917771 | $6.15\times10^{-9}$ | 0.96% | intron (*DEPTOR*) | 0.93 | *DEPTOR* | *DEPTOR, RP11-760H22.2* | *DEPTOR, DSCC1, KB-1471A8.1, RP11-760H22.2, TAF2* | *DEPTOR* |
| rs12545863 | 8 | 120918111 | $6.15\times10^{-9}$ | 0.96% | intron (*DEPTOR*) | 0.93 | *DEPTOR* | *DEPTOR, RP11-760H22.2* | *DEPTOR, DSCC1, KB-1471A8.1, RP11-760H22.2, TAF2* | - |

| | | | | | | | | | | |
|---|---|---|---|---|---|---|---|---|---|---|
| rs7002839 | 8 | 120918748 | $6.15\times10^{-9}$ | 0.96% | intron (*DEPTOR*) | 0.93 | *DEPTOR* | *DEPTOR, RP11-760H22.2* | *DEPTOR, DSCC1, KB-1471A8.1, RP11-760H22.2, TAF2* | *DEPTOR* |
| rs7006905 | 8 | 120918923 | $6.15\times10^{-9}$ | 0.96% | intron (*DEPTOR*) | 0.93 | *DEPTOR* | *DEPTOR, RP11-760H22.2* | *DEPTOR, DSCC1, KB-1471A8.1, RP11-760H22.2, TAF2* | *DEPTOR* |
| rs9720591 | 8 | 120916840 | $6.36\times10^{-9}$ | 0.93% | intron (*DEPTOR*) | 0.93 | *DEPTOR* | *DEPTOR, RP11-760H22.2* | *DEPTOR, DSCC1, KB-1471A8.1, RP11-760H22.2, TAF2* | *DEPTOR* |
| rs4073560 | 8 | 120917388 | $6.36\times10^{-9}$ | 0.93% | intron (*DEPTOR*) | 0.93 | *DEPTOR* | *DEPTOR, RP11-760H22.2* | *DEPTOR, DSCC1, KB-1471A8.1, RP11-760H22.2, TAF2* | *DEPTOR* |
| rs10110216 | 8 | 120887547 | $7.01\times10^{-9}$ | 0.85% | intron (*DEPTOR*) | 0.84 | *DEPTOR, DSCC1* | *DEPTOR, RP11-760H22.2* | *DEPTOR, DSCC1, KB-1471A8.1, RP11-760H22.2, TAF2* | *DEPTOR* |
| rs10110223 | 8 | 120887566 | $7.05\times10^{-9}$ | 0.84% | intron (*DEPTOR*) | 0.84 | *DEPTOR, DSCC1* | *DEPTOR, RP11-760H22.2* | *DEPTOR, DSCC1, KB-1471A8.1, RP11-760H22.2, TAF2* | *DEPTOR* |
| rs13257252 | 8 | 120912233 | $7.13\times10^{-9}$ | 0.84% | intron (*DEPTOR*) | 0.92 | *DEPTOR, DSCC1* | *DEPTOR, RP11-760H22.2* | *DEPTOR, DSCC1, KB-1471A8.1, RP11-760H22.2, TAF2* | *DEPTOR* |
| rs35272074 | 8 | 120910783 | $7.39\times10^{-9}$ | 0.81% | intron (*DEPTOR*) | 0.92 | *DEPTOR, DSCC1* | *DEPTOR, RP11-760H22.2* | *DEPTOR, DSCC1, KB-1471A8.1, RP11-760H22.2, TAF2* | *DEPTOR* |
| rs10103660 | 8 | 120910787 | $7.39\times10^{-9}$ | 0.81% | intron (*DEPTOR*) | 0.92 | *DEPTOR, DSCC1* | *DEPTOR, RP11-760H22.2* | *DEPTOR, DSCC1, KB-1471A8.1, RP11-760H22.2, TAF2* | *DEPTOR* |
| rs13281299 | 8 | 120909493 | $7.40\times10^{-9}$ | 0.81% | intron (*DEPTOR*) | 0.92 | *DEPTOR, DSCC1* | *DEPTOR, RP11-760H22.2* | *DEPTOR, DSCC1, KB-1471A8.1, RP11-760H22.2, TAF2* | *DEPTOR* |
| rs7814496 | 8 | 120909628 | $7.40\times10^{-9}$ | 0.81% | intron (*DEPTOR*) | 0.92 | *DEPTOR, DSCC1* | *DEPTOR, RP11-760H22.2* | *DEPTOR, DSCC1, KB-1471A8.1, RP11-760H22.2, TAF2* | *DEPTOR* |
| rs7814520 | 8 | 120909665 | $7.40\times10^{-9}$ | 0.81% | intron (*DEPTOR*) | 0.92 | *DEPTOR* | *DEPTOR, RP11-760H22.2* | *DEPTOR, DSCC1, KB-1471A8.1, RP11-760H22.2, TAF2* | *DEPTOR* |
| rs7832923 | 8 | 120909780 | $7.40\times10^{-9}$ | 0.81% | intron (*DEPTOR*) | 0.92 | *DEPTOR, DSCC1* | *DEPTOR, RP11-760H22.2* | *DEPTOR, DSCC1, KB-1471A8.1, RP11-760H22.2, TAF2* | *DEPTOR* |
| rs6469861 | 8 | 120909913 | $7.40\times10^{-9}$ | 0.81% | intron (*DEPTOR*) | 0.92 | *DEPTOR, DSCC1* | *DEPTOR, RP11-760H22.2* | *DEPTOR, DSCC1, KB-1471A8.1, RP11-760H22.2, TAF2* | *DEPTOR* |
| rs6469862 | 8 | 120910036 | $7.40\times10^{-9}$ | 0.81% | intron (*DEPTOR*) | 0.92 | *DEPTOR, DSCC1* | *DEPTOR, RP11-760H22.2* | *DEPTOR, DSCC1, KB-1471A8.1, RP11-760H22.2, TAF2* | *DEPTOR* |
| rs6469863 | 8 | 120910150 | $7.40\times10^{-9}$ | 0.81% | intron (*DEPTOR*) | 0.92 | *DEPTOR, DSCC1* | *DEPTOR, RP11-760H22.2* | *DEPTOR, DSCC1, KB-1471A8.1, RP11-760H22.2, TAF2* | *DEPTOR* |
| rs6469864 | 8 | 120910225 | $7.40\times10^{-9}$ | 0.81% | intron (*DEPTOR*) | 0.92 | *DEPTOR, DSCC1* | *DEPTOR, RP11-760H22.2* | *DEPTOR, DSCC1, KB-1471A8.1, RP11-760H22.2, TAF2* | *DEPTOR* |
| rs6469865 | 8 | 120910235 | $7.40\times10^{-9}$ | 0.81% | intron (*DEPTOR*) | 0.92 | *DEPTOR* | *DEPTOR, RP11-760H22.2* | *DEPTOR, DSCC1, KB-1471A8.1, RP11-760H22.2, TAF2* | *DEPTOR* |

| | | | | | | | | | | |
|---|---|---|---|---|---|---|---|---|---|---|
| rs6987580 | 8 | 120910380 | $7.40\times10^{-9}$ | 0.81% | intron (*DEPTOR*) | 0.92 | *DEPTOR, DSCC1* | *DEPTOR, RP11-760H22.2* | *DEPTOR, DSCC1, KB-1471A8.1, RP11-760H22.2, TAF2* | *DEPTOR* |
| rs6988011 | 8 | 120910538 | $7.40\times10^{-9}$ | 0.81% | intron (*DEPTOR*) | 0.92 | *DEPTOR, DSCC1* | *DEPTOR, RP11-760H22.2* | *DEPTOR, DSCC1, KB-1471A8.1, RP11-760H22.2, TAF2* | *DEPTOR* |
| rs6993375 | 8 | 120911630 | $7.40\times10^{-9}$ | 0.81% | intron (*DEPTOR*) | 0.92 | *DEPTOR* | *DEPTOR, RP11-760H22.2* | *DEPTOR, DSCC1, KB-1471A8.1, RP11-760H22.2, TAF2* | *DEPTOR* |
| rs6993797 | 8 | 120911645 | $7.40\times10^{-9}$ | 0.81% | intron (*DEPTOR*) | 0.92 | *DEPTOR* | *DEPTOR, RP11-760H22.2* | *DEPTOR, DSCC1, KB-1471A8.1, RP11-760H22.2, TAF2* | *DEPTOR* |
| rs6989741 | 8 | 120912154 | $7.40\times10^{-9}$ | 0.81% | intron (*DEPTOR*) | 0.92 | *DEPTOR, DSCC1* | *DEPTOR, RP11-760H22.2* | *DEPTOR, DSCC1, KB-1471A8.1, RP11-760H22.2, TAF2* | *DEPTOR* |
| rs13258592 | 8 | 120912429 | $7.40\times10^{-9}$ | 0.81% | intron (*DEPTOR*) | 0.92 | *DEPTOR, DSCC1* | *DEPTOR, RP11-760H22.2* | *DEPTOR, DSCC1, KB-1471A8.1, RP11-760H22.2, TAF2* | *DEPTOR* |
| rs10093230 | 8 | 120912846 | $7.40\times10^{-9}$ | 0.81% | intron (*DEPTOR*) | 0.92 | *DEPTOR, DSCC1* | *DEPTOR, RP11-760H22.2* | *DEPTOR, DSCC1, KB-1471A8.1, RP11-760H22.2, TAF2* | *DEPTOR* |
| rs9649947 | 8 | 120913357 | $7.40\times10^{-9}$ | 0.81% | intron (*DEPTOR*) | 0.92 | *DEPTOR, DSCC1* | *DEPTOR, RP11-760H22.2* | *DEPTOR, DSCC1, KB-1471A8.1, RP11-760H22.2, TAF2* | *DEPTOR* |
| rs7465609 | 8 | 120913860 | $7.40\times10^{-9}$ | 0.81% | intron (*DEPTOR*) | 0.92 | *DEPTOR, DSCC1* | *DEPTOR, RP11-760H22.2* | *DEPTOR, DSCC1, KB-1471A8.1, RP11-760H22.2, TAF2* | - |
| rs7465612 | 8 | 120913885 | $7.40\times10^{-9}$ | 0.81% | intron (*DEPTOR*) | 0.92 | *DEPTOR, DSCC1* | *DEPTOR, RP11-760H22.2* | *DEPTOR, DSCC1, KB-1471A8.1, RP11-760H22.2, TAF2* | - |
| rs13279398 | 8 | 120914270 | $7.40\times10^{-9}$ | 0.81% | intron (*DEPTOR*) | 0.92 | *DEPTOR, DSCC1* | *DEPTOR, RP11-760H22.2* | *DEPTOR, DSCC1, KB-1471A8.1, RP11-760H22.2, TAF2* | *DEPTOR* |
| rs13277947 | 8 | 120914432 | $7.40\times10^{-9}$ | 0.81% | intron (*DEPTOR*) | 0.92 | *DEPTOR, DSCC1* | *DEPTOR, RP11-760H22.2* | *DEPTOR, DSCC1, KB-1471A8.1, RP11-760H22.2, TAF2* | *DEPTOR* |
| rs13277992 | 8 | 120914527 | $7.40\times10^{-9}$ | 0.81% | intron (*DEPTOR*) | 0.92 | *DEPTOR, DSCC1* | *DEPTOR, RP11-760H22.2* | *DEPTOR, DSCC1, KB-1471A8.1, RP11-760H22.2, TAF2* | *DEPTOR* |
| rs13253140 | 8 | 120914908 | $7.40\times10^{-9}$ | 0.81% | intron (*DEPTOR*) | 0.92 | *DEPTOR, DSCC1* | *DEPTOR, RP11-760H22.2* | *DEPTOR, DSCC1, KB-1471A8.1, RP11-760H22.2, TAF2* | *DEPTOR* |
| rs11777705 | 8 | 120915232 | $7.40\times10^{-9}$ | 0.81% | intron (*DEPTOR*) | 0.92 | *DEPTOR* | *DEPTOR, RP11-760H22.2* | *DEPTOR, DSCC1, KB-1471A8.1, RP11-760H22.2, TAF2* | *DEPTOR* |
| rs12681402 | 8 | 120915862 | $7.40\times10^{-9}$ | 0.81% | intron (*DEPTOR*) | 0.92 | *DEPTOR, DSCC1* | *DEPTOR, RP11-760H22.2* | *DEPTOR, DSCC1, KB-1471A8.1, RP11-760H22.2, TAF2* | *DEPTOR* |
| rs7461290 | 8 | 120916394 | $7.40\times10^{-9}$ | 0.81% | intron (*DEPTOR*) | 0.92 | *DEPTOR, DSCC1* | *DEPTOR, RP11-760H22.2* | *DEPTOR, DSCC1, KB-1471A8.1, RP11-760H22.2, TAF2* | *DEPTOR* |
| rs9721029 | 8 | 120916555 | $7.40\times10^{-9}$ | 0.81% | intron (DEPTOR) | 0.92 | *DEPTOR, DSCC1* | *DEPTOR, RP11-760H22.2* | *DEPTOR, DSCC1, KB-1471A8.1, RP11-760H22.2, TAF2* | *DEPTOR* |

| | | | | | | | | | | |
|---|---|---|---|---|---|---|---|---|---|---|
| rs9721042 | 8 | 120916651 | $7.40{\times}10^{-9}$ | 0.81% | intron (*DEPTOR*) | 0.92 | *DEPTOR* | *DEPTOR, RP11-760H22.2* | *DEPTOR, DSCC1, KB-1471A8.1, RP11-760H22.2, TAF2* | *DEPTOR* |
| rs10098306 | 8 | 120908921 | $7.50{\times}10^{-9}$ | 0.80% | intron (*DEPTOR*) | 0.92 | *DEPTOR, DSCC1* | *DEPTOR, RP11-760H22.2* | *DEPTOR, DSCC1, KB-1471A8.1, RP11-760H22.2, TAF2* | *DEPTOR* |
| rs7459671 | 8 | 120908886 | $8.66{\times}10^{-9}$ | 0.70% | intron (*DEPTOR*) | 0.92 | *DEPTOR, DSCC1* | *DEPTOR, RP11-760H22.2* | *DEPTOR, DSCC1, KB-1471A8.1, RP11-760H22.2, TAF2* | *DEPTOR* |
| rs7465181 | 8 | 120908589 | $8.72{\times}10^{-9}$ | 0.69% | intron (*DEPTOR*) | 0.92 | *DEPTOR, DSCC1* | *DEPTOR, RP11-760H22.2* | *DEPTOR, DSCC1, KB-1471A8.1, RP11-760H22.2, TAF2* | *DEPTOR* |
| rs7462250 | 8 | 120908607 | $9.14{\times}10^{-9}$ | 0.66% | intron (*DEPTOR*) | 0.92 | *DEPTOR, DSCC1* | *DEPTOR, RP11-760H22.2* | *DEPTOR, DSCC1, KB-1471A8.1, RP11-760H22.2, TAF2* | *DEPTOR* |
| rs1467044 | 8 | 120887041 | $9.71{\times}10^{-9}$ | 0.62% | intron (*DEPTOR*) | 0.84 | *DEPTOR, DSCC1* | *DEPTOR, RP11-760H22.2* | *DEPTOR, DSCC1, KB-1471A8.1, RP11-760H22.2, TAF2* | *DEPTOR* |
| rs7003126 | 8 | 120888752 | $9.71{\times}10^{-9}$ | 0.62% | intron (*DEPTOR*) | 0.84 | *DEPTOR, DSCC1* | *DEPTOR, RP11-760H22.2* | *DEPTOR, DSCC1, KB-1471A8.1, RP11-760H22.2, TAF2* | *DEPTOR* |
| rs11781657 | 8 | 120889242 | $9.71{\times}10^{-9}$ | 0.62% | intron (*DEPTOR*) | 0.84 | *DEPTOR, DSCC1* | *DEPTOR, RP11-760H22.2* | *DEPTOR, DSCC1, KB-1471A8.1, RP11-760H22.2, TAF2* | *DEPTOR* |
| rs4871013 | 8 | 120907990 | $9.73{\times}10^{-9}$ | 0.62% | intron (*DEPTOR*) | 0.92 | *DEPTOR, DSCC1* | *DEPTOR, RP11-760H22.2* | *DEPTOR, DSCC1, KB-1471A8.1, RP11-760H22.2, TAF2* | *DEPTOR* |
| rs4871772 | 8 | 120907895 | $1.21{\times}10^{-8}$ | 0.51% | intron (*DEPTOR*) | 0.92 | *DEPTOR, DSCC1* | *DEPTOR, RP11-760H22.2* | *DEPTOR, DSCC1, KB-1471A8.1, RP11-760H22.2, TAF2* | *DEPTOR* |
| rs4871012 | 8 | 120907974 | $1.21{\times}10^{-8}$ | 0.51% | intron (*DEPTOR*) | 0.92 | *DEPTOR, DSCC1* | *DEPTOR, RP11-760H22.2* | *DEPTOR, DSCC1, KB-1471A8.1, RP11-760H22.2, TAF2* | *DEPTOR* |
| rs4871773 | 8 | 120907911 | $1.21{\times}10^{-8}$ | 0.51% | intron (*DEPTOR*) | 0.92 | *DEPTOR, DSCC1* | *DEPTOR, RP11-760H22.2* | *DEPTOR, DSCC1, KB-1471A8.1, RP11-760H22.2, TAF2* | *DEPTOR* |
| rs13265367 | 8 | 120904009 | $1.74{\times}10^{-8}$ | 0.36% | intron (*DEPTOR*) | 0.90 | *DEPTOR, DSCC1* | *DEPTOR, RP11-760H22.2* | *DEPTOR, DSCC1, KB-1471A8.1, RP11-760H22.2, TAF2* | *DEPTOR* |
| rs10107251 | 8 | 120904348 | $1.74{\times}10^{-8}$ | 0.36% | intron (*DEPTOR*) | 0.90 | *DEPTOR* | *DEPTOR, RP11-760H22.2* | *DEPTOR, DSCC1, KB-1471A8.1, RP11-760H22.2, TAF2* | *DEPTOR* |
| rs10107361 | 8 | 120904427 | $1.74{\times}10^{-8}$ | 0.36% | intron (*DEPTOR*) | 0.90 | *DEPTOR, DSCC1* | *DEPTOR, RP11-760H22.2* | *DEPTOR, DSCC1, KB-1471A8.1, RP11-760H22.2, TAF2* | *DEPTOR* |
| rs10094455 | 8 | 120904676 | $1.74{\times}10^{-8}$ | 0.36% | intron (*DEPTOR*) | 0.90 | *DEPTOR* | *DEPTOR, RP11-760H22.2* | *DEPTOR, DSCC1, KB-1471A8.1, RP11-760H22.2, TAF2* | *DEPTOR* |
| rs10094458 | 8 | 120904688 | $1.74{\times}10^{-8}$ | 0.36% | intron (*DEPTOR*) | 0.90 | *DEPTOR, DSCC1* | *DEPTOR, RP11-760H22.2* | *DEPTOR, DSCC1, KB-1471A8.1, RP11-760H22.2, TAF2* | *DEPTOR* |
| rs10094587 | 8 | 120904800 | $1.74{\times}10^{-8}$ | 0.36% | intron (*DEPTOR*) | 0.90 | *DEPTOR, DSCC1* | *DEPTOR, RP11-760H22.2* | *DEPTOR, DSCC1, KB-1471A8.1, RP11-760H22.2, TAF2* | *DEPTOR* |

| | | | | | | | | | | |
|---|---|---|---|---|---|---|---|---|---|---|
| rs13275184 | 8 | 120905104 | $1.74\times10^{-8}$ | 0.36% | intron (*DEPTOR*) | 0.90 | *DEPTOR, DSCC1* | *DEPTOR, RP11-760H22.2* | *DEPTOR, DSCC1, KB-1471A8.1, RP11-760H22.2, TAF2* | - |
| rs13249122 | 8 | 120905919 | $1.74\times10^{-8}$ | 0.36% | intron (*DEPTOR*) | 0.90 | *DEPTOR, DSCC1* | *DEPTOR, RP11-760H22.2* | *DEPTOR, DSCC1, KB-1471A8.1, RP11-760H22.2, TAF2* | *DEPTOR* |
| rs13250594 | 8 | 120906391 | $1.74\times10^{-8}$ | 0.36% | intron (*DEPTOR*) | 0.90 | *DEPTOR, DSCC1* | *DEPTOR, RP11-760H22.2* | *DEPTOR, DSCC1, KB-1471A8.1, RP11-760H22.2, TAF2* | *DEPTOR* |
| rs13259990 | 8 | 120907183 | $1.75\times10^{-8}$ | 0.36% | intron (*DEPTOR*) | 0.90 | *DEPTOR, DSCC1* | *DEPTOR, RP11-760H22.2* | *DEPTOR, DSCC1, KB-1471A8.1, RP11-760H22.2, TAF2* | *DEPTOR* |
| rs13261304 | 8 | 120907420 | $1.79\times10^{-8}$ | 0.35% | intron (*DEPTOR*) | 0.90 | *DEPTOR, DSCC1* | *DEPTOR, RP11-760H22.2* | *DEPTOR, DSCC1, KB-1471A8.1, RP11-760H22.2, TAF2* | *DEPTOR* |
| rs12681623 | 8 | 120903856 | $1.81\times10^{-8}$ | 0.35% | intron (*DEPTOR*) | 0.90 | *DEPTOR, DSCC1* | *DEPTOR, RP11-760H22.2* | *DEPTOR, DSCC1, KB-1471A8.1, RP11-760H22.2, TAF2* | *DEPTOR* |
| rs13260933 | 8 | 120907250 | $1.86\times10^{-8}$ | 0.34% | intron (*DEPTOR*) | 0.90 | *DEPTOR, DSCC1* | *DEPTOR, RP11-760H22.2* | *DEPTOR, DSCC1, KB-1471A8.1, RP11-760H22.2, TAF2* | *DEPTOR* |

**iv) Chromosome 10**

| rsid | Chr | Position | GWAS *P* | Posterior Probability | Annotation | $R^2$ with sentinel | eQTL | | | |
|---|---|---|---|---|---|---|---|---|---|---|
| | | | | | | | Lung eQTL | GTEx (lung) | GTEx (non-lung tissue) | NESDA-NTR |
| rs537322302 | 10 | 93271016 | $3.43\times10^{-8}$ | 48.25% | intron (*HECTD2*) | Sentinel | - | - | - | - |
| rs547164341 | 10 | 93285553 | $3.41\times10^{-6}$ | 42.03% | intergenic | 0.77 | - | - | - | - |
| rs143984698 | 10 | 93059485 | 0.012 | 5.31% | intergenic | 0.20 | - | - | - | - |

**v) Chromosome 20**

| rsid | chr | Position | GWAS *P* | Posterior Probability | Annotation | $R^2$ with sentinel | eQTL | | | |
|---|---|---|---|---|---|---|---|---|---|---|
| | | | | | | | Lung eQTL | GTEx (lung) | GTEx (non-lung tissue) | NESDA-NTR |
| rs116905074 | 20 | 62377853 | $8.28\times10^{-10}$ | 33.16% | intron (*ZBTB46*) | 0.71 | - | - | - | - |
| rs41308092 | 20 | 62324391 | $7.65\times10^{-10}$ | 28.99% | intron (*RTEL1*, *RTEL1-TNFRSF6B*) | Sentinel | - | - | *LIME1* | - |
| rs118130858 | 20 | 62345681 | $1.28\times10^{-9}$ | 22.25% | intron (*ZGPAT*, *RP4-583P15.15*) | 0.81 | - | - | - | - |
| rs115610405 | 20 | 62325833 | $2.42\times10^{-9}$ | 14.20% | missense (*RTEL1*, *RTEL1-TNFRSF6B*)<br><br>[SIFT: Tolerated<br>PolyPhen: Possibly damaging<br>CADD: Likely benign<br>REVEL: Likely benign<br>MetaLR: Tolerated<br>MutationAssessor: Low] | 0.86 | - | - | - | - |

#### Supplementary Table 4 - SNPsea results for enrichment

##### i) Enrichment in differential gene expression between IPF cases and controls in four lung epithelial cell types

No variants showed enrichment in genes that are differentially expressed between 6 IPF cases and 3 healthy controls.

| Cell type | *P* value |
|---|---|
| Normal AT2 cells | 0.366 |
| Indeterminate cells | 0.485 |
| Basal cells | 0.236 |
| Club/Goblet | 0.475 |

##### ii) Enrichment of IPF risk loci in biological pathways

Below is a table of the 20 biological pathways with lowest p values in the SNPsea analysis. No pathways reached a Bonferroni corrected significance threshold for 1,751 tests of $P = 2.86 \times 10^{-5}$.

| Gene Ontology code | Pathway name | *P* value |
|---|---|---|
| GO:0000776 | Kinetochore | 0.001 |
| GO:0007059 | Chromosome segregation | 0.001 |
| GO:2001236 | - | 0.001 |
| GO:0000775 | Chromosome, centromeric region | 0.002 |
| GO:0000793 | Condensed chromosome | 0.002 |
| GO:0000779 | Condensed chromosome, centromeric region | 0.003 |
| GO:0044454 | Nuclear chromosome part | 0.003 |
| GO:0044427 | Chromosomal part | 0.004 |
| GO:0000819 | Sister chromatid segregation | 0.005 |
| GO:0002262 | Myeloid cell homeostasis | 0.005 |
| GO:0005198 | Structural molecule activity | 0.005 |
| GO:0006913 | Nucleocytoplasmic transport | 0.006 |
| GO:0051169 | Nuclear transport | 0.006 |
| GO:0000070 | Mitotic sister chromatid segregation | 0.007 |
| GO:0005085 | Guanyl-nucleotide exchange factor activity | 0.007 |
| GO:0022403 | Cell cycle phase | 0.007 |
| GO:0060589 | Nucleoside-triphosphatase regulator activity | 0.007 |
| GO:0000280 | Nuclear division | 0.008 |
| GO:0005819 | Spindle | 0.008 |
| GO:0000794 | Condensed nuclear chromosome | 0.009 |

#### Supplementary Table 5 - Results for IPF variants in interstitial lung abnormalities and lung function GWAS

This table shows the results from the interstitial lung abnormality (ILA) and lung function analyses for the 16 signals that reached genome-wide significance in the IPF case-control discovery meta-analysis. All results are presented for the allele that is associated with increased risk of IPF. The IPF odds ratios are the discovery meta-analysis results. $FEV_1$ is the forced expiratory volume after 1 second, FVC is the forced vital capacity and PEF is the peak expiratory flow. Variants that show an association after multiple testing corrections for 96 tests ($P < 0.00052$) are shown in green (variants that have $P < 0.05$ but do not reach significance are shown in yellow).

| Chr | rsid | Locus | Effect allele | EAF | IPF<br>OR<br>[95% CI]<br>*P* | Any ILA<br>OR<br>[95% CI]<br>*P* | Subpleural ILA<br>OR<br>[95% CI]<br>*P* | $FEV_1$<br>β<br>[95% CI]<br>*P* | FVC<br>β<br>[95% CI]<br>*P* | $FEV_1$ / FVC<br>β<br>[95% CI]<br>*P* | PEF<br>β<br>[95% CI]<br>*P* |
|---|---|---|---|---|---|---|---|---|---|---|---|
| 3 | rs78238620 | *KIF15* | A | 5.3% | 1.54<br>[1.38, 1.73]<br>$2.94\times10^{-14}$ | 1.15<br>[0.95, 1.40]<br>0.161 | 1.18<br>[0.95, 1.46]<br>0.126 | −0.011<br>[−0.022, 0.000]<br>0.069 | −0.022<br>[-0.033, 0.011]<br>$2.92\times10^{-4}$ | 0.017<br>[0.006, 0.028]<br>0.005 | 0.000<br>[−0.011, 0.011]<br>0.851 |
| 3 | rs12696304 | *LRRC34/TERC* | G | 27.90% | 1.31<br>[1.21, 1.40]<br>$7.09\times10^{-13}$ | 1.10<br>[0.99, 1.21]<br>0.073 | 1.18<br>[1.05, 1.31]<br>0.004 | −0.006<br>[−0.011, 0.000]<br>0.046 | −0.007<br>[−0.012, −0.001]<br>0.023 | 0.001<br>[−0.004, 0.007]<br>0.673 | 0.001<br>[−0.005, 0.006]<br>0.770 |
| 4 | rs2013701 | *FAM13A* | G | 51.30% | 1.28<br>[1.19, 1.35]<br>$3.30\times10^{-13}$ | 1.13<br>[1.03, 1.23]<br>0.009 | 1.11<br>[1.01, 1.23]<br>0.039 | 0.006<br>[0.001, 0.011]<br>0.013 | −0.014<br>[−0.019, −0.009]<br>$1.02\times10^{-7}$ | 0.043<br>[0.038, 0.048]<br>$3.56\times10^{-60}$ | 0.019<br>[0.014, 0.024]<br>$2.16\times10^{-13}$ |
| 5 | rs7725218 | *TERT* | G | 67.50% | 1.39<br>[1.30, 1.49]<br>$1.54\times10^{-20}$ | 1.05<br>[0.95, 1.16]<br>0.333 | 1.06<br>[0.95, 1.18]<br>0.310 | −0.005<br>[−0.010, 0.000]<br>0.042 | −0.007<br>[−0.012, −0.002]<br>0.007 | 0.004<br>[−0.001, 0.009]<br>0.154 | 0.001<br>[−0.004, 0.007]<br>0.526 |
| 6 | rs2076295 | *DSP* | G | 46.90% | 1.46<br>[1.37, 1.56]<br>$2.79\times10^{-30}$ | 1.18<br>[1.08, 1.29]<br>$2.84\times10^{-4}$ | 1.21<br>[1.10, 1.33]<br>$1.56\times10^{-4}$ | −0.002<br>[−0.007, 0.003]<br>0.432 | −0.013<br>[−0.018, −0.009]<br>$4.22\times10^{-7}$ | 0.023<br>[0.019, 0.028]<br>$2.52\times10^{-19}$ | 0.006<br>[0.001, 0.011]<br>0.020 |
| 7 | rs12699415 | *MAD1L1* | A | 42.0% | 1.28<br>[1.22, 1.35]<br>$5.50\times10^{-20}$ | 1.04<br>[0.95, 1.14]<br>0.438 | 0.98<br>[0.89, 1.09]<br>0.707 | −0.007<br>[−0.012, −0.002]<br>0.011 | −0.011<br>[−0.016, −0.007]<br>$1.41\times10^{-5}$ | 0.008<br>[0.003, 0.012]<br>0.005 | −0.001<br>[−0.006, 0.004]<br>0.581 |
| 7 | rs2897075 | 7q22.1 | T | 39.10% | 1.30<br>[1.21, 1.38]<br>$3.10\times10^{-14}$ | 1.03<br>[0.94, 1.13]<br>0.560 | 1.07<br>[0.97, 1.19]<br>0.165 | 0.007<br>[0.003, 0.012]<br>0.004 | −0.004<br>[-0.009, 0.000]<br>0.143 | 0.025<br>[0.020, 0.030]<br>$6.06\times10^{-21}$ | 0.012<br>[0.007, 0.017]<br>$2.86\times10^{-6}$ |
| 8 | rs28513081 | *DEPTOR* | A | 57.2% | 1.20<br>[1.14, 1.27]<br>$1.84\times10^{-11}$ | 1.03<br>[0.94, 1.12]<br>0.494 | 1.05<br>[0.95, 1.16]<br>0.331 | 0.001<br>[−0.004, 0.006]<br>0.822 | −0.005<br>[−0.010, −0.001]<br>0.045 | 0.011<br>[0.006, 0.016]<br>$4.22\times10^{-5}$ | 0.005<br>[0.000, 0.010]<br>0.073 |

| | | | | | | | | | | | |
|---|---|---|---|---|---|---|---|---|---|---|---|
| 10 | rs537322302 | *HECTD2* | G | 0.3% | 3.82<br>[2.25, 6.48]<br>$7.41\times10^{-7}$ | 3.25<br>[1.01, 10.5]<br>0.049 | 6.23<br>[1.87, 20.8]<br>0.003 | −0.005<br>[−0.042, 0.032]<br>0.832 | −0.014<br>[−0.051, 0.023]<br>0.524 | 0.019<br>[−0.018, 0.056]<br>0.386 | 0.021<br>[−0.016, 0.059]<br>0.312 |
| 11 | rs35705950 | *MUC5B* | T | 14.90% | 4.84<br>[4.37, 5.36]<br>$1.18\times10^{-203}$ | 1.98<br>[1.75, 2.24]<br>$4.45\times10^{-27}$ | 2.23<br>[1.94, 2.57]<br>$4.94\times10^{-29}$ | −0.005<br>[−0.013, 0.002]<br>0.147 | −0.013<br>[−0.021, −0.006]<br>$8.21\times10^{-4}$ | 0.015<br>[0.008, 0.023]<br>$7.11\times10^{-4}$ | 0.008<br>[0.000, 0.016]<br>0.063 |
| 13 | rs9577395 | *ATP11A* | C | 79.30% | 1.30<br>[1.20, 1.41]<br>$1.34\times10^{-10}$ | 1.07<br>[0.96, 1.19]<br>0.236 | 1.08<br>[0.96, 1.21]<br>0.225 | 0.011<br>[0.005, 0.017]<br>$2.82\times10^{-4}$ | 0.006<br>[0.000, 0.012]<br>0.041 | 0.012<br>[0.007, 0.018]<br>$2.04\times10^{-4}$ | 0.011<br>[0.005, 0.017]<br>$3.70\times10^{-4}$ |
| 15 | rs59424629 | *IVD* | G | 53.90% | 1.32<br>[1.22, 1.41]<br>$7.30\times10^{-16}$ | 1.11<br>[1.01, 1.20]<br>0.027 | 1.15<br>[1.04, 1.25]<br>0.005 | −0.014<br>[−0.019, −0.009]<br>$1.14\times10^{-7}$ | −0.015<br>[−0.020, 0.011]<br>$6.97\times10^{-9}$ | 0.000<br>[−0.005, 0.004]<br>0.851 | −0.005<br>[−0.000, 0.000]<br>0.072 |
| 15 | rs62023891 | *AKAP13* | A | 30.00% | 1.27<br>[1.18, 1.36]<br>$1.27\times10^{-10}$ | 1.14<br>[1.03, 1.25]<br>0.008 | 1.09<br>[0.98, 1.21]<br>0.124 | −0.001<br>[−0.006, 0.004]<br>0.726 | 0.001<br>[−0.005, 0.006]<br>0.834 | −0.003<br>[−0.008, 0.003]<br>0.356 | −0.004<br>[−0.010, 0.001]<br>0.109 |
| 17 | rs2077551 | *MAPT* | T | 81.40% | 1.41<br>[1.30, 1.54]<br>$2.83\times10^{-16}$ | 1.23<br>[1.07, 1.41]<br>0.003 | 1.23<br>[1.05, 1.43]<br>0.008 | 0.045<br>[0.039, 0.052]<br>$1.02\times10^{-42}$ | 0.044<br>[0.038, 0.050]<br>$3.21\times10^{-39}$ | 0.012<br>[0.006, 0.018]<br>$5.81\times10^{-4}$ | 0.025<br>[0.018, 0.031]<br>$2.04\times10^{-13}$ |
| 19 | rs12610495 | *DPP9* | G | 30.50% | 1.31<br>[1.22, 1.42]<br>$2.92\times10^{-12}$ | 1.12<br>[1.00, 1.25]<br>0.041 | 1.22<br>[1.08, 1.37]<br>0.001 | −0.001<br>[−0.006, 0.004]<br>0.682 | −0.004<br>[−0.009, 0.001]<br>0.127 | 0.004<br>[−0.001, 0.009]<br>0.153 | 0.005<br>[0.000, 0.011]<br>0.051 |
| 20 | rs41308092 | *RTEL1* | A | 2.1% | 1.83<br>[1.52, 2.20]<br>$1.38\times10^{-10}$ | 0.76<br>[0.48, 1.19]<br>0.232 | 0.87<br>[0.54, 1.40]<br>0.563 | −0.016<br>[−0.033, 0.001]<br>0.084 | −0.022<br>[−0.039, −0.004]<br>0.022 | 0.009<br>[−0.008, 0.027]<br>0.321 | −0.011<br>[−0.028, 0.007]<br>0.335 |

### Supplementary Figures

#### Supplementary Figure 1 - Study level QC

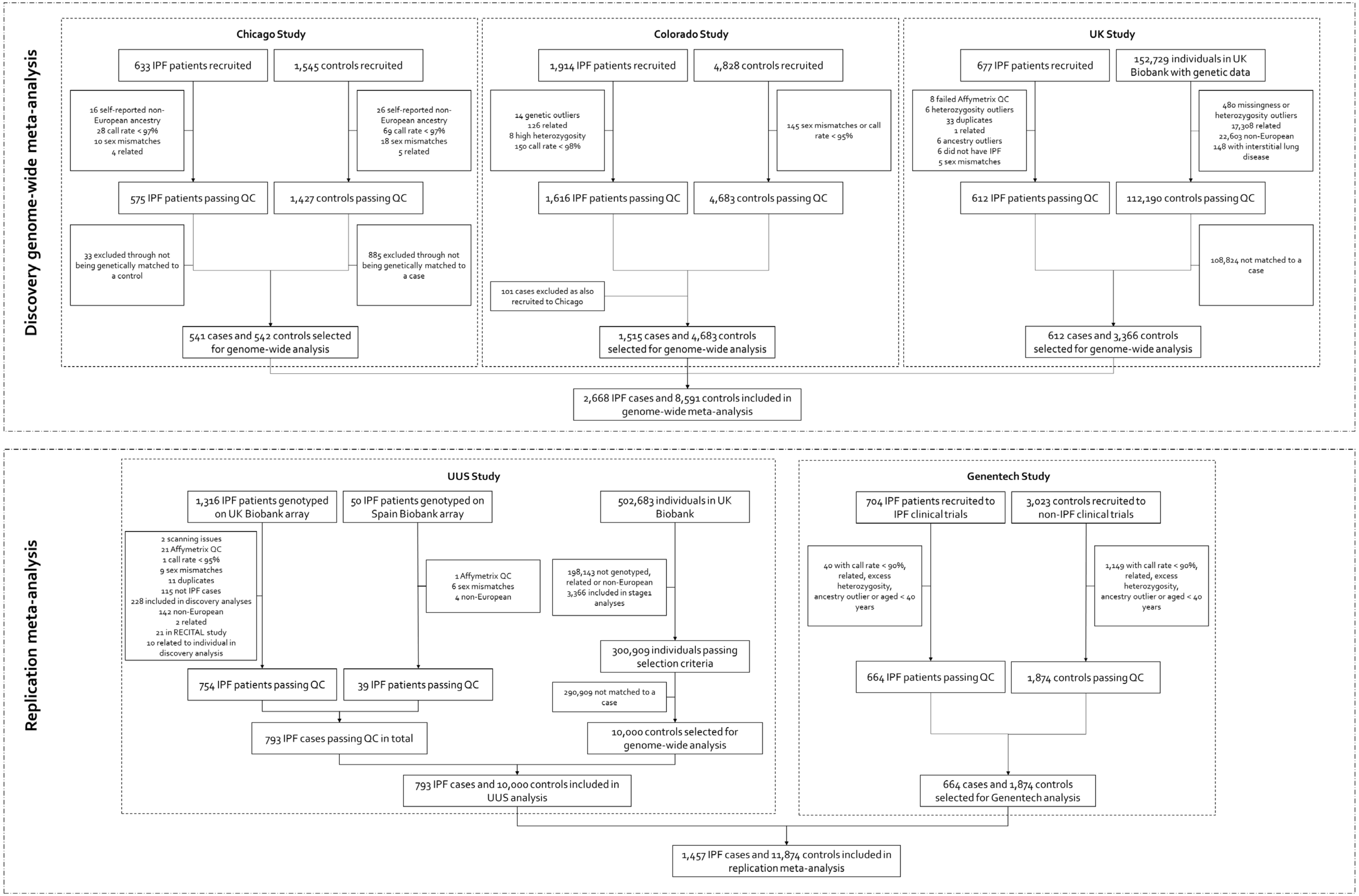

**Supplementary Figure 2 – Overlapping variants between discovery studies**
Venn diagram showing the number of variants in each study and the amount of overlapping variants between studies used for the discovery genome-wide meta-analysis.

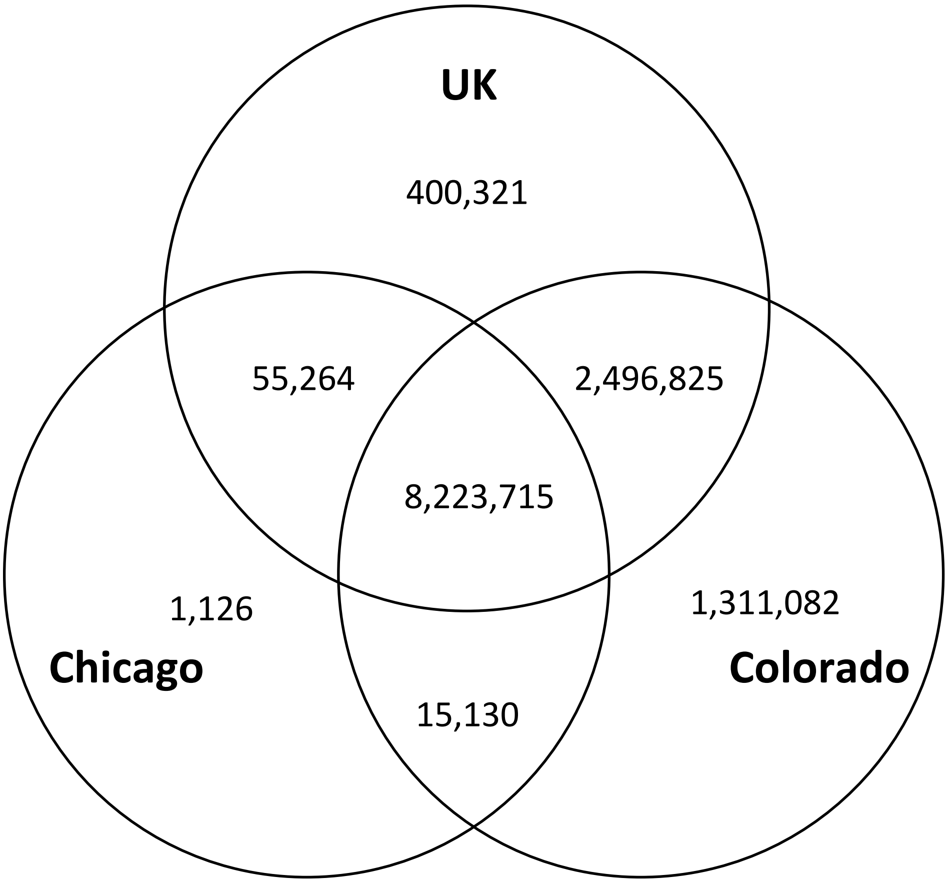

#### Supplementary Figure 3 - QQ plot for discovery genome-wide meta-analysis

QQ plot for the genome-wide analysis with expected −log(*P* value) on the x axis and observed −log(*P* value) on the y axis. The red line shows where the expected equals the observed.

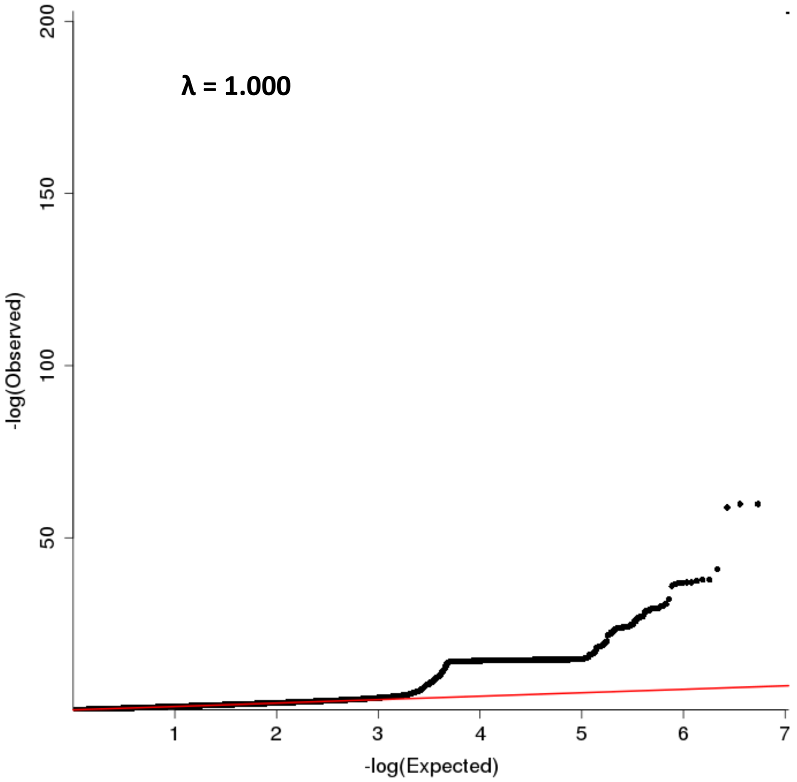

The QQ plot above shows an increasing line before plateuing with a large number of variants with the same p value. This is due to the inversion region around *KANSL1* and *MAPT* where a large number of variants in very high LD show similar strengths of association with IPF. Below is the QQ plot excluding this inversion region on chromosome 17.

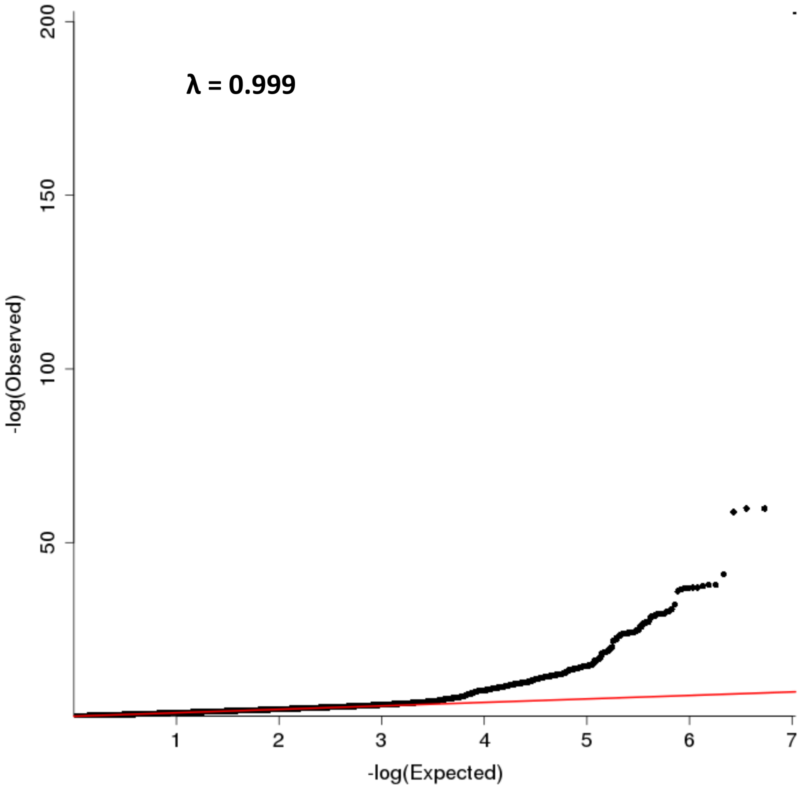

**Supplementary Figure 4 - Region plots for all 17 previously reported association signals**

Region plots for each of the 17 previously reported signals. The x axis shows chromosomal position and y axis the −log(*P* value). The sentinel (or previously reported variant if there is no signal in the meta-analysis) is shown in blue with all other variants coloured by LD with the sentinel variant. Credible sets were calculated for each signal and variants in the credible set are shown by squares. The red horizontal line shows $P = 5 \times 10^{-8}$.

**i) *TERC***

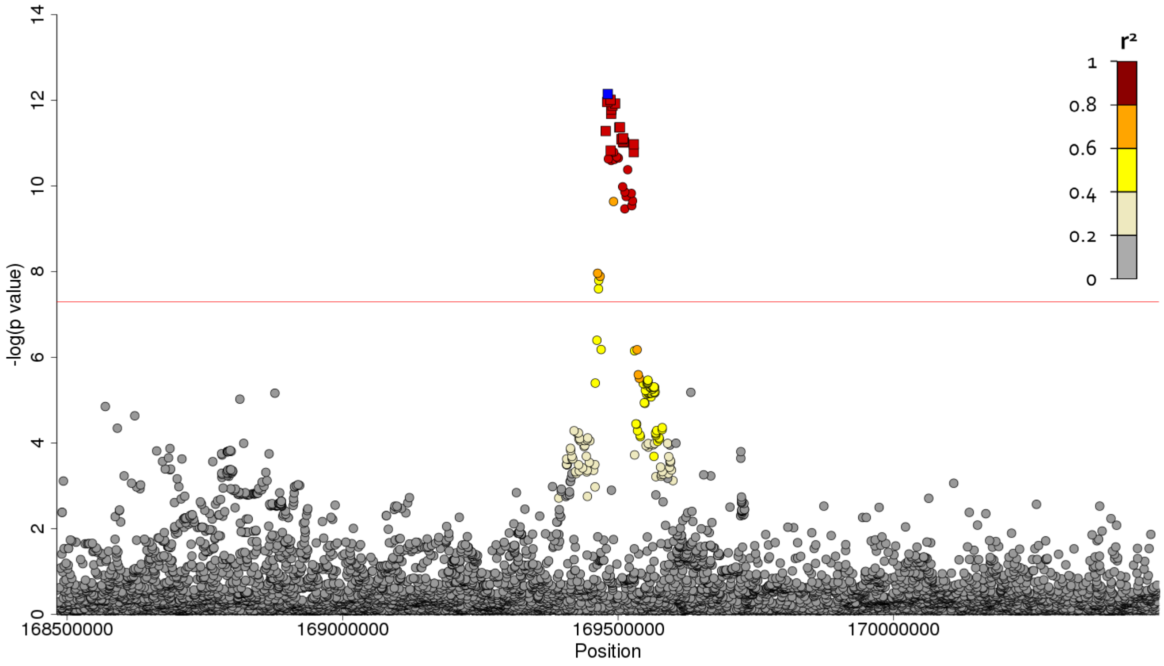

**ii) *FAM13A***

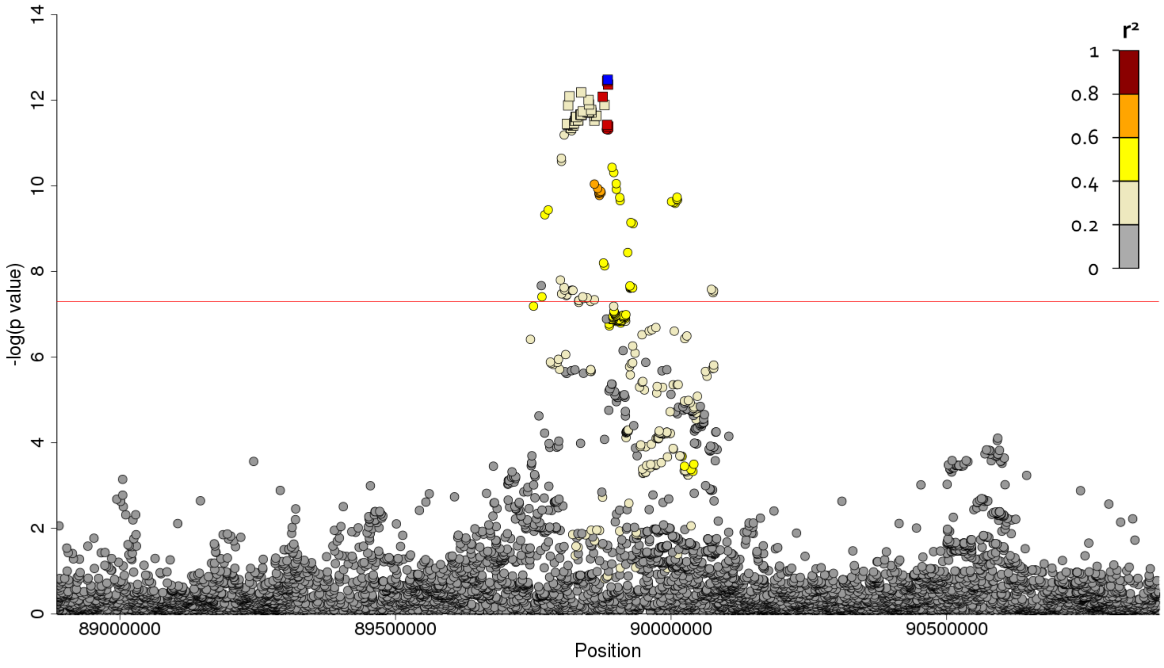

**iii) *TERT***

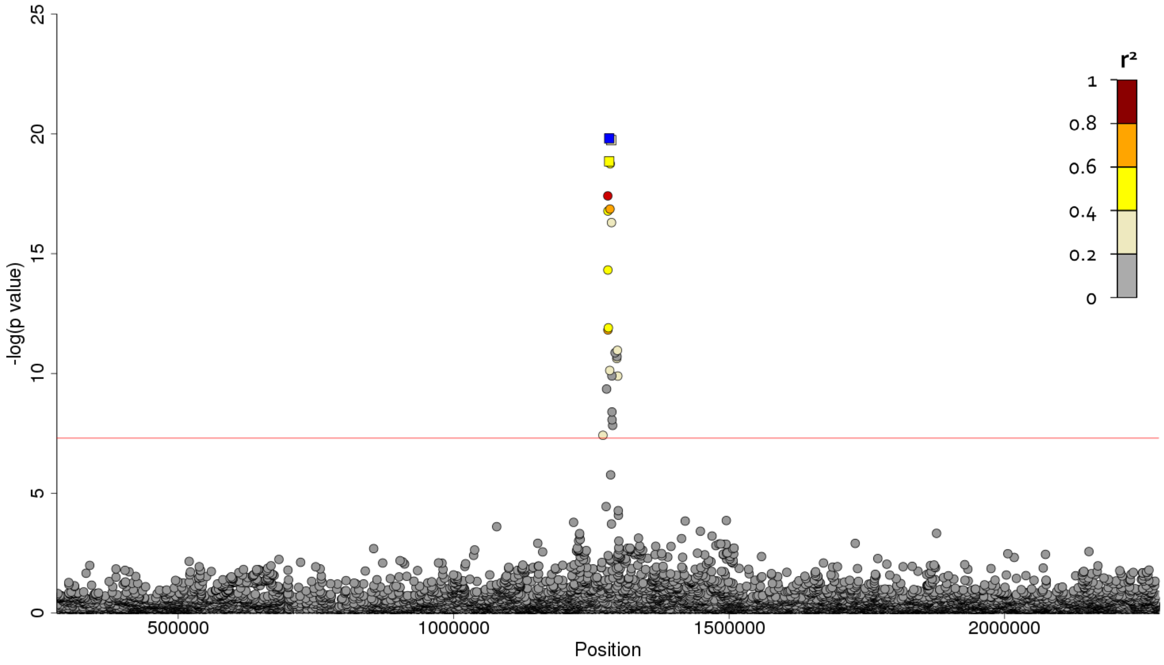

**iv) *DSP***

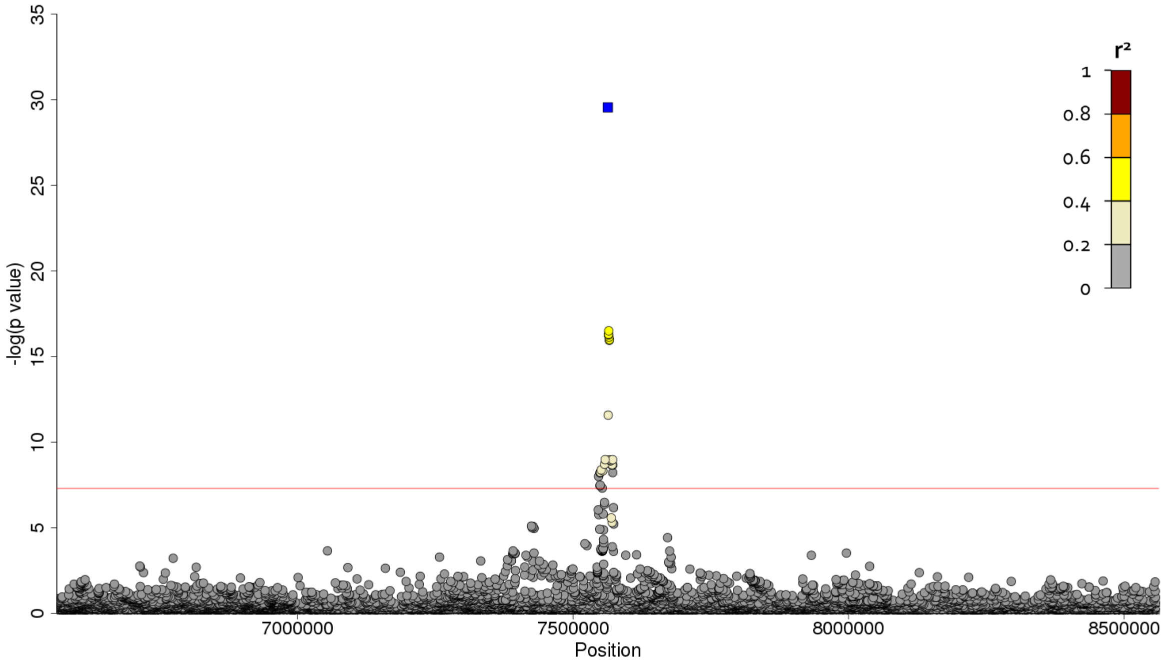

v) *EHMT2*

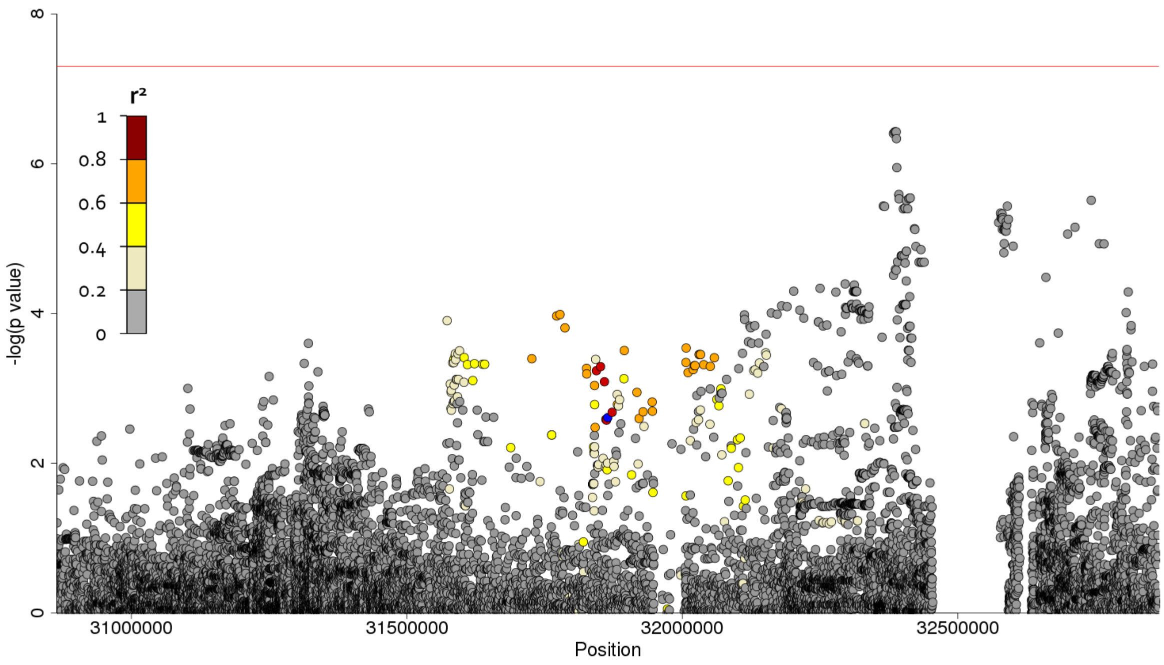

vi) 7q22.1

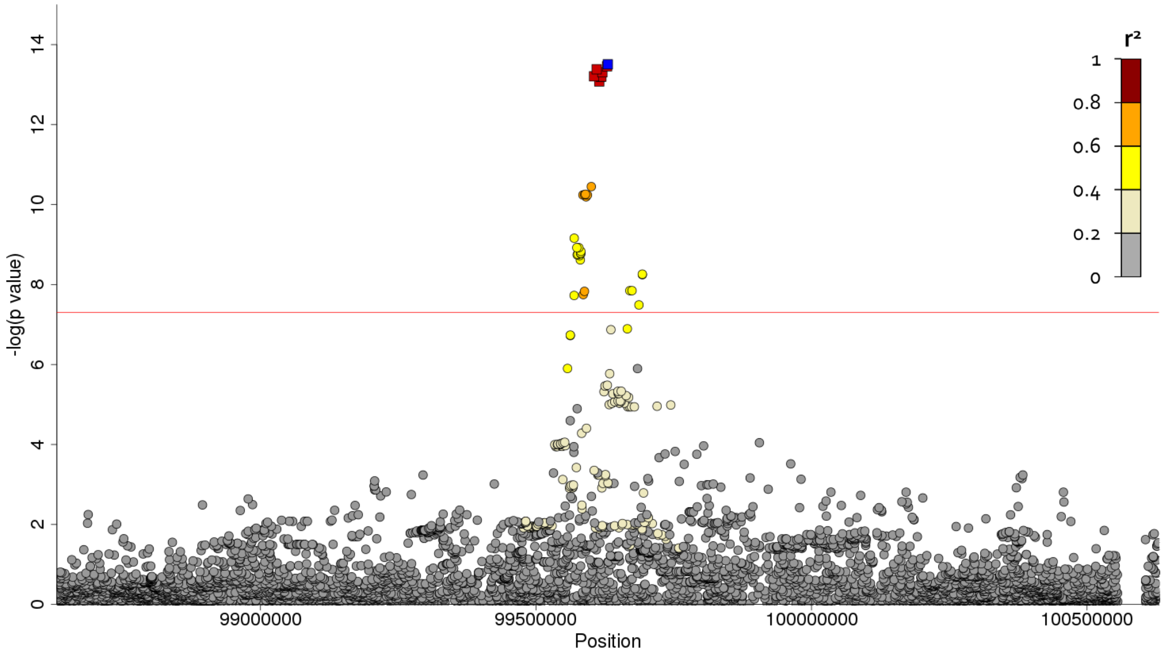

**vii)** ***OBFC1***

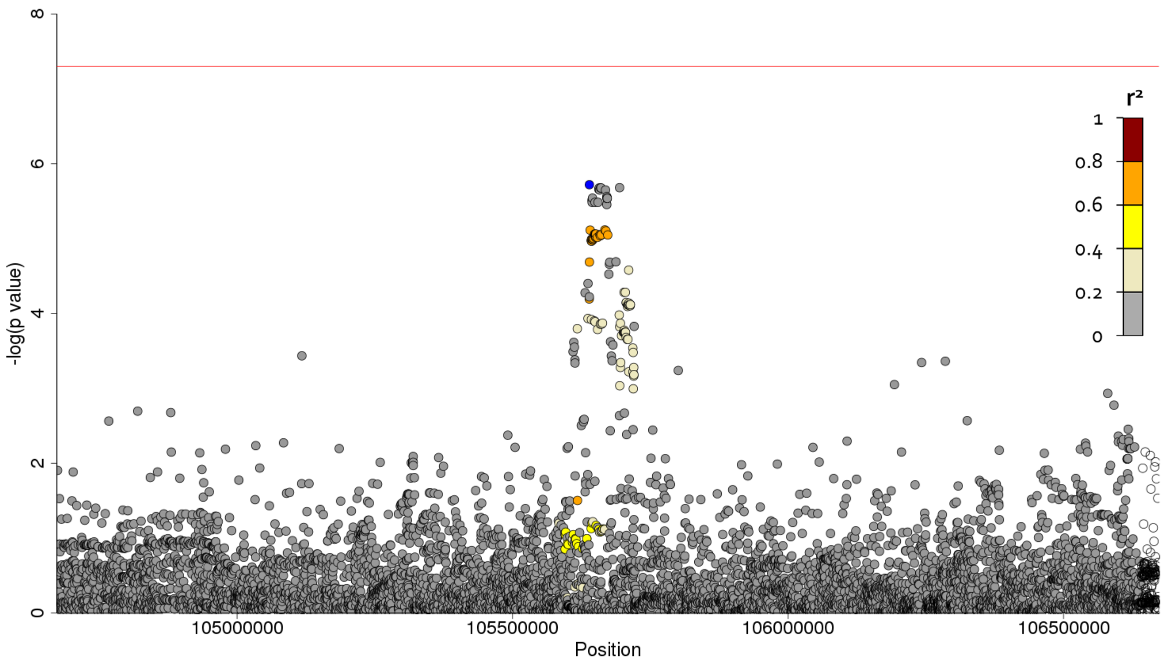

**viii)** ***MUC5B***

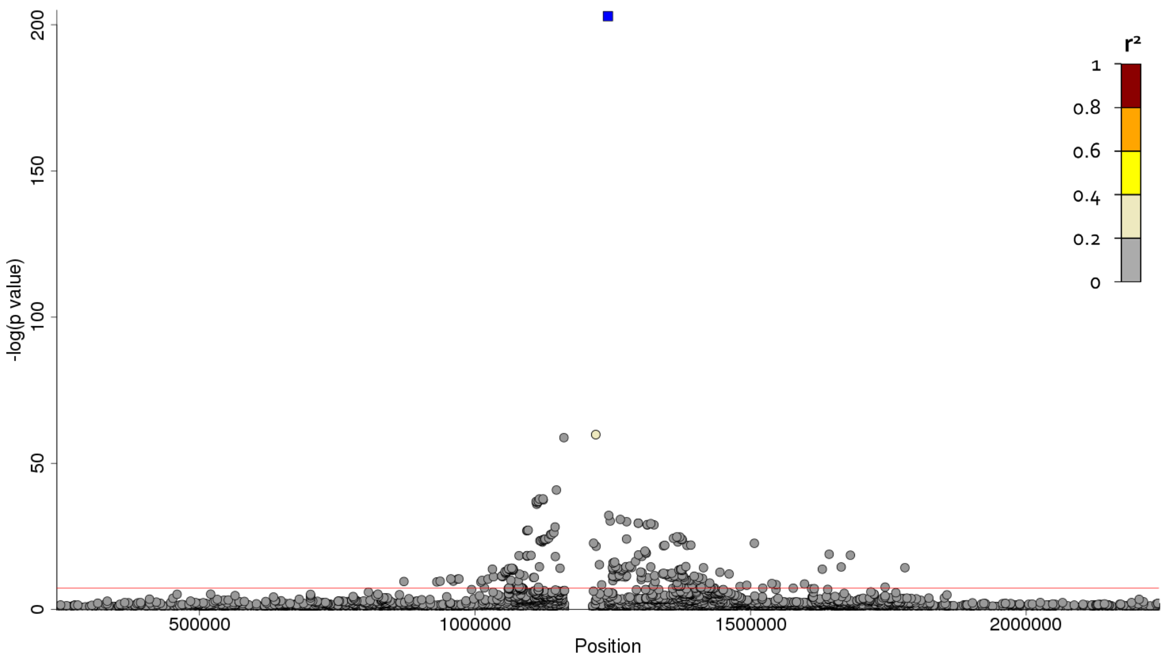

**ix) *ATP11A***

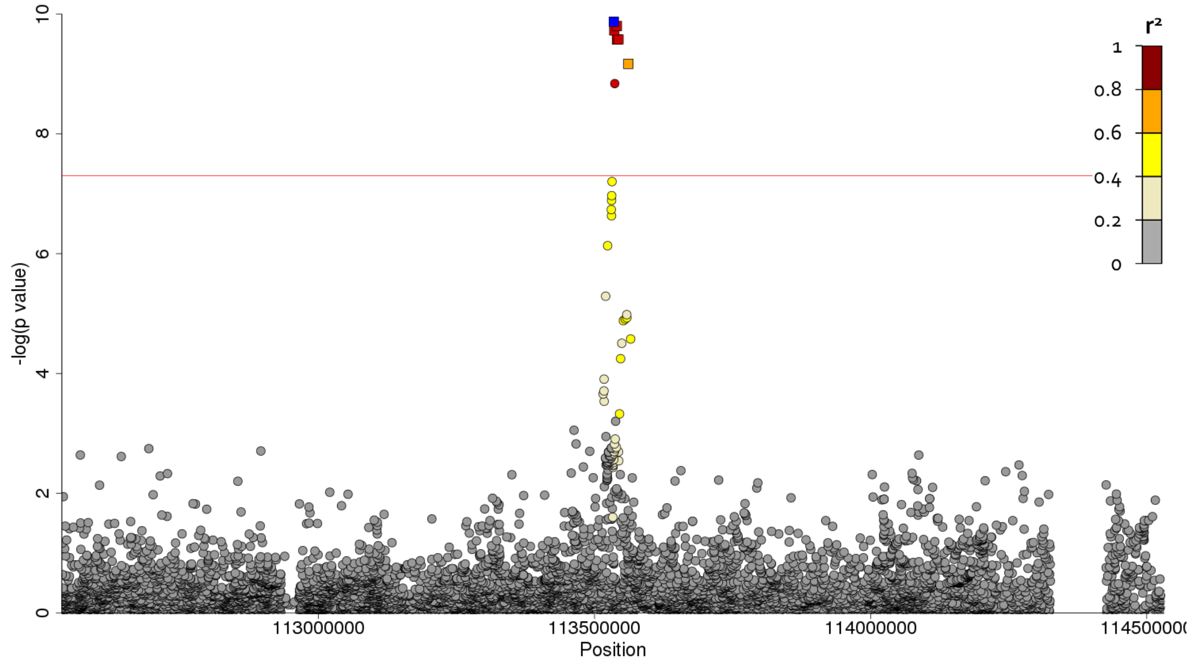

**x) *MDGA2***

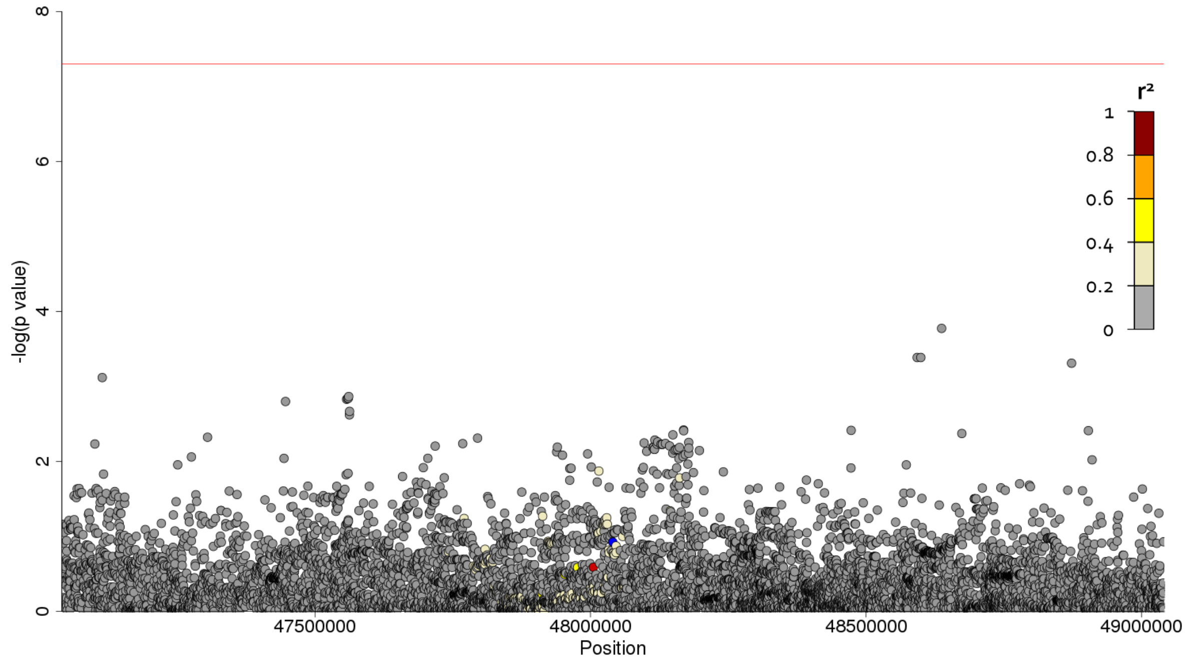

**xi) *IVD***

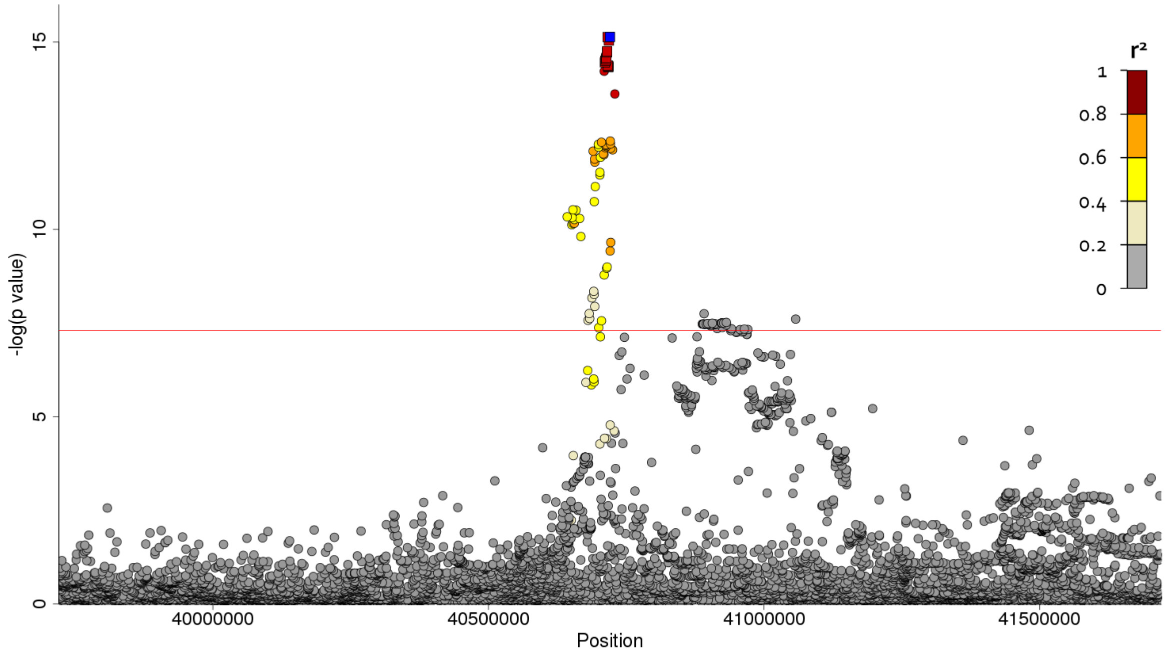

**xii) *AKAP13***

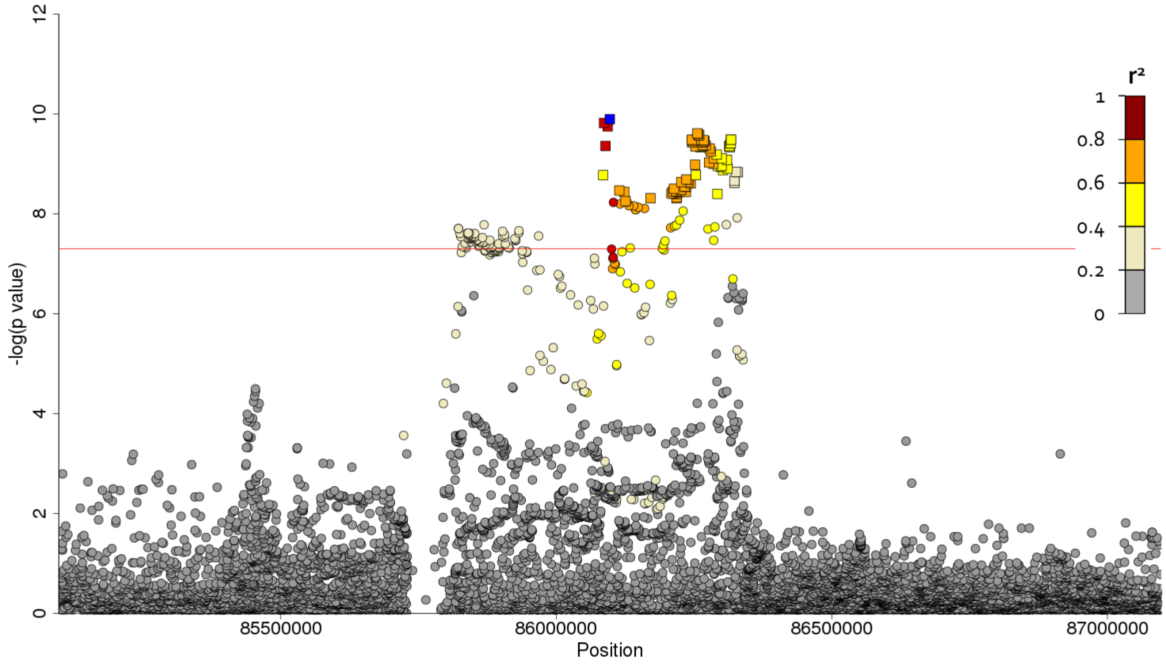

xiii) *MAPT*

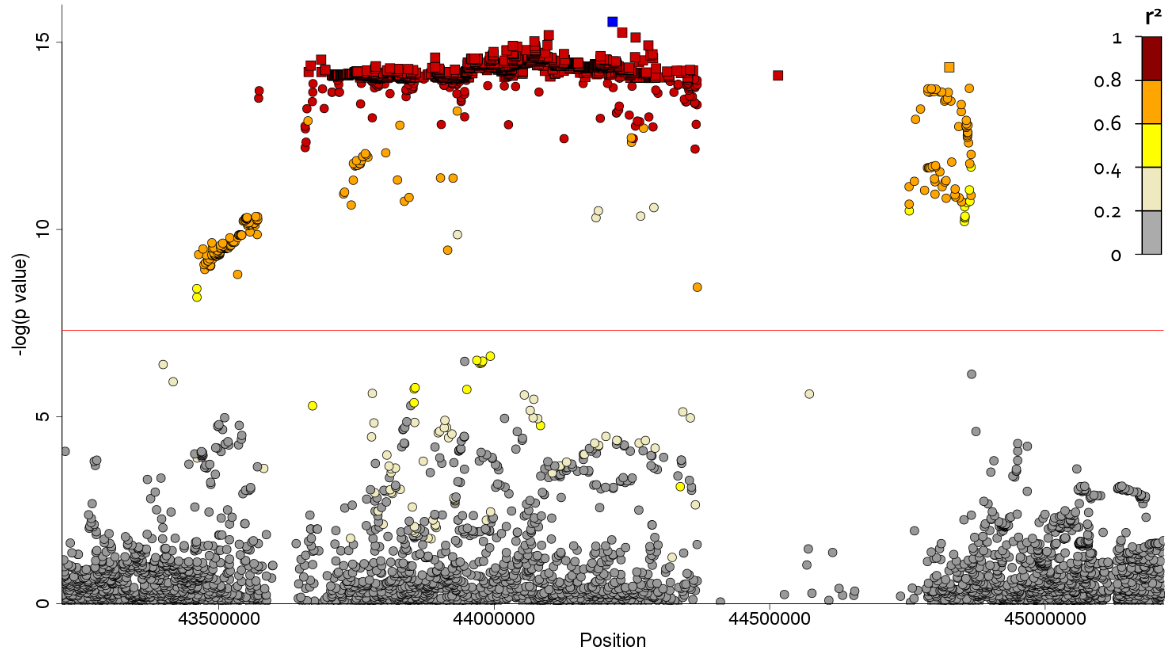

xiv) *DPP9*

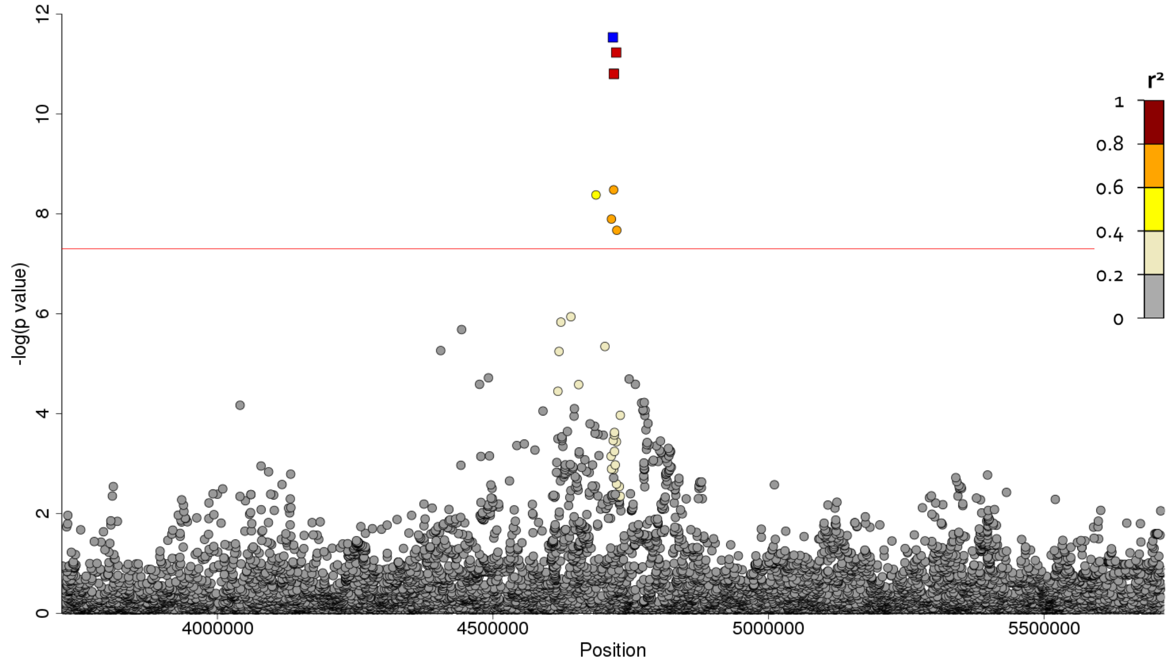

**Supplementary Figure 5 - Region plots and conditional analyses for the five novel IPF association signals in the discovery genome-wide analysis**

Region plots for each of the five novel signals in discovery analysis and after conditioning on the sentinel variant. The x axis shows chromosomal position and y axis the −log($P$ value). The sentinel is shown in blue with all other variants coloured by LD with the sentinel variant. Credible sets were calculated and variants in the credible set are shown by squares. The red horizontal line shows $P = 5 \times 10^{-8}$.

**i) Chromosome 3**

**Region plot**

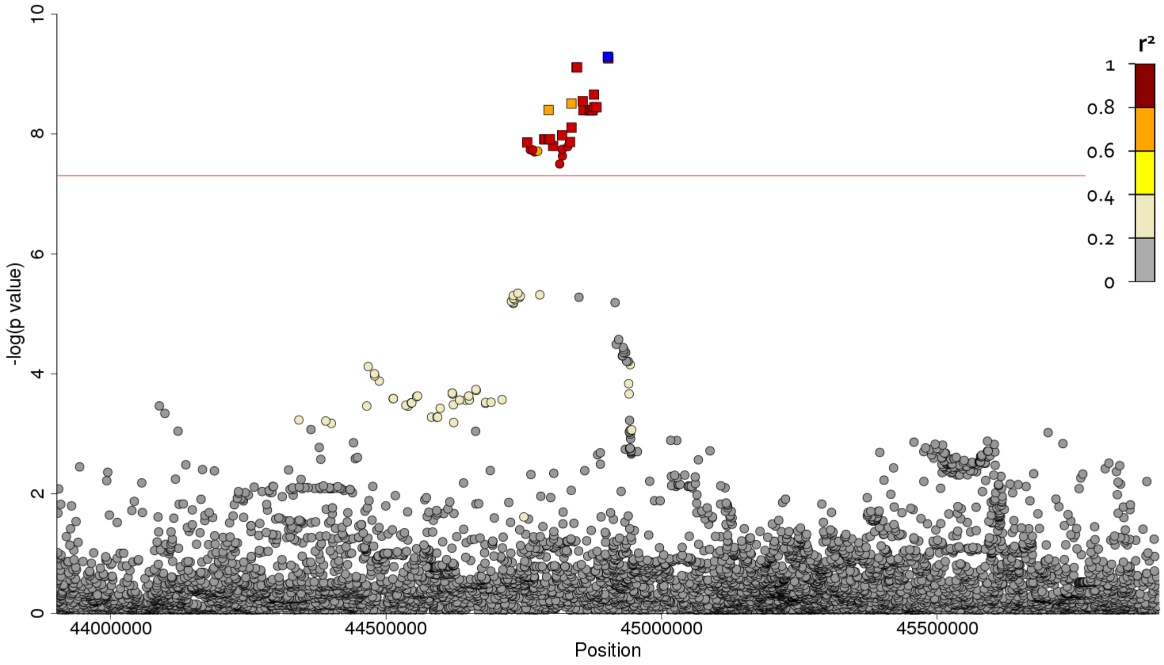

**After conditioning on sentinel variant**

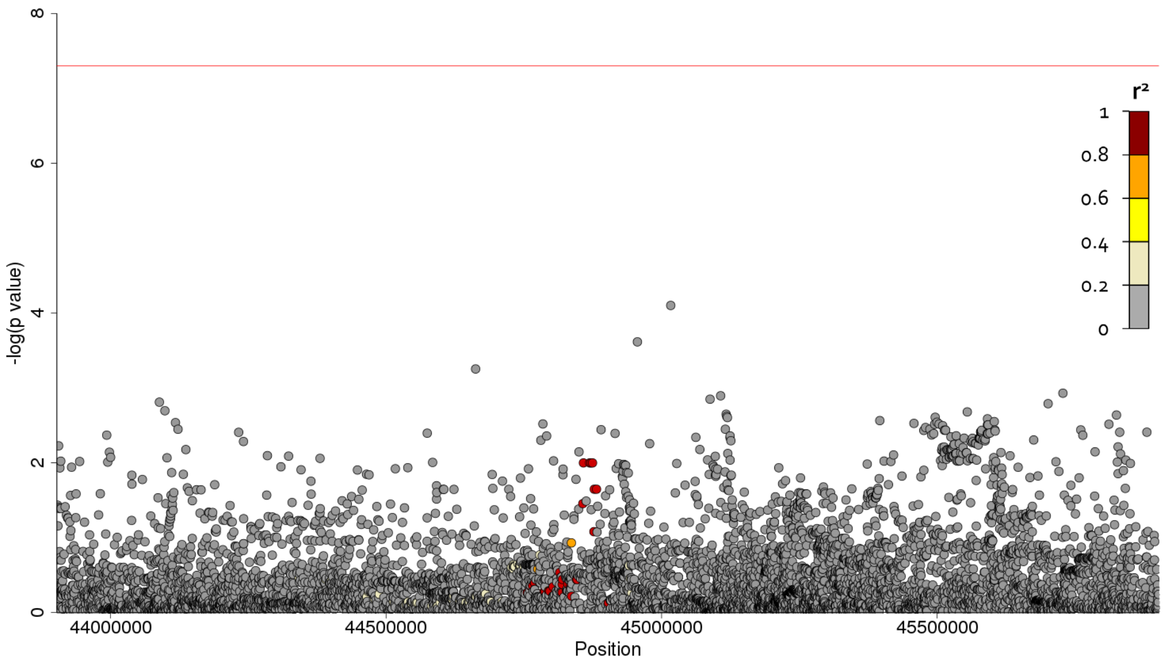

#### ii) Chromosome 7

#### Region plot

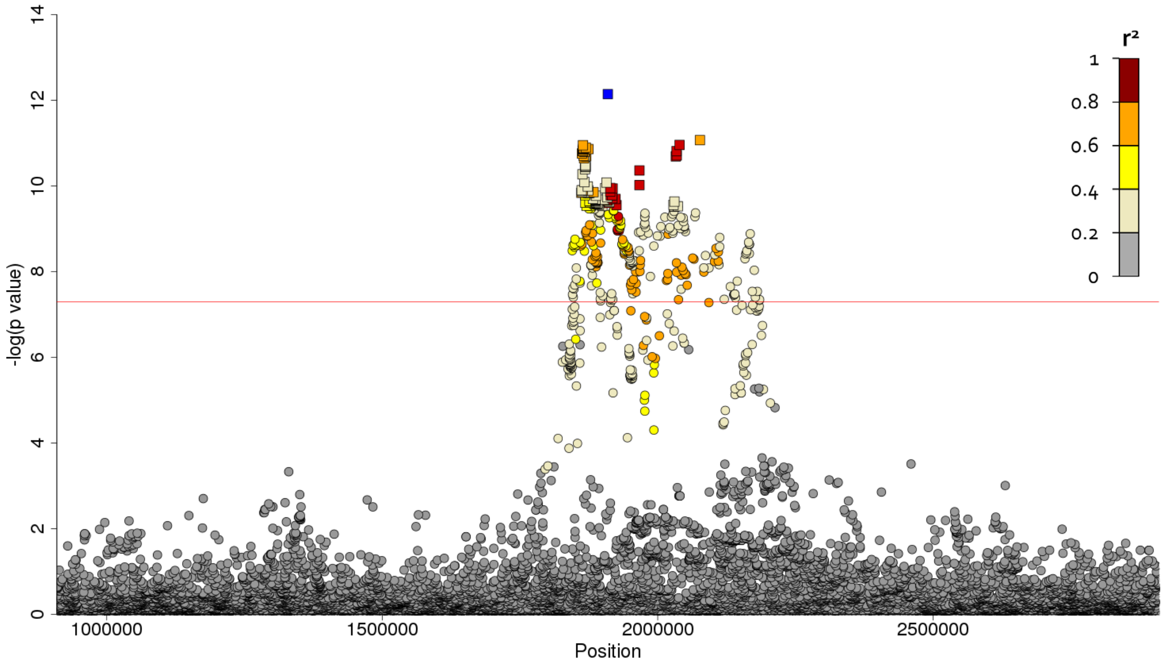

#### After conditioning on sentinel variant

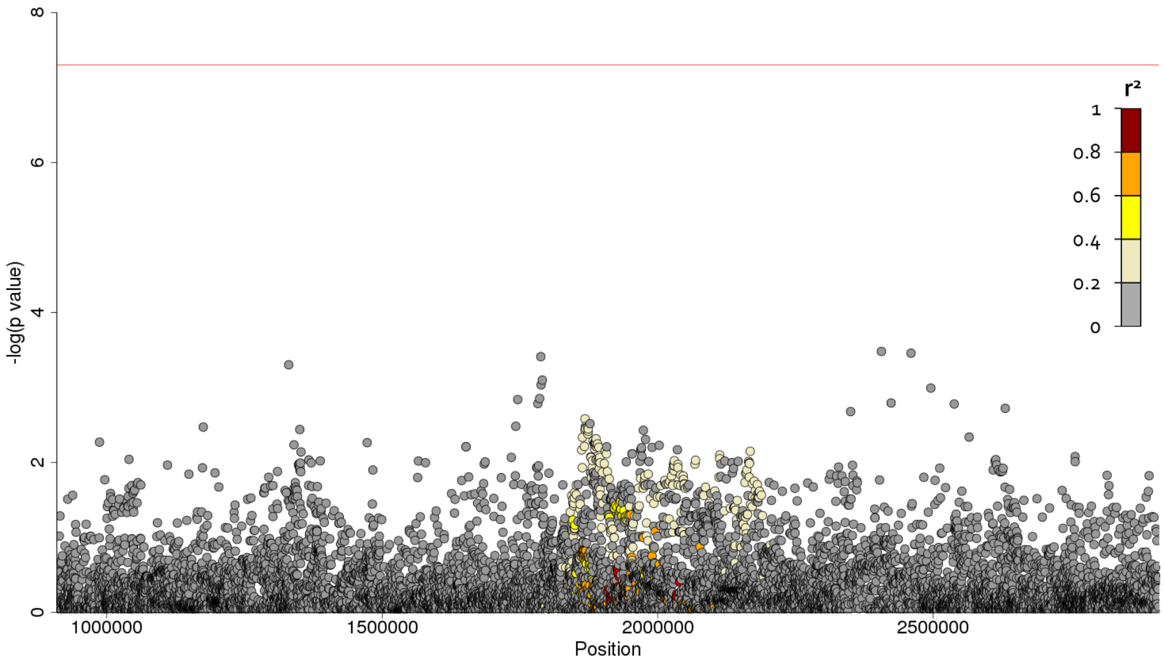

##### iii) Chromosome 8

##### Region plot

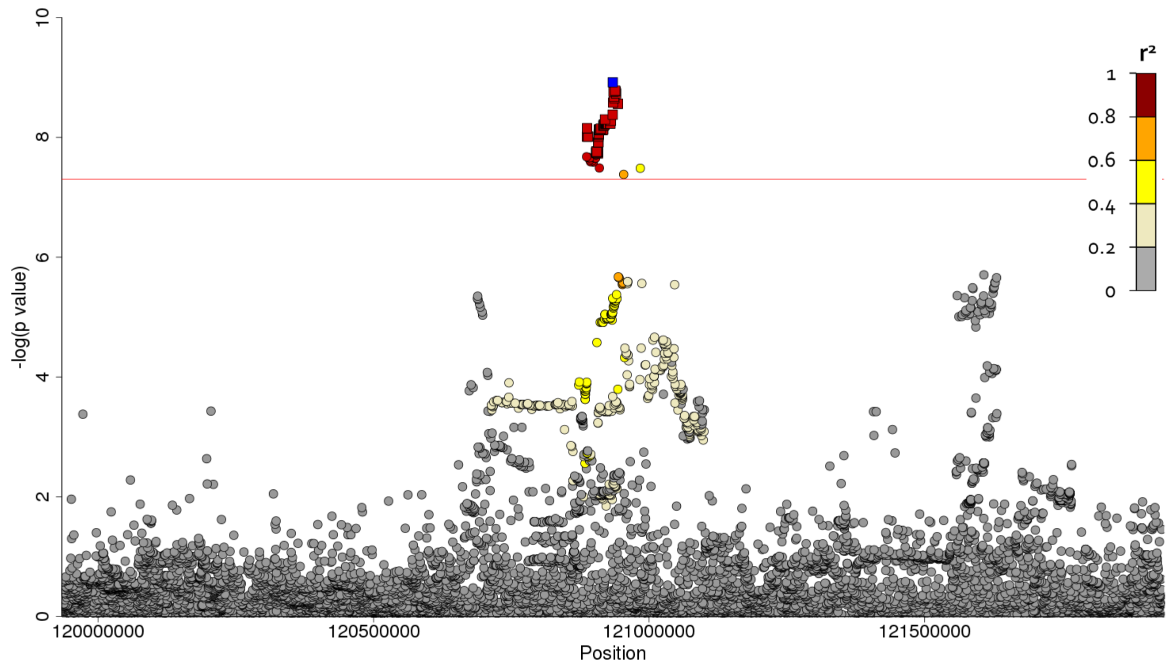

##### After conditioning on sentinel variant

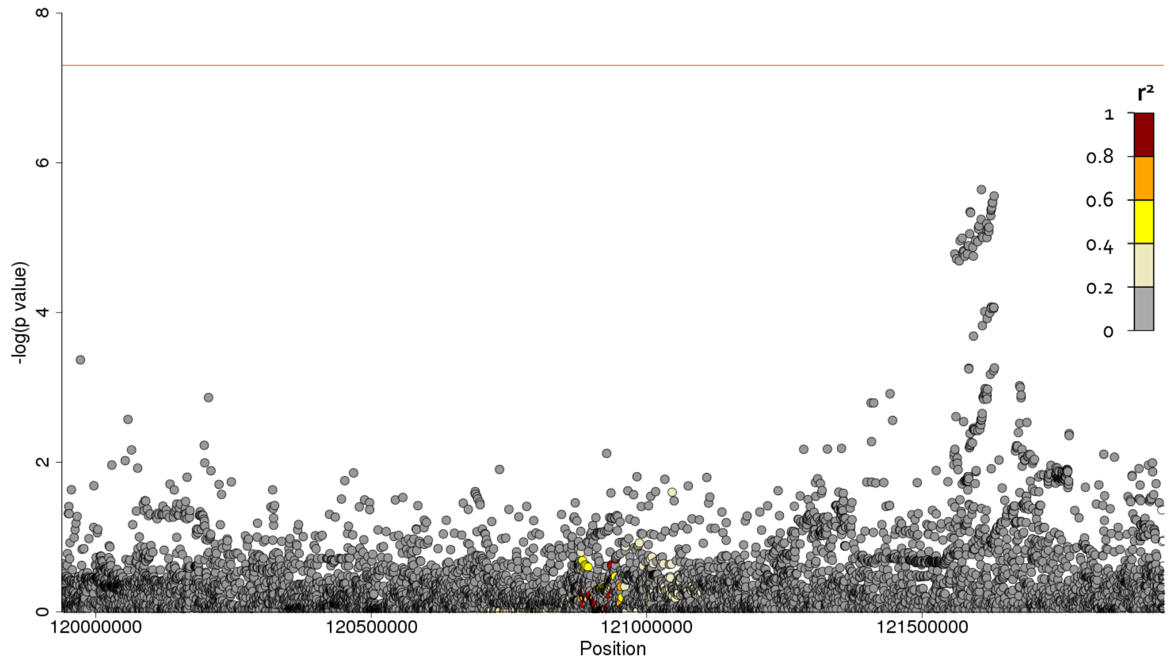

##### iv) Chromosome 10

##### Region plot

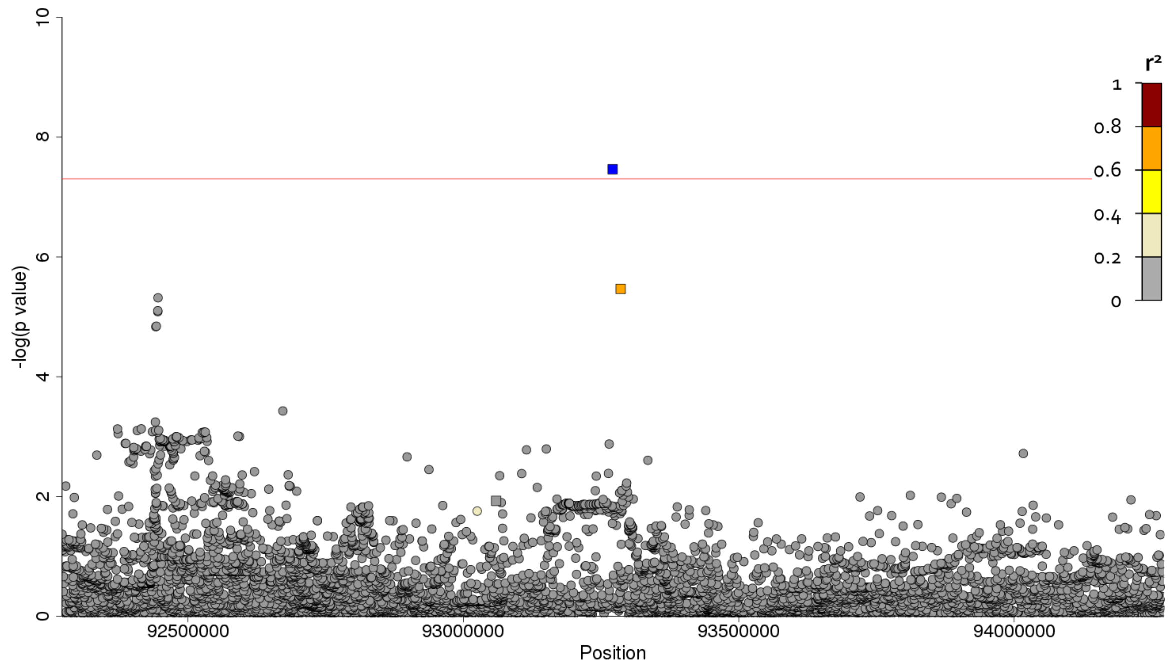

##### After conditioning on sentinel variant

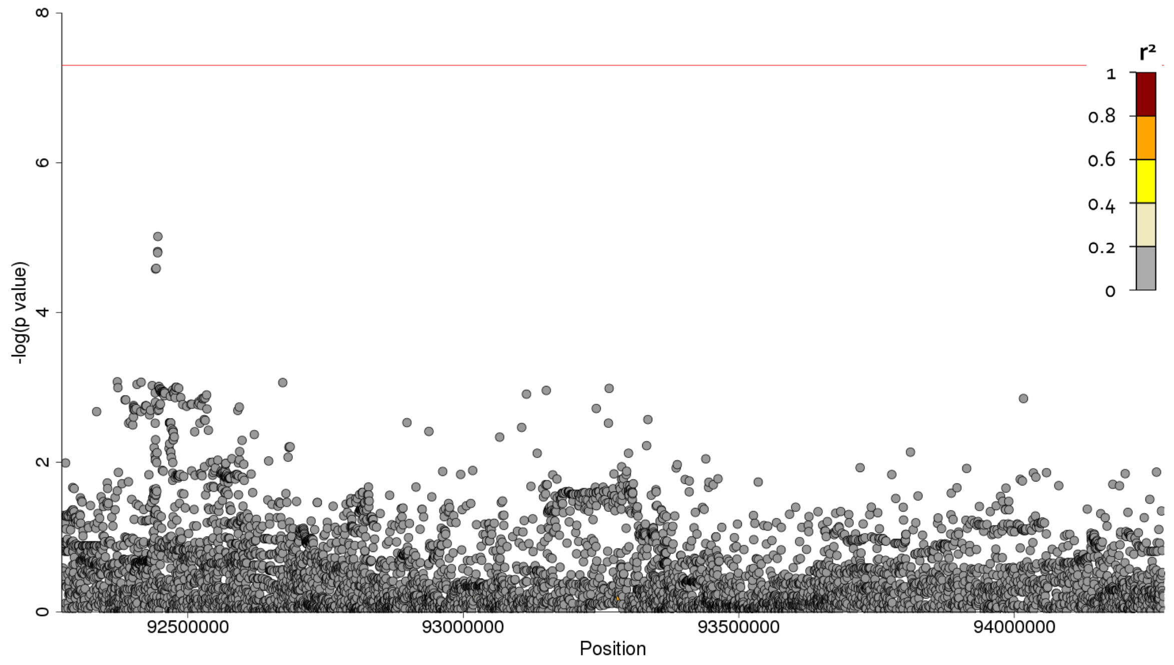

#### v) Chromosome 20

##### Region plot

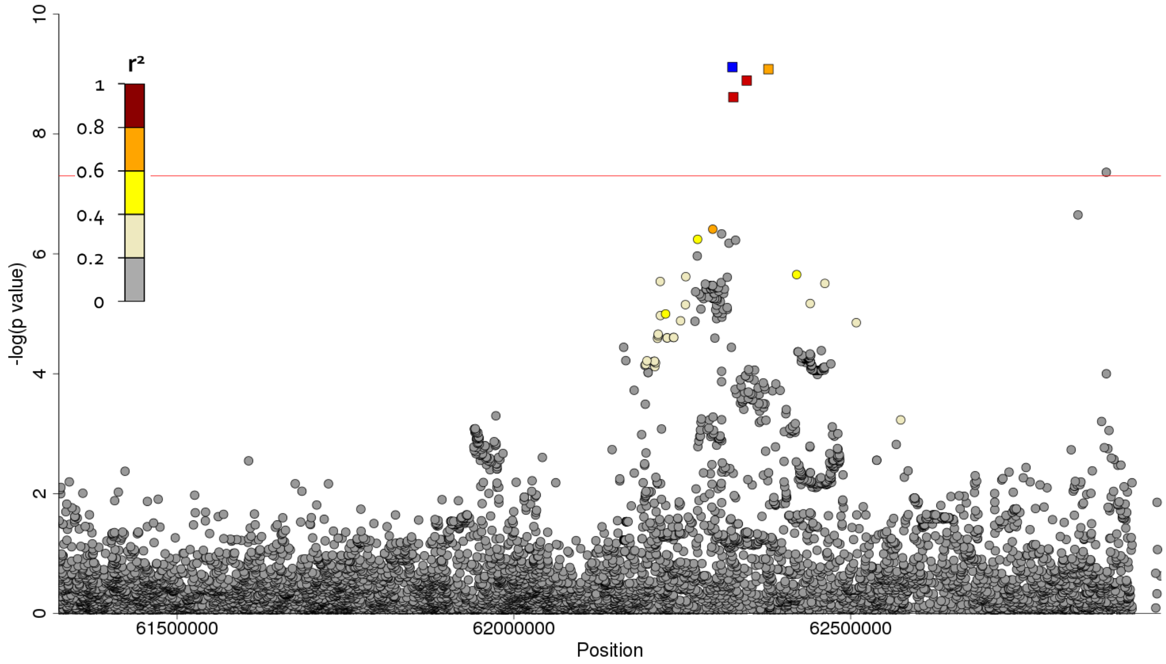

##### After conditioning on sentinel variant

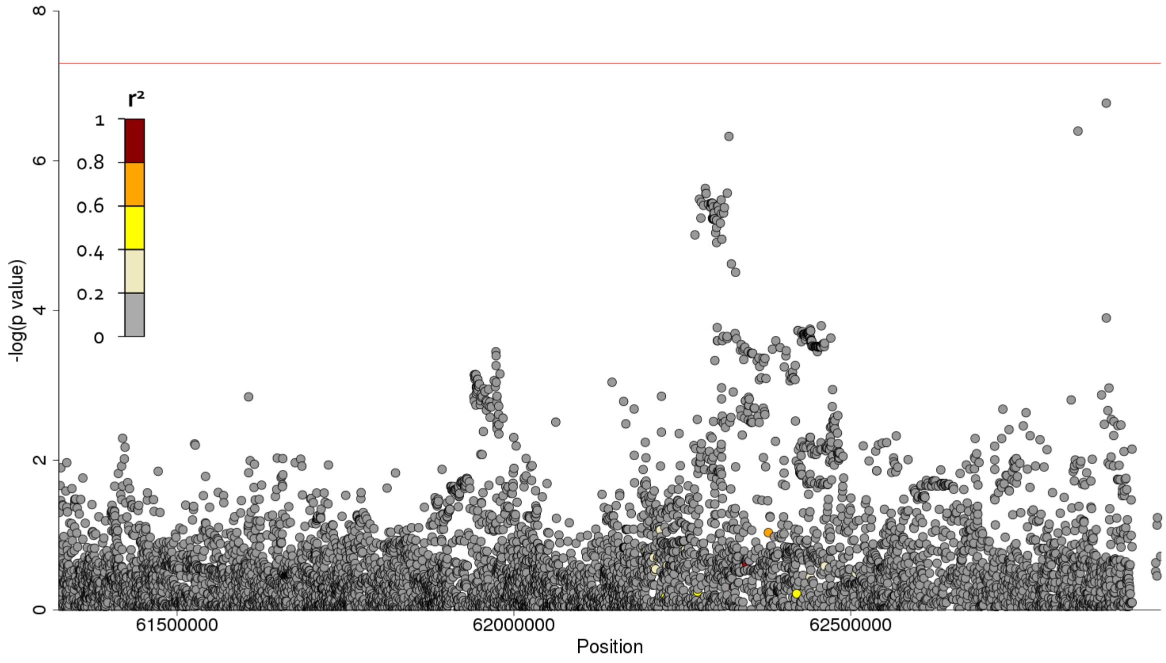

#### Supplementary Figure 6 - Forest plot of study level results

##### i) Forest plot for rs78238620

**rs78238620 (KIF15)**

| Study | OR | 95% CI | p |
|---|---|---|---|
| Chicago | 1.74 | [1.16, 2.60] | 0.007 |
| Colorado | 1.47 | [1.23, 1.78] | $4.01 \times 10^{-5}$ |
| UK | 1.77 | [1.35, 2.33] | $4.54 \times 10^{-5}$ |
| **Discovery meta-analysis** | **1.58** | **[1.37, 1.83]** | $\mathbf{5.12 \times 10^{-10}}$ |
| UUS | 1.46 | [1.18, 1.80] | $4.39 \times 10^{-4}$ |
| Genentech | 1.55 | [1.11, 2.16] | 0.010 |
| **Replication meta-analysis** | **1.48** | **[1.24, 1.77]** | $\mathbf{1.43 \times 10^{-5}}$ |
| **Discovery and replication meta** | **1.54** | **[1.38, 1.72]** | $\mathbf{4.05 \times 10^{-14}}$ |

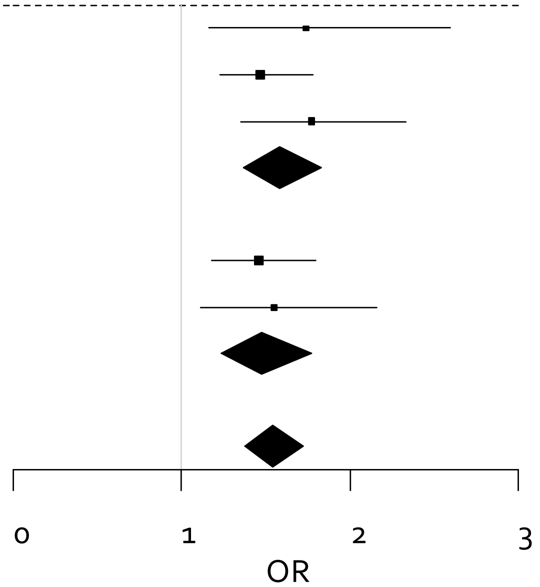

##### ii) Forest plot for rs12699415

**rs12699415 (MAD1L1)**

| Study | OR | 95% CI | p |
|---|---|---|---|
| Chicago | 1.43 | [1.20, 1.69] | $5.31 \times 10^{-5}$ |
| Colorado | 1.23 | [1.12, 1.33] | $3.67 \times 10^{-6}$ |
| UK | 1.30 | [1.15, 1.47] | $3.51 \times 10^{-5}$ |
| **Discovery meta-analysis** | **1.28** | **[1.19, 1.37]** | $\mathbf{7.15 \times 10^{-13}}$ |
| UUS | 1.24 | [1.12, 1.38] | $6.39 \times 10^{-5}$ |
| Genentech | 1.41 | [1.20, 1.66] | $3.82 \times 10^{-5}$ |
| **Replication meta-analysis** | **1.29** | **[1.18, 1.41]** | $\mathbf{2.27 \times 10^{-8}}$ |
| **Discovery and replication meta** | **1.28** | **[1.21, 1.35]** | $\mathbf{9.38 \times 10^{-20}}$ |

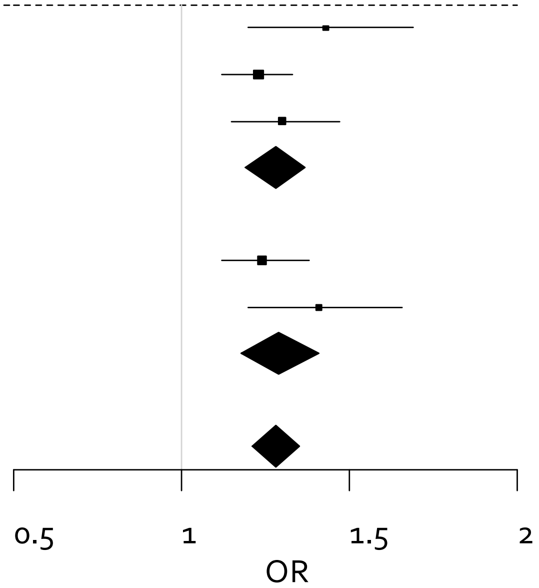

##### iii) Forest plot for rs28513081

**rs28513081 (DEPTOR)**

| Study | OR | 95% CI | p |
|---|---|---|---|
| Chicago | 0.78 | [0.66, 0.93] | 0.005 |
| Colorado | 0.84 | [0.77, 0.91] | $4.69 \times 10^{-5}$ |
| UK | 0.79 | [0.70, 0.89] | $1.94 \times 10^{-4}$ |
| **Discovery meta-analysis** | **0.82** | **[0.76, 0.87]** | **$1.20 \times 10^{-9}$** |
| UUS | 0.90 | [0.81, 1.00] | 0.050 |
| Genentech | 0.81 | [0.69, 0.95] | 0.010 |
| **Replication meta-analysis** | **0.87** | **[0.80, 0.95]** | **0.002** |
| **Discovery and replication meta** | **0.83** | **[0.79, 0.88]** | **$1.84 \times 10^{-11}$** |

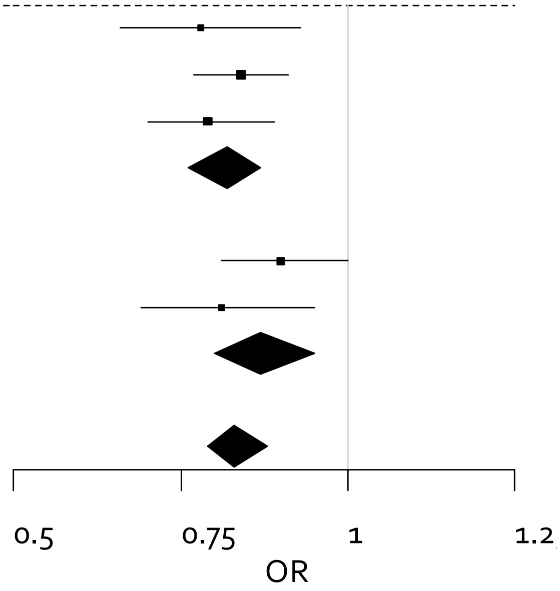

##### iv) Forest plot for rs537322302

**rs537322302 (HECTD2)**

| Study | OR | 95% CI | p |
|---|---|---|---|
| Chicago | - | - | - |
| Colorado | 7.52 | [2.46, 23.0] | $3.98 \times 10^{-4}$ |
| UK | 8.04 | [3.10, 20.8] | $1.79 \times 10^{-5}$ |
| **Discovery meta-analysis** | **7.82** | **[3.77, 16.2]** | **$3.43 \times 10^{-8}$** |
| UUS | 1.75 | [0.81, 3.78] | 0.155 |
| Genentech | - | - | - |
| **Replication meta-analysis** | **1.75** | **[0.81, 3.78]** | **0.155** |
| **Discovery and replication meta** | **3.85** | **[2.27, 6.54]** | **$6.25 \times 10^{-7}$** |

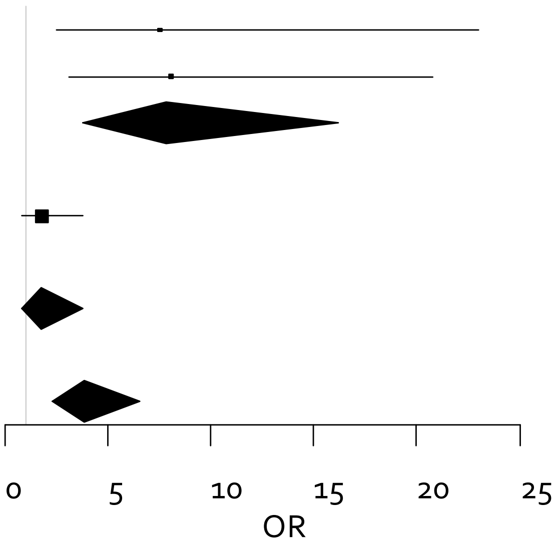

**v) Forest plot for rs41308092**

**rs41308092 (RTEL1)**

| Study | OR | 95% CI | p |
|---|---|---|---|
| Chicago | 2.06 | [1.03, 4.12] | 0.040 |
| Colorado | 2.10 | [1.54, 2.86] | $2.61 \times 10^{-6}$ |
| UK | 2.18 | [1.41, 3.37] | $4.86 \times 10^{-4}$ |
| **Discovery meta-analysis** | **2.12** | **[1.67, 2.69]** | **$7.65 \times 10^{-10}$** |
| UUS | 1.61 | [1.14, 2.28] | 0.007 |
| Genentech | 1.13 | [0.66, 1.93] | 0.660 |
| **Replication meta-analysis** | **1.45** | **[1.08, 1.94]** | **0.012** |
| **Discovery and replication meta** | **1.82** | **[1.51, 2.19]** | **$2.24 \times 10^{-10}$** |

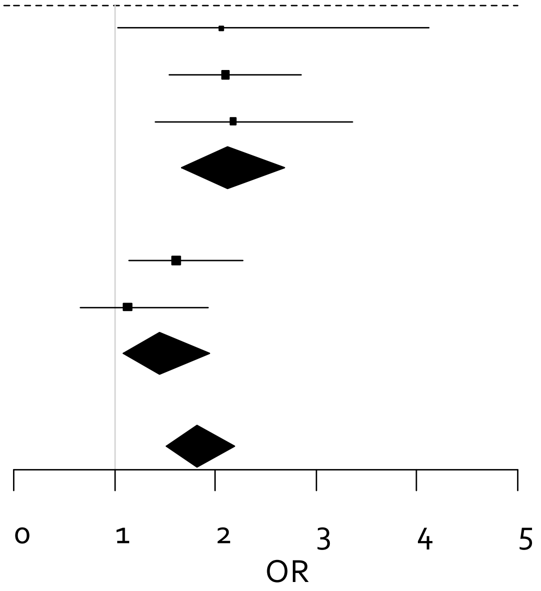

**<u>Supplementary Figure 7 - GWAS vs eQTL results for chromosome 3 signal</u>**

Each point represents a variant with chromosomal position on the x axis and −log(*P* value) on the y axis. Above the x axis is the −log(*P* value) from the IPF susceptibility discovery genome-wide meta-analysis and below the y axis is the −log(*P* value) from the eQTL database. The sentinel variant from the IPF susceptibility analysis is coloured in blue with all other variants coloured by LD with the sentinel variant (variants in red have $r^2 \geq 0.8$ with the sentinel variant, variants in orange have $0.6 \leq r^2 < 0.8$, variants in yellow have $0.4 \leq r^2 < 0.6$, variants in light yellow have $0.2 \leq r^2 < 0.4$ and variants in grey have $r^2 < 0.2$ with the sentinel variant. The area in green on the x axis denotes the location of the gene implicated by the eQTL analysis.

**i) *KIF15* - Brain (Putamen) - Colocalisation probability = 95.6%**

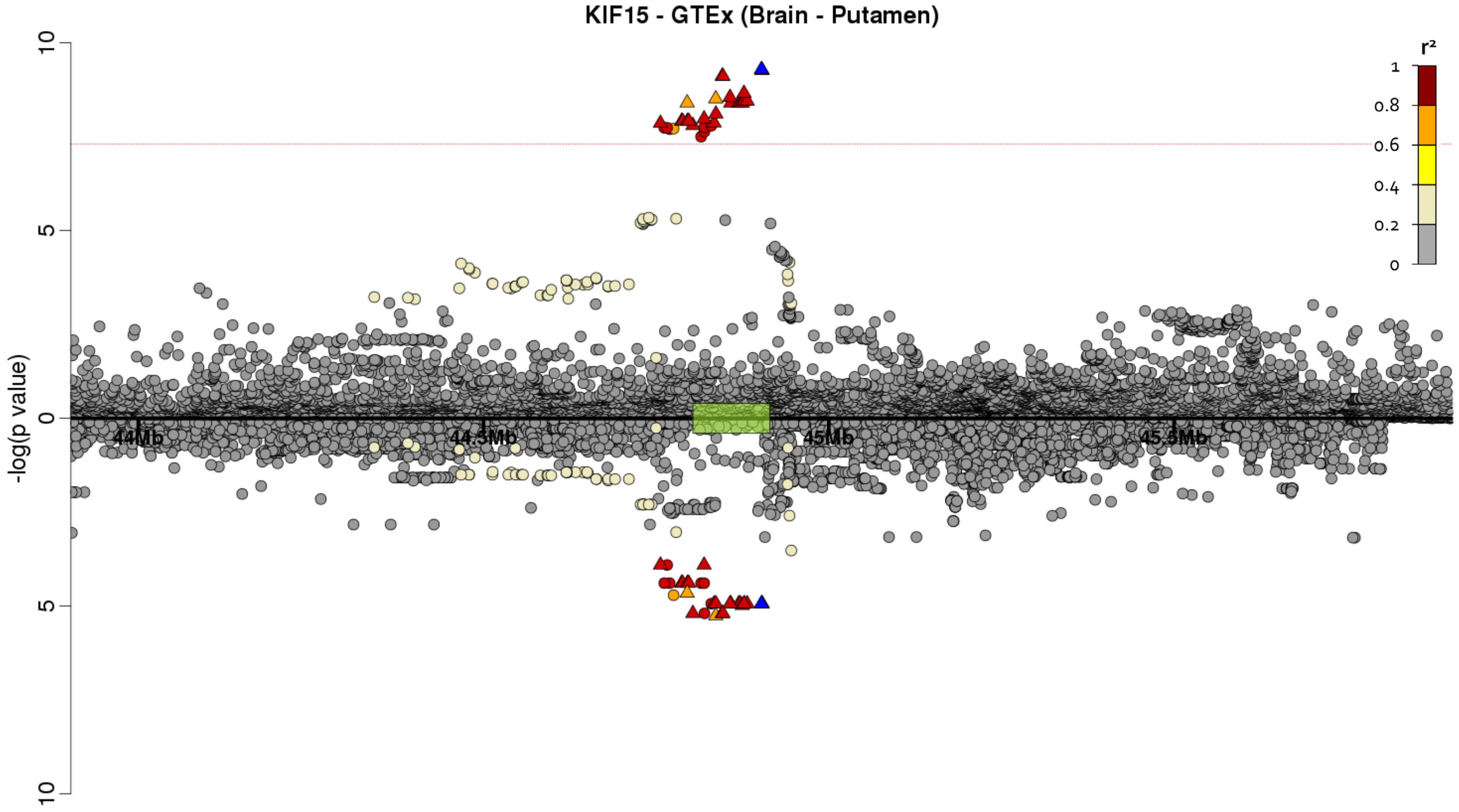

**ii) *TMEM42* - Thyroid - Colocalisation probability = 93.1%**

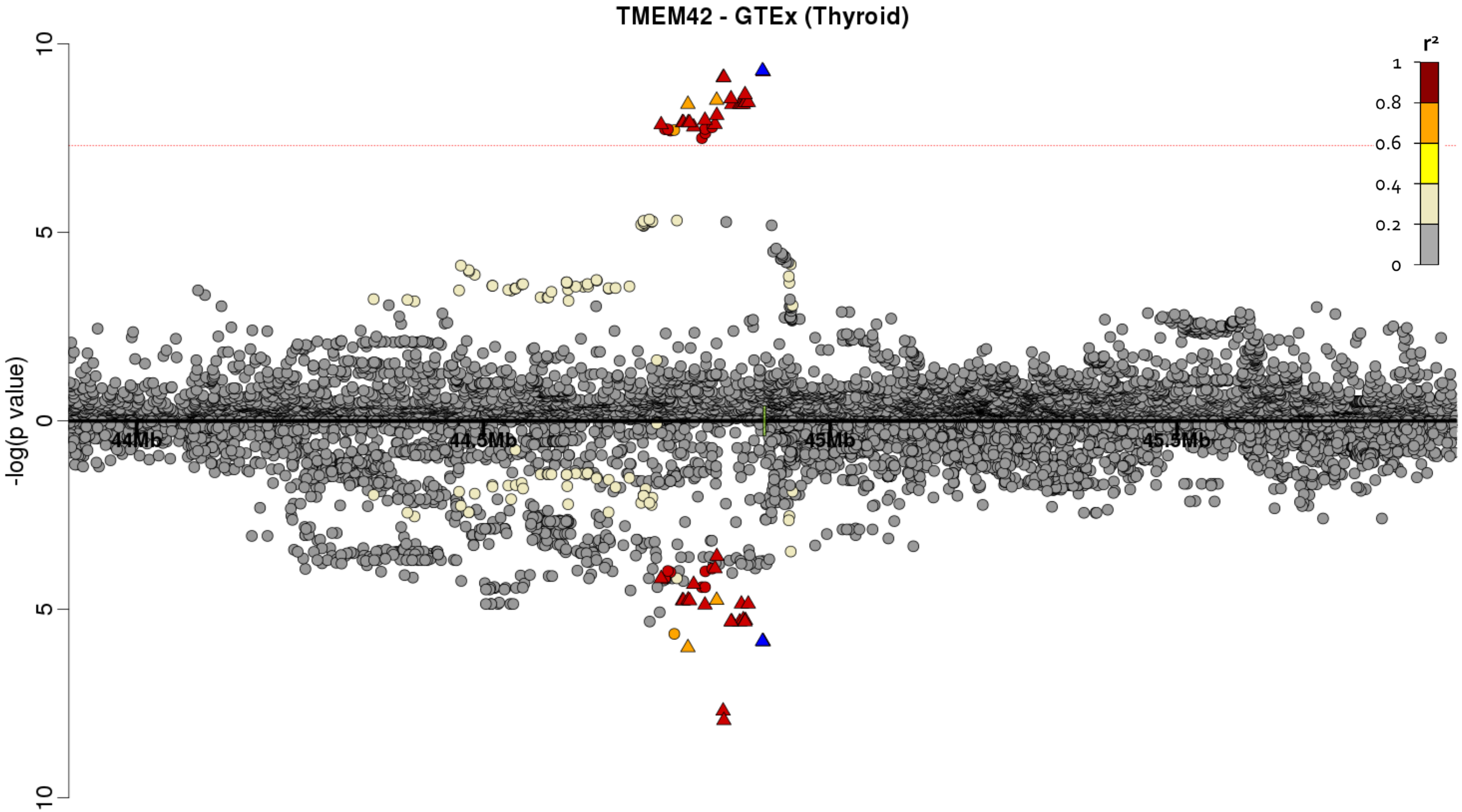

#### Supplementary Figure 8 - GWAS vs eQTL for chromosome 7 signal

Each point represents a variant with chromosomal position on the x axis and −log(*P* value) on the y axis. Above the x axis is the −log(*P* value) from the IPF susceptibility discovery genome-wide meta-analysis and below the y axis is the −log(*P* value) from the eQTL database. The sentinel variant from the IPF susceptibility analysis is coloured in blue with all other variants coloured by LD with the sentinel variant (variants in red have $r^2 \geq 0.8$ with the sentinel variant, variants in orange have $0.6 \leq r^2 < 0.8$, variants in yellow have $0.4 \leq r^2 < 0.6$, variants in light yellow have $0.2 \leq r^2 < 0.4$ and variants in grey have $r^2 < 0.2$ with the sentinel variant. The area in green on the x axis denotes the location of the gene implicated by the eQTL analysis.

**i) *MAD1L1* - Heart (Atrial Appendage) - Colocalisation probability = 95.3%**

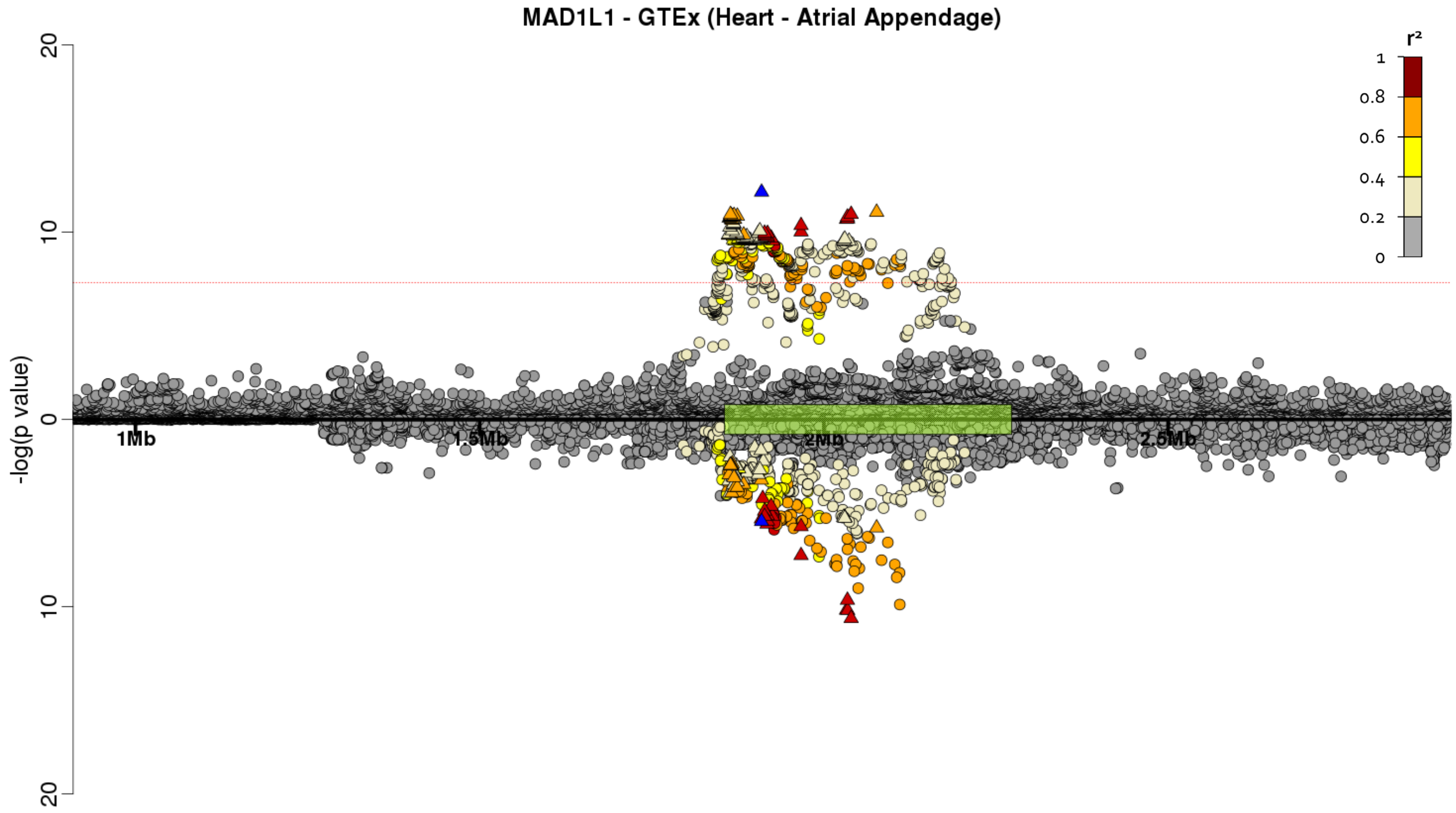

#### Supplementary Figure 9 - GWAS vs eQTL for chromosome 8 signal

Each point represents a variant with chromosomal position on the x axis and −log(*P* value) on the y axis. Above the x axis is the −log(*P* value) from the IPF susceptibility discovery genome-wide meta-analysis and below the y axis is the −log(*P* value) from the eQTL database. The sentinel variant from the IPF susceptibility analysis is coloured in blue with all other variants coloured by LD with the sentinel variant (variants in red have $r^2 \geq 0.8$ with the sentinel variant, variants in orange have $0.6 \leq r^2 < 0.8$, variants in yellow have $0.4 \leq r^2 < 0.6$, variants in light yellow have $0.2 \leq r^2 < 0.4$ and variants in grey have $r^2 < 0.2$ with the sentinel variant. The area in green on the x axis denotes the location of the gene implicated by the eQTL analysis.

**i) *DEPTOR* - Colon (Sigmoid) - Colocalisation probability = 89.6%**

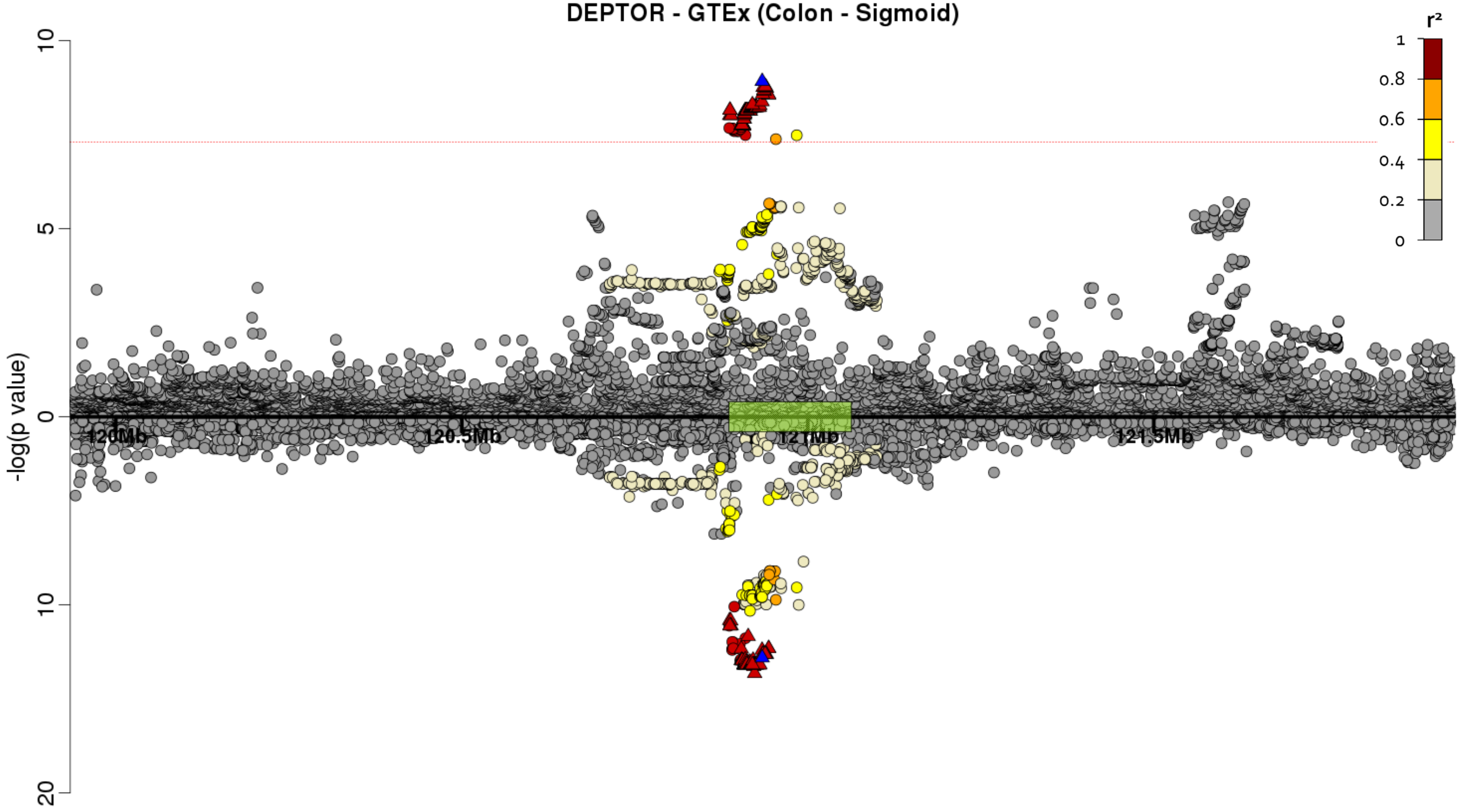

ii) *DEPTOR* - Lung - Colocalisation probability = 89.2%

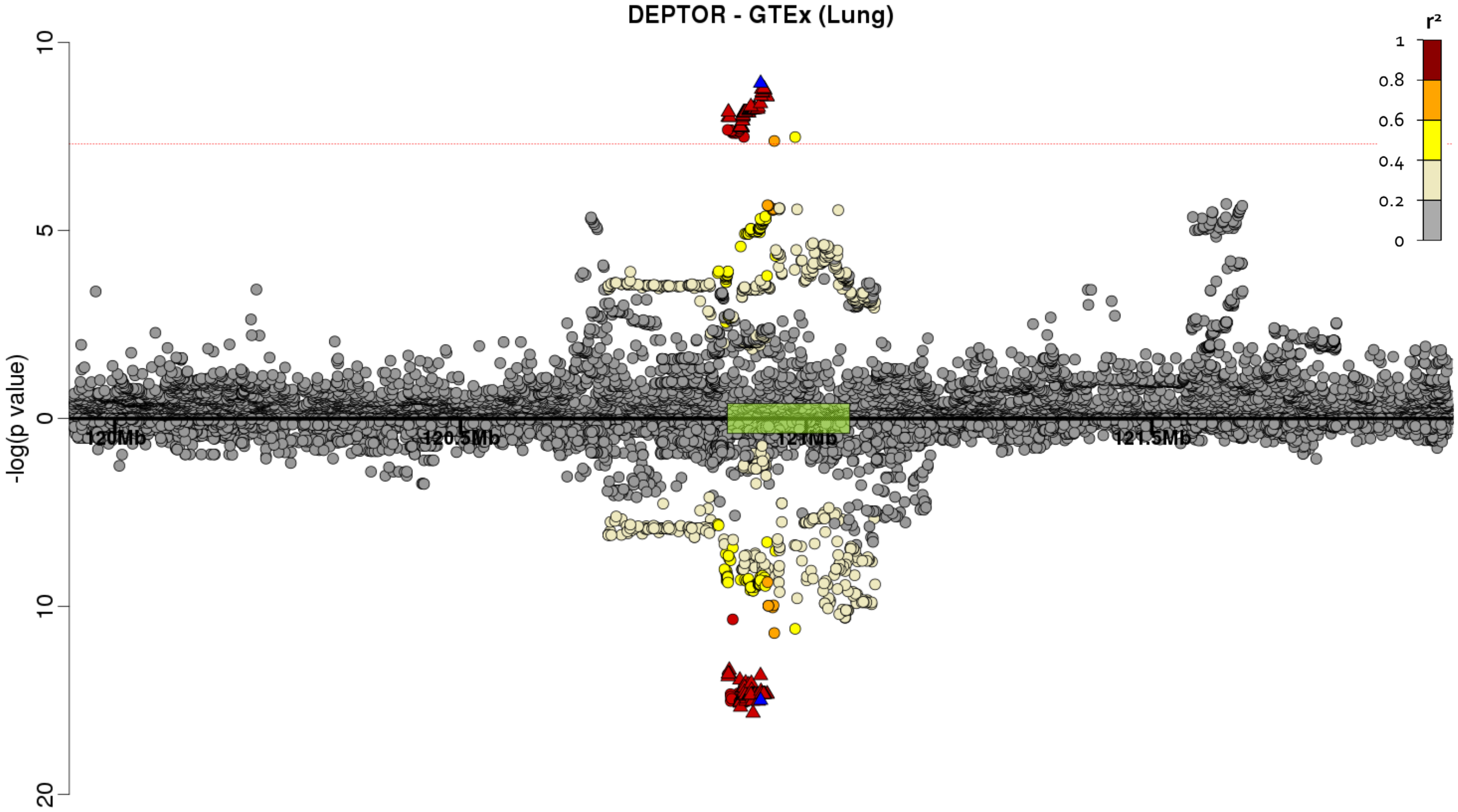

**iii) *DEPTOR* - Lung - Colocalisation probability = 89.5%**

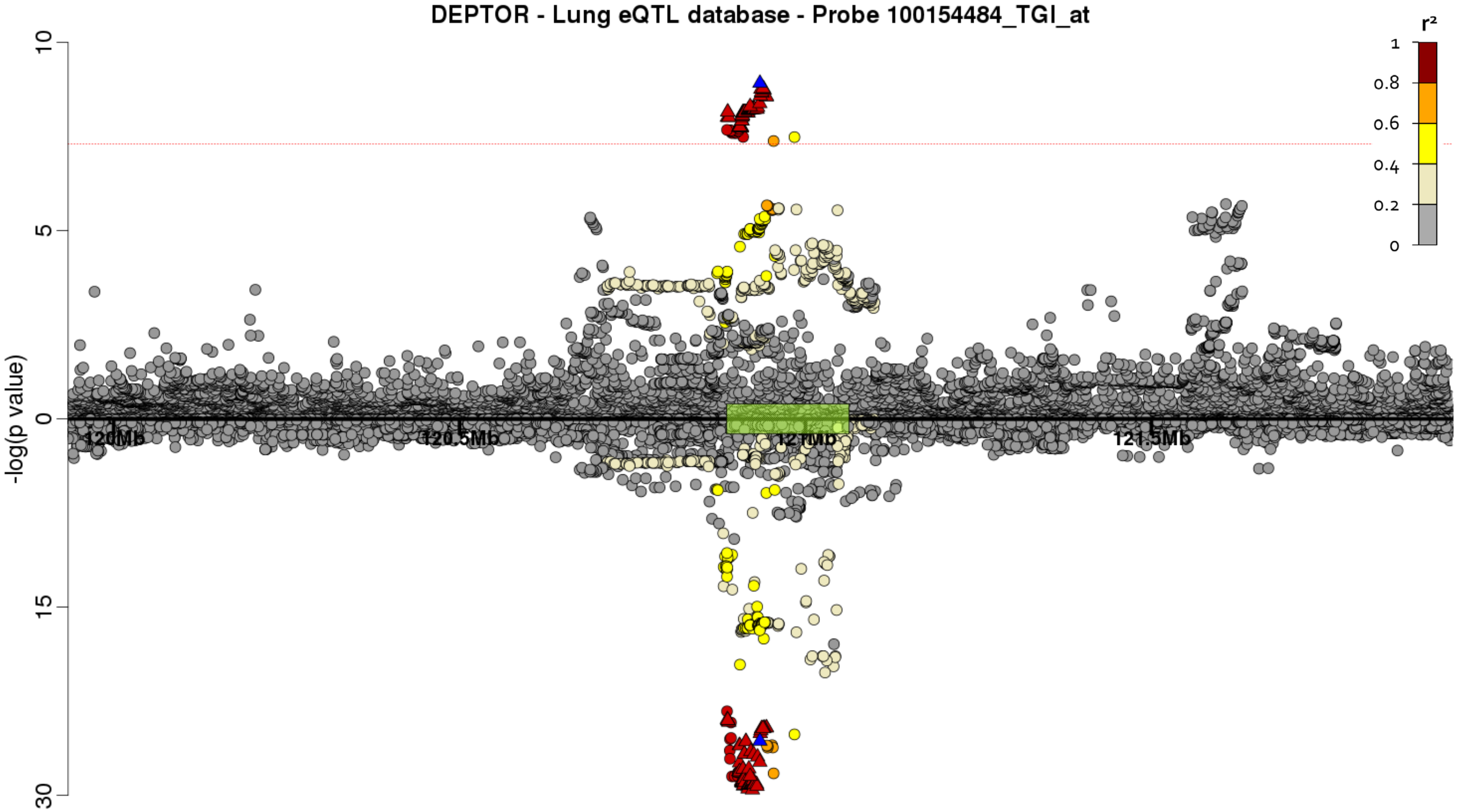

**Note:** The lung eQTL dataset showed two independent signals of association for *DEPTOR* expression. The eQTL results here are those obtained after conditioning on the top eQTL for *DEPTOR* to condition out the strongest signal which was driven by different variants to those driving the IPF risk association.

**iv) *DEPTOR* - Lung - Colocalisation probability = 89.9%**

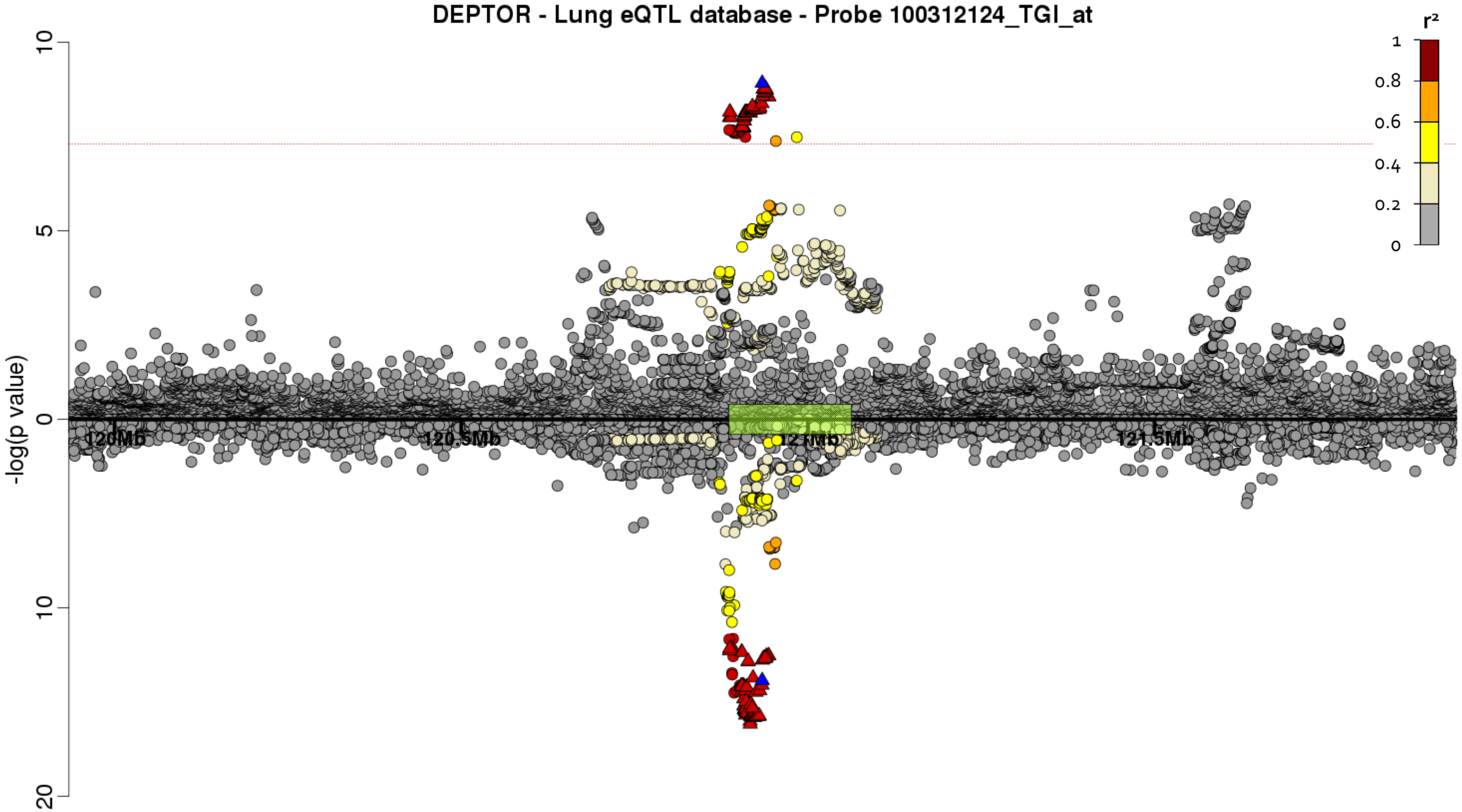

**Note:** The lung eQTL dataset showed two independent signals of association for *DEPTOR* expression. The eQTL results here are those obtained after conditioning on the top eQTL for *DEPTOR* to condition out the strongest signal which was driven by different variants to those driving the IPF risk association.

v) ***DEPTOR*** **- Skin (Not sun exposed) - Colocalisation probability = 90.0%**

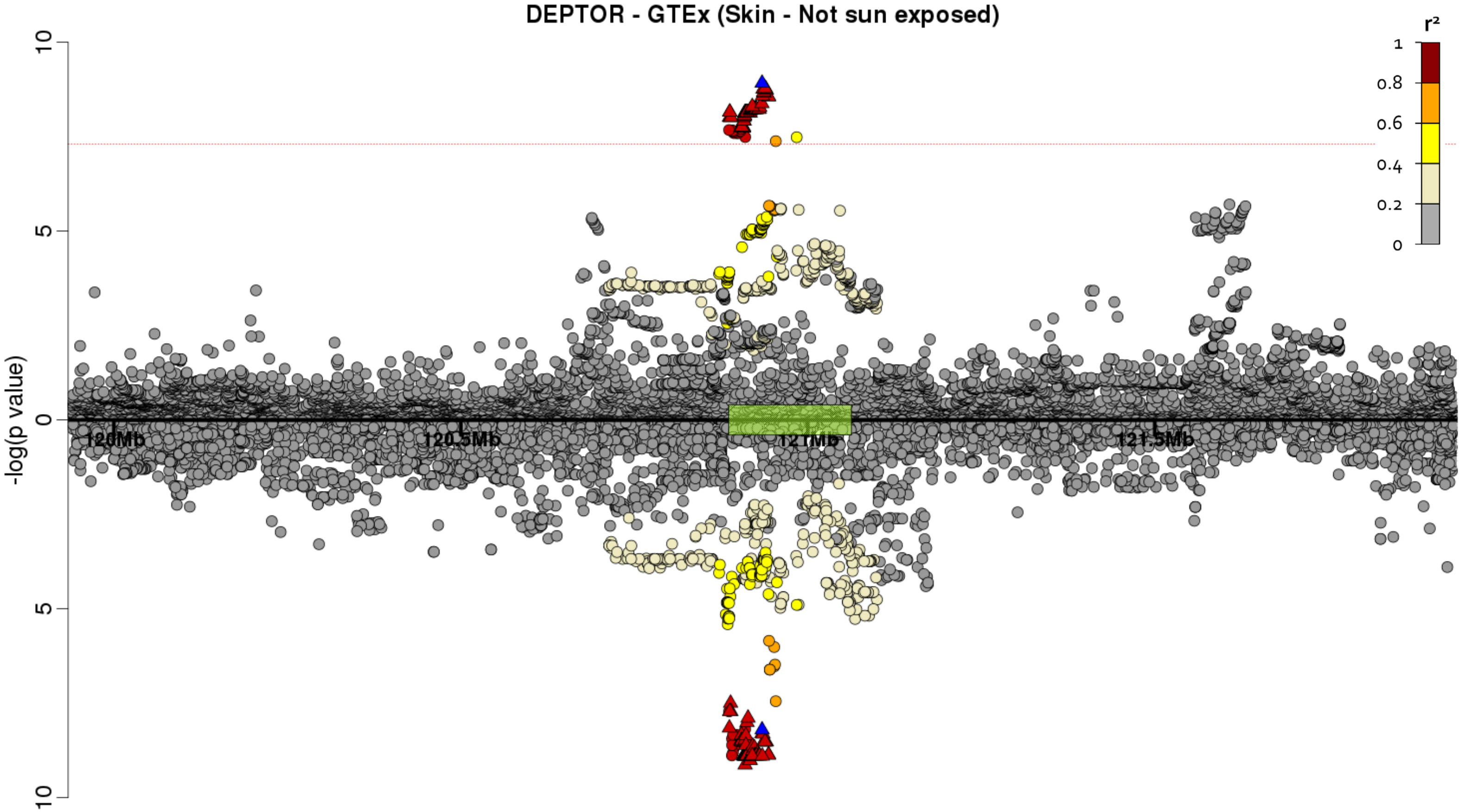

**vi) *DEPTOR* - Skin (Sun exposed) - Colocalisation probability = 86.5%**

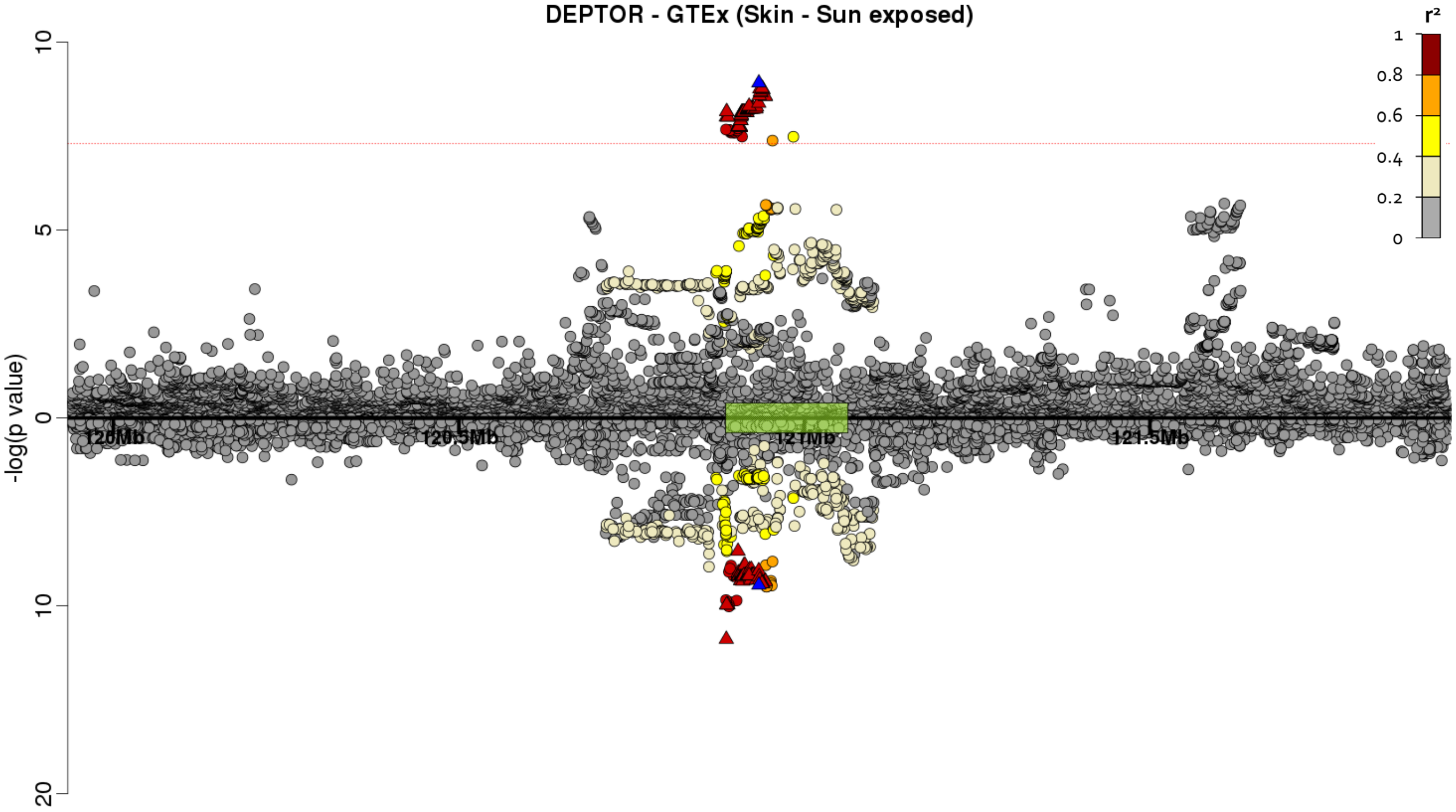

**vii) *DEPTOR* - Whole blood - Colocalisation probability = 93.7%**

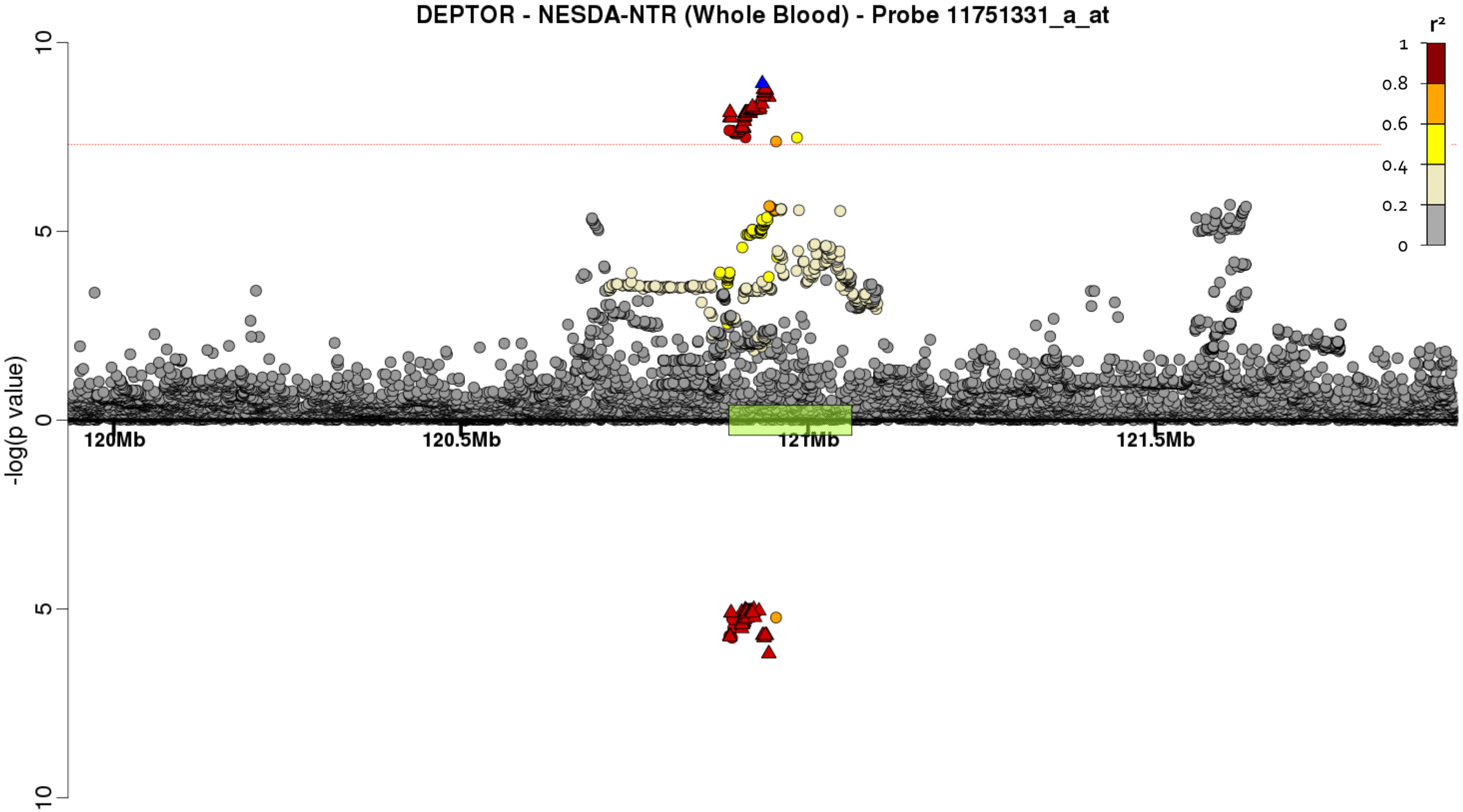

**viii) *TAF2* - Colon (Transverse) - Colocalisation probability = 87.5%**

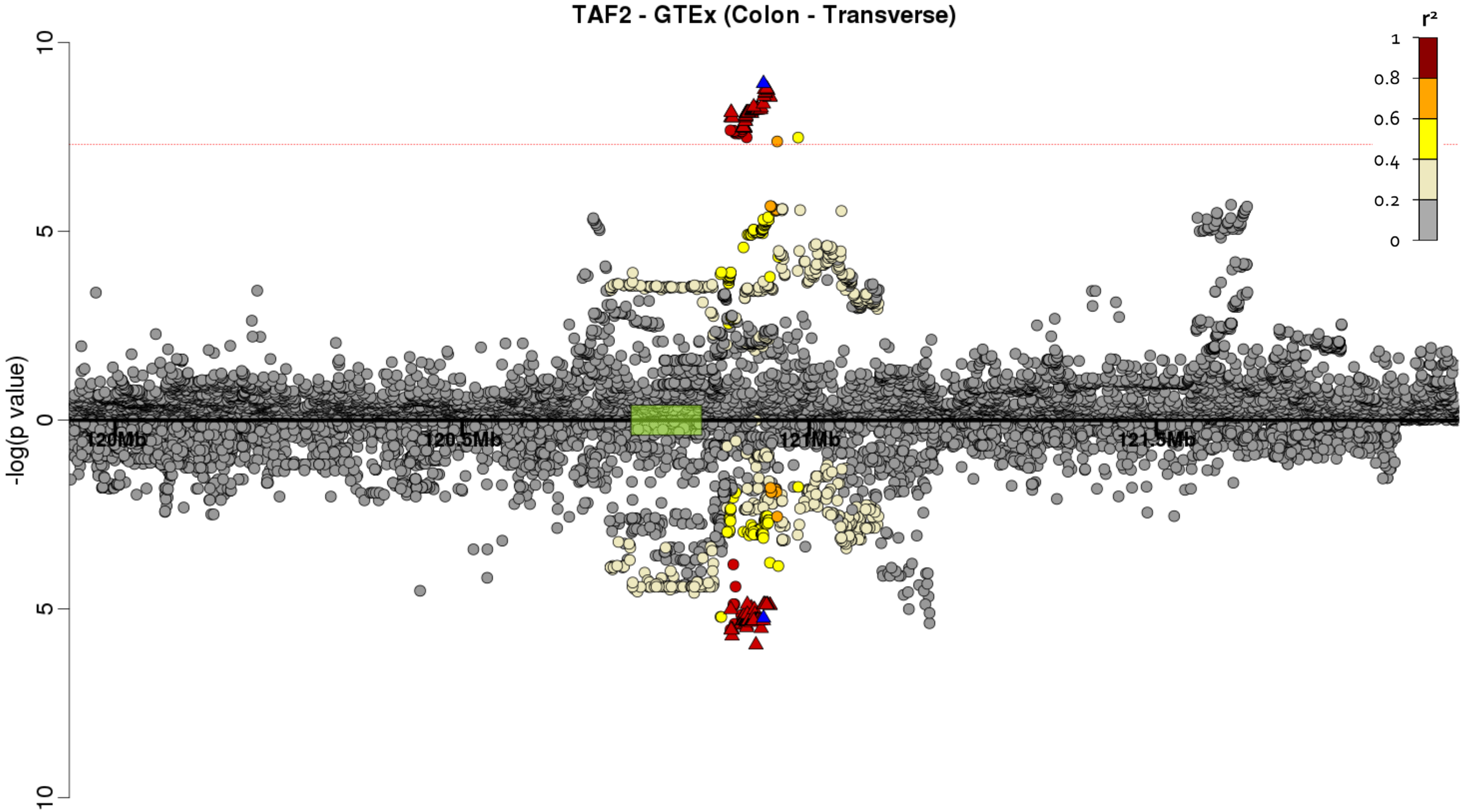

**ix) RP11-760H22.2 - Adipose (Subcutaneous) - Colocalisation probability = 84.9%**

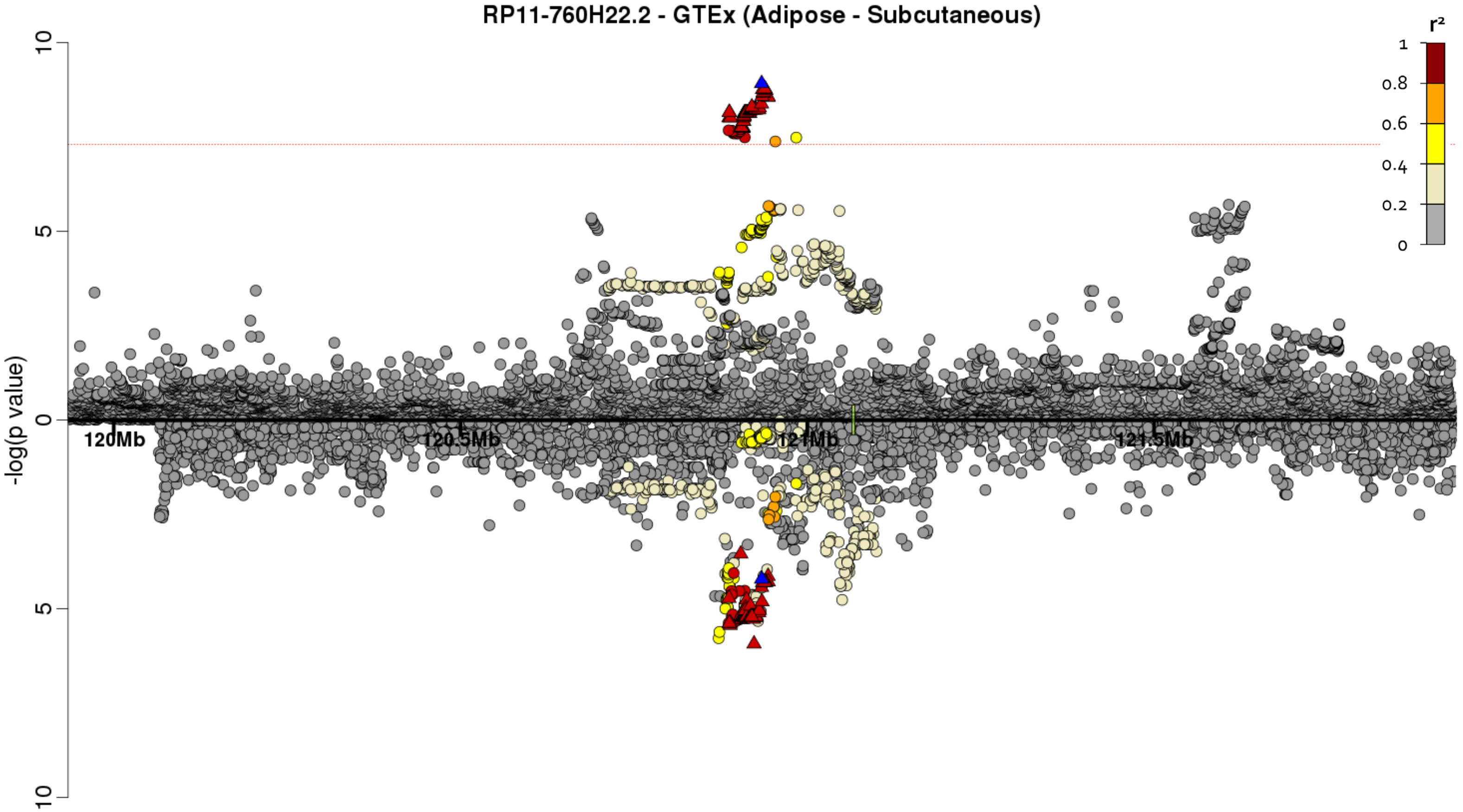

**x) RP11-760H22.2 - Colon (Sigmoid) - Colocalisation probability = 88.6%**

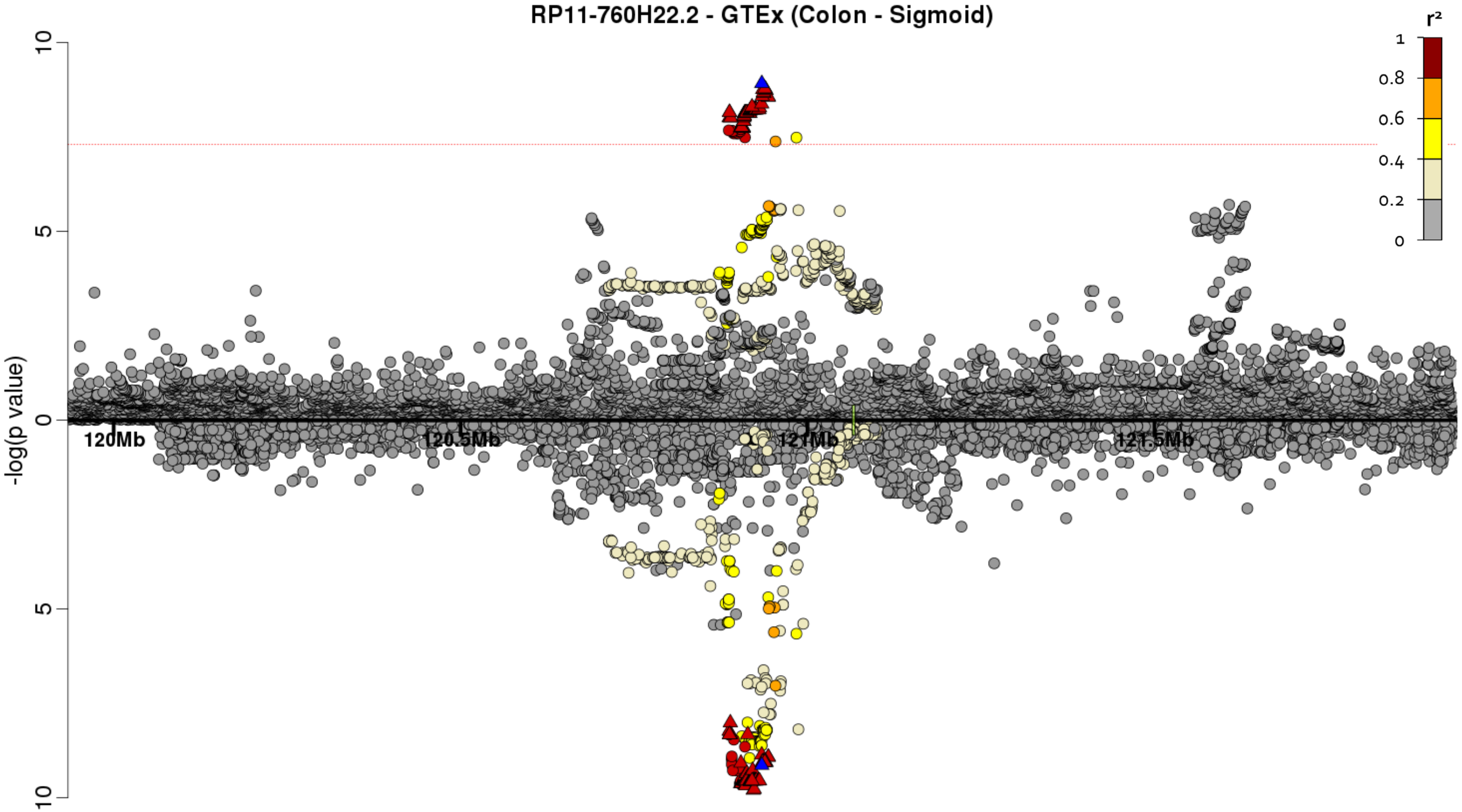

**xi) RP11-760H22.2 - Lung - Colocalisation probability = 90.0%**

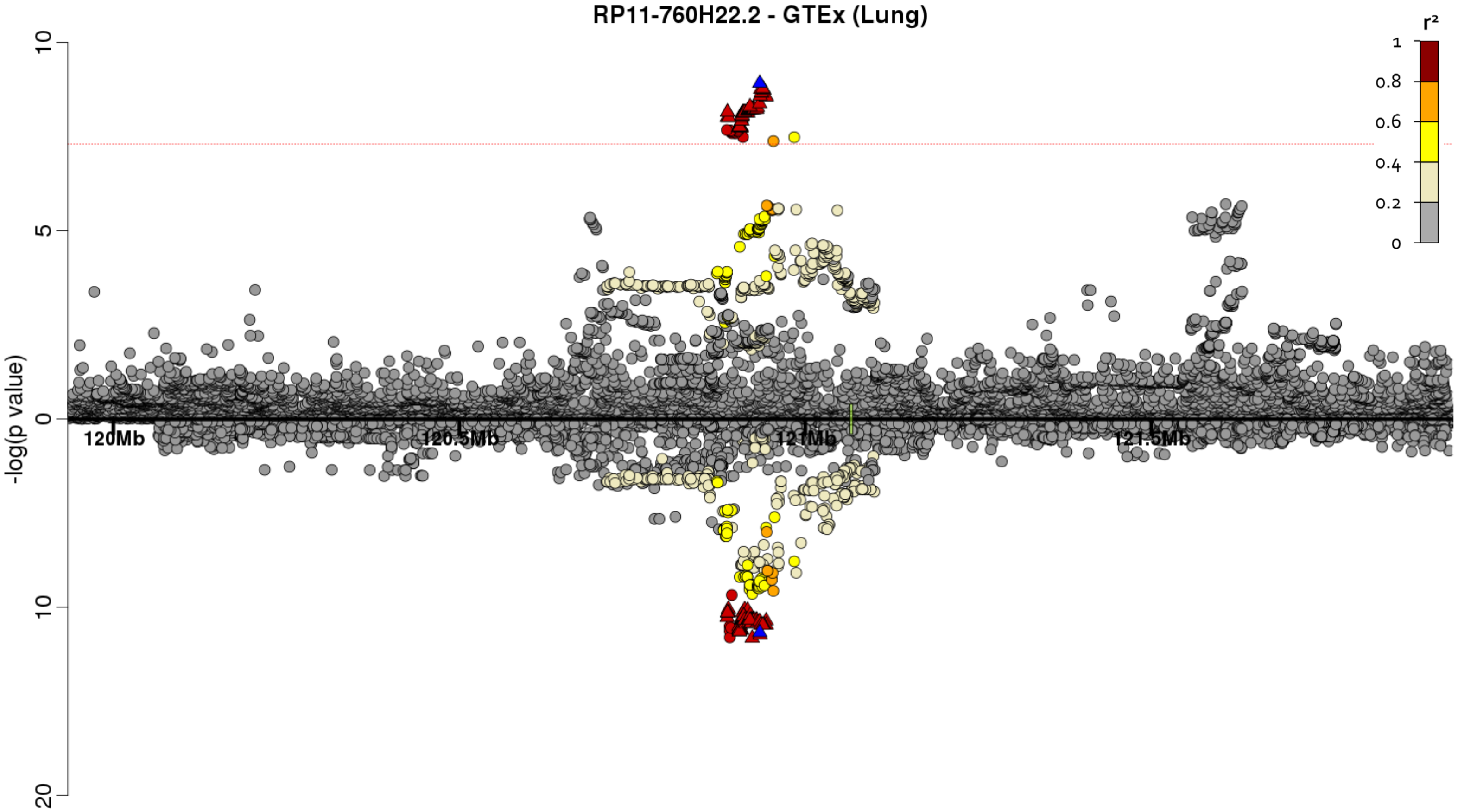

**xii) KB-1471A8.1 - Adipose (Subcutaneous) - Colocalisation probability = 85.6%**

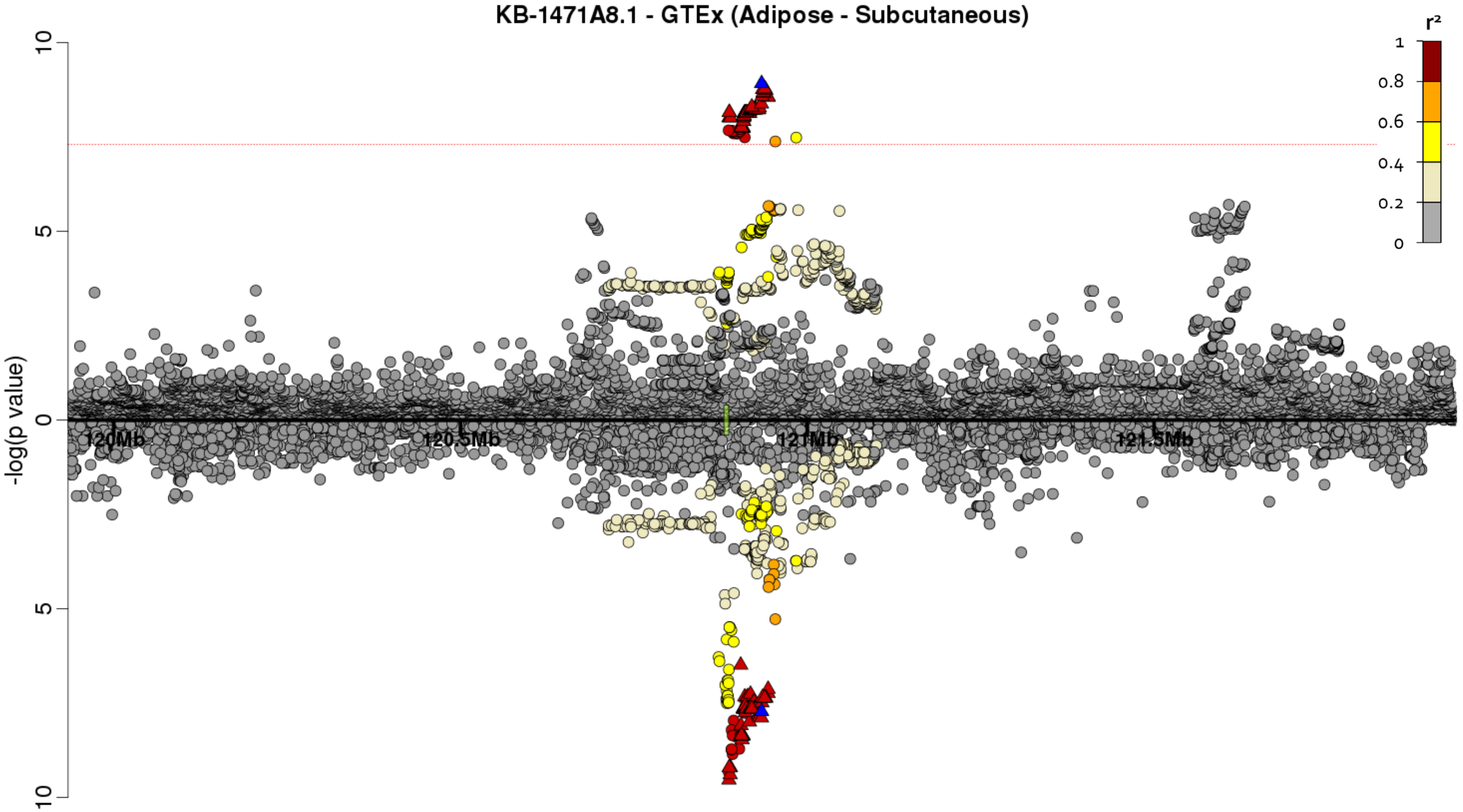

**xiii) KB-1471A8.1 - Adipose (Visceral) - Colocalisation probability = 90.9%**

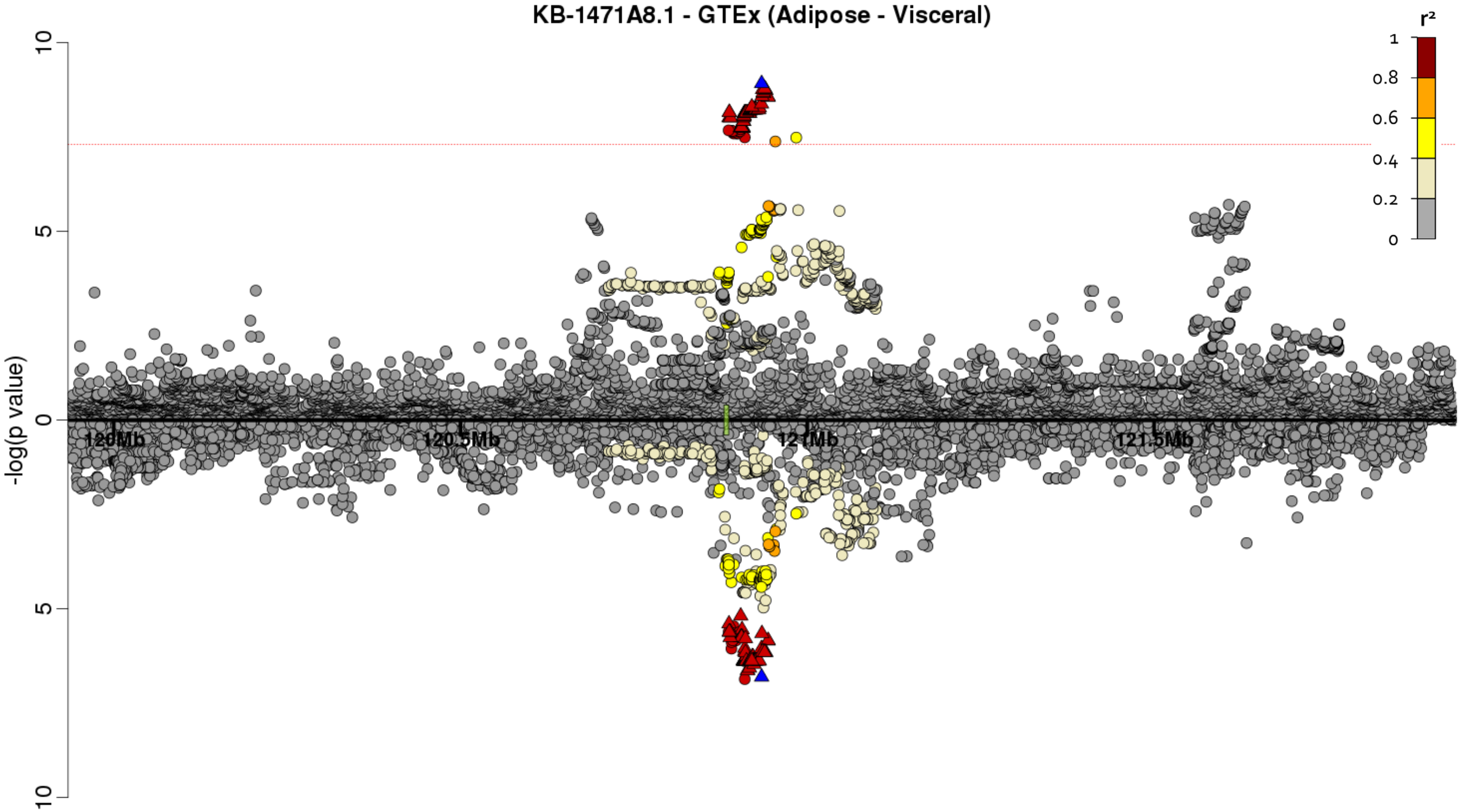

**xiv) KB-1471A8.1 - Skin (Sun exposed) - Colocalisation probability = 88.7%**

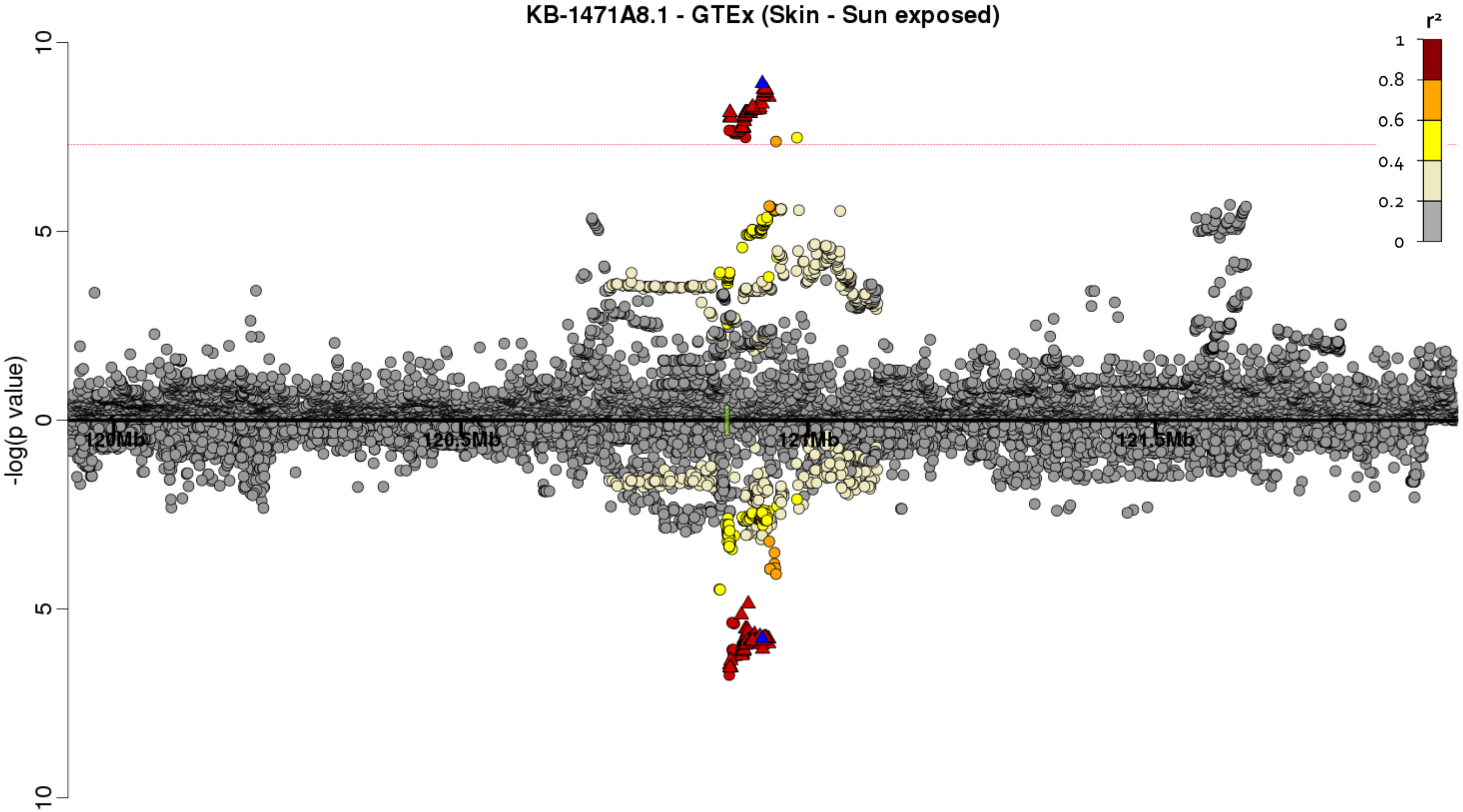

#### Supplementary Figure 10 - FORGE analysis for enrichment in regulatory regions

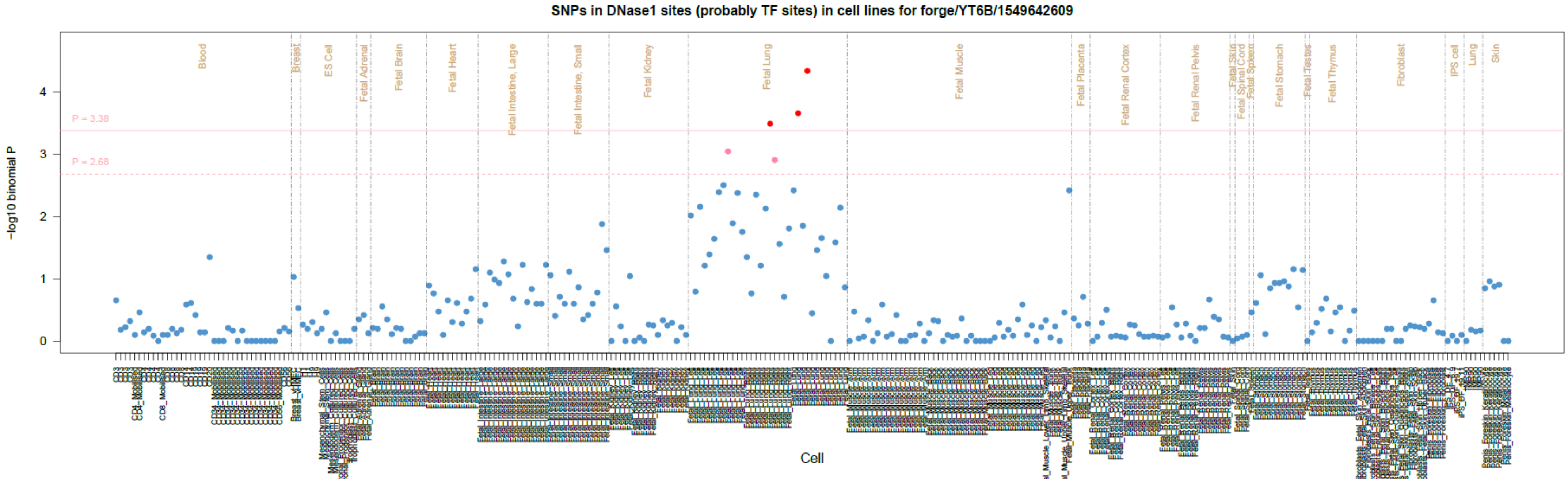

**Supplementary Figure 11 - GARFIELD analysis for enrichment in DNase I hypersensitivity sites by tissue**

Radial plots for enrichment. The distance from the centre is equal to the odds ratio with the peaks shown in black to be when using a *P* threshold in the IPF genome-wide analysis of $5 \times 10^{-8}$ and in blue for $5 \times 10^{-5}$. There are two rings of dots on the outside which show whether the site is significantly enriched after adjusting for multiple testing ($P < 3.59 \times 10^{-4}$). If there is a dot on the outermost ring then the site is significantly enriched when including all variants with $P < 5 \times 10^{-8}$ and if there is a dot on the inner ring then the site is enriched when including all variants with $P < 5 \times 10^{-5}$. If the site is significant for both thresholds there will be two dots.

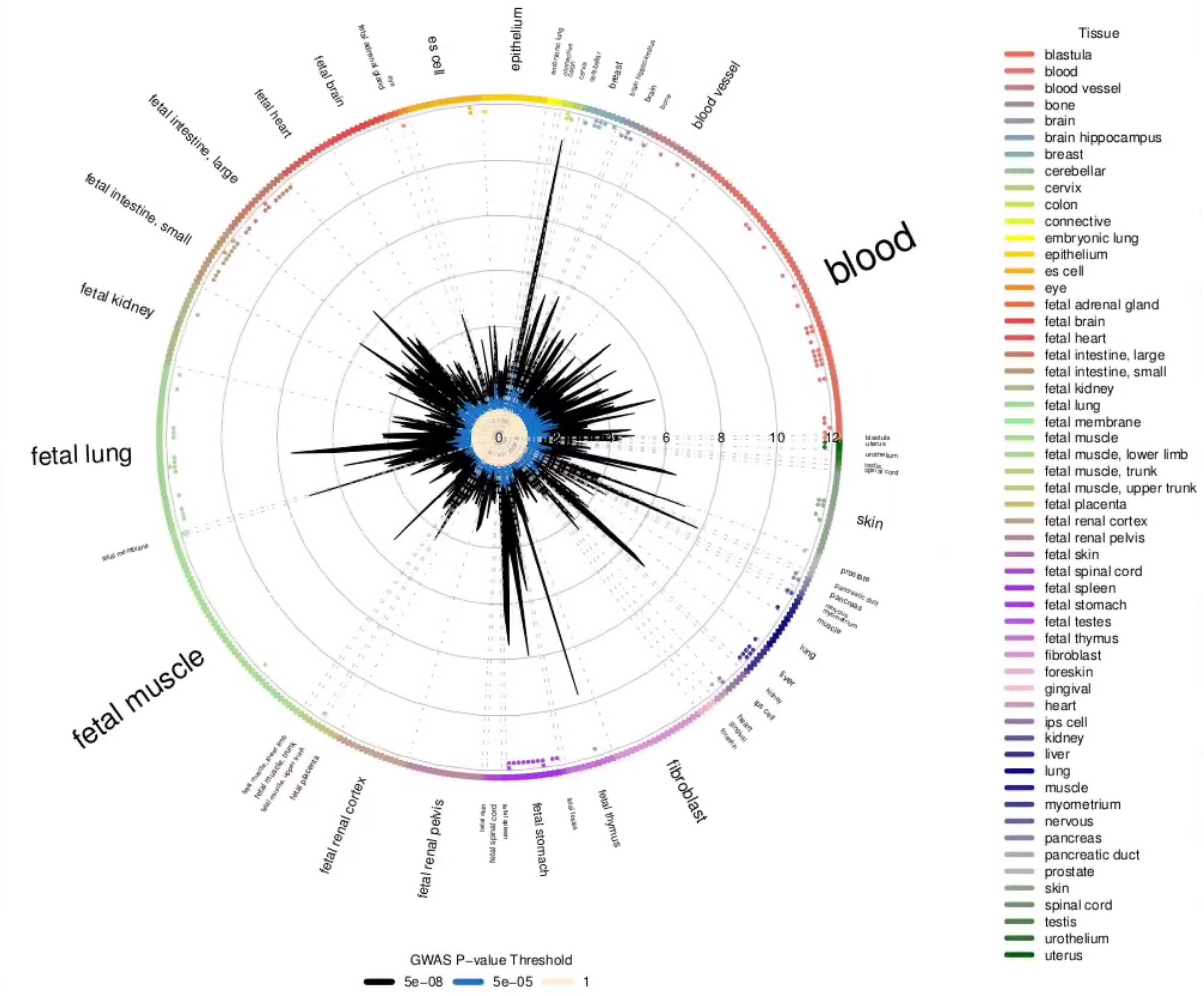

**Supplementary Figure 12 - Polygenic risk score in target dataset by *P* threshold used**

The x axis shows the *P* threshold ($P_T$) used for selecting variants to include in the risk score calculation. The black line and y axis on the left side shows the significance (−log(*P* value)) for the risk score for each $P_T$ tested. The red dotted line shows the threshold of 0.001 used for determining whether the risk score was significantly associated with IPF susceptibility. The orange line and y axis on the right side shows the model fit (Nagelkerke's $R^2$) of the risk score at each $P_T$ tested.

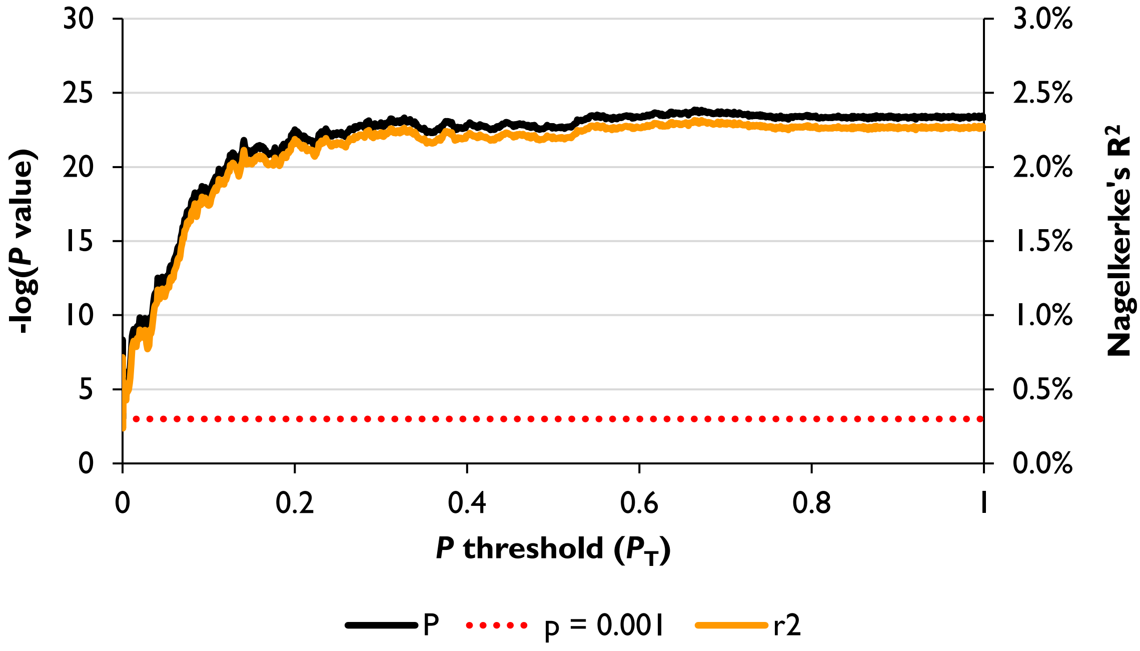
